# Optimizing and benchmarking polygenic risk scores with GWAS summary statistics

**DOI:** 10.1101/2022.10.26.513833

**Authors:** Zijie Zhao, Tim Gruenloh, Meiyi Yan, Yixuan Wu, Zhongxuan Sun, Jiacheng Miao, Yuchang Wu, Jie Song, Qiongshi Lu

**Affiliations:** Department of Biostatistics and Medical Informatics, University of Wisconsin-Madison, WI; Department of Statistics, University of Wisconsin-Madison, WI; Center for Demography of Health and Aging, University of Wisconsin-Madison, Madison, WI

**Author notes:** To whom correspondence should be addressed: Dr. Qiongshi Lu.

**Keywords:** genome-wide association study, GWAS summary statistics, polygenic risk score, PRS model-tuning, PRS benchmark, ensemble learning

## Abstract

**Background:** Polygenic risk score (PRS) is a major research topic in human genetics. However, a significant gap exists between PRS methodology and applications in practice due to often unavailable individual-level data for various PRS tasks including model fine-tuning, benchmarking, and ensemble learning.

**Results:** We introduce an innovative statistical framework to optimize and benchmark PRS models using summary statistics of genome-wide association studies. This framework builds upon our previous work and can fine-tune virtually all existing PRS models while accounting for linkage disequilibrium. In addition, we provide an ensemble learning strategy named PUMAS-ensemble to combine multiple PRS models into an ensemble score without requiring external data for model fitting. Through extensive simulations and analysis of many complex traits in the UK Biobank, we demonstrate that this approach closely approximates gold-standard analytical strategies based on external validation, and substantially outperforms state-of-the-art PRS methods.

**Conclusions:** Our method is a powerful and general modeling technique that can continue to combine the best-performing PRS methods out there through ensemble learning and could become an integral component for all future PRS applications.

## Background

Genetic risk prediction is a main focus in human genetics research and a key step towards precision medicine(1-3). Continued success in genome-wide association studies (GWAS) in the past decade has facilitated the development of polygenic risk scores (PRS) that aggregate the effects of millions of single nucleotide polymorphisms (SNPs) for many complex traits(4-6). Compared to earlier statistical methods that require individual-level data for model training(7-10), PRS which only relies on GWAS summary data is much more generally applicable due to the wide availability of GWAS summary statistics. Although earlier PRS models struggled to produce accurate prediction results, recent and more sophisticated PRS methods have achieved substantially improved prediction accuracy through statistical regularization and biological data integration(11-17). In numerous studies, PRS has shown promising performance in stratifying disease risk and great potential in informing early lifestyle changes or medical interventions(18-21).

Despite the progress, several lingering challenges create a significant gap between PRS methodology and applications. A main recurring issue we highlight (and address) throughout the paper is that PRS modelers often assume the existence of independent individual-level datasets that can be used for additional model tuning. But in practice, GWAS summary statistics are used for PRS model training, meaning that conventional sample splitting schemes cannot be used. Additional datasets that are independent from both training and testing samples also rarely exist. This suggests that model-tuning samples will have to come from the precious testing dataset which inevitably reduces the sample size and statistical power in downstream applications.

This disconnection between impractical method requirements and limited data availability can lead to a variety of problems. For example, many PRS methods have tuning parameters that could substantially swing model performance when not chosen properly(12-15, 22-24). Conventionally, these parameters need to be fine-tuned on a separate dataset with individual-level genotypes and phenotypes. Although some recent methods employ fully Bayesian or empirical Bayesian techniques to bypass model fine-tuning(25-27), these hyperparameter-free PRS do not always outperform fine-tuned models, trading predictive accuracy for computational feasibility(28, 29). Second, no PRS method universally outperforms all other approaches. The empirical performance of a PRS model depends on GWAS sample size, genetic architecture of the phenotype, quality of GWAS summary statistics, and heterogeneity between training and testing samples(30-33). Thus, it is of great interest to systematically and impartially benchmark various PRS methods for each trait, ideally in an independent dataset(11, 30, 34). Third, several recent studies have employed ensemble learning which combines multiple PRS models via another regression(28, 29) and showed improved PRS accuracy in both within- and cross-ancestry prediction applications(35-37). This brute-force approach has shown superior performance compared to any single PRS method but is data-demanding – the second level regression model needs to be fit on a separate dataset. Finally, we note that it may be of interest to combine all these tasks in practice, e.g., benchmarking an ensemble learner that combines multiple PRS models which all need to be tuned separately, which really seems like an impossible task.

In this paper, we seek a solution to these problems. We base our statistical framework on PUMAS, a method we recently introduced to perform Monte Carlo cross-validation (MCCV) using GWAS summary statistics(38). We have shown that PUMAS can effectively fine-tune PRS models with clumped SNPs(39) and similar approaches have since been adopted in other applications(40-42). Here, we first demonstrate that PUMAS can now fine-tune and benchmark state-of-the-art PRS models without SNP pruning. Second, we introduce an extension to the PUMAS framework named PUMAS-ensemble which is an innovative strategy to perform ensemble learning using GWAS summary data alone. Taken together, we showcase a sophisticated statistical framework for fine-tuning, benchmarking, and combining PRS models using GWAS summary statistics as input. We demonstrate the performance of our approach through extensive simulations and analysis of 21 complex traits in UK Biobank (UKB). On average, the PUMAS-ensemble ensemble PRS achieves a 8.93% relative gain in predictive R^2^ compared to LDpred2-auto and a 17.68% gain compared to PRS-CS-auto, respectively. We also apply our method to 31 well-powered GWAS with publicly available summary statistics and provide a catalog of ensemble PRS with benchmarked predictive performance.

## Results

### Method Overview

First, we present an overview of the PUMAS-ensemble workflow. Statistical details and technical discussions are presented in the **Methods** section. For illustration, first we assume individual-level data is available. In this case, we would divide the samples into 4 independent sets for PRS training, model fine-tuning, constructing ensemble PRS, and benchmarking model performance, respectively (**Figure 1A**). The main goal of our new approach is to mimic this procedure when only summary statistics are available. Using PUMAS, we could sample marginal association statistics for a subset of individuals in the GWAS(38). Doing this repeatedly, we could divide the full GWAS summary data to corresponding training, tuning, ensemble learning, and testing summary statistics (**Figure 1B**). Using these four sets of sub-sampled summary statistics, we train a series of PRS models, fine-tune each PRS model to select the besting tuning parameters, apply PUMAS-ensemble to combine PRS models through linear regression, and finally evaluate the predictive performance of PRS models. The entire procedure only requires GWAS summary statistics and linkage disequilibrium (LD) references as input.

**Figure 1.**
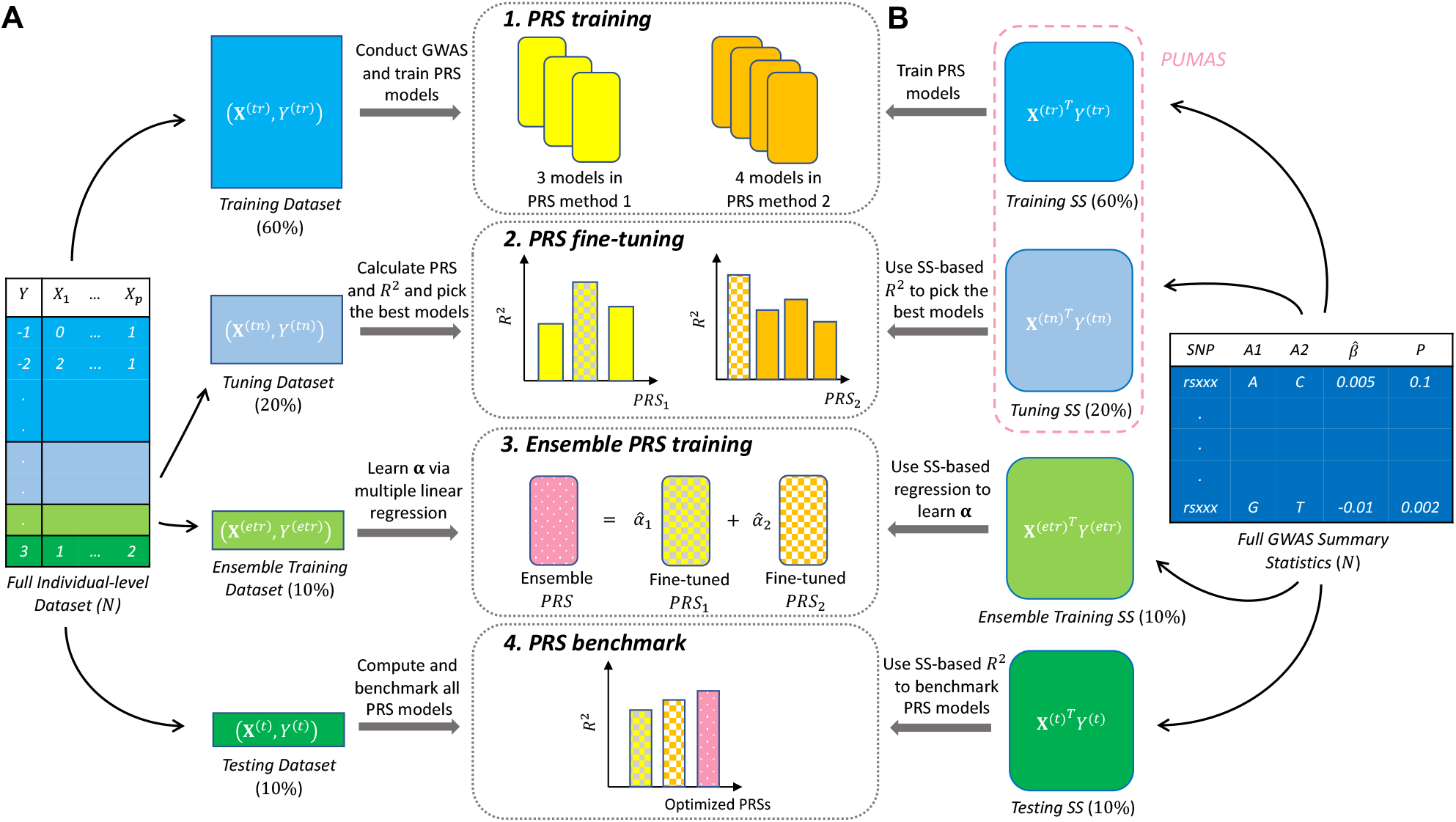
Workflow of PRS construction and evaluation. (**A**) Conventional approach divides the entire individual-level dataset to different subset of samples for each of 4 stages of PRS analysis. (**B**) PUMAS-ensemble directly partitions the full summary-level data to corresponding summary statistics for different analytical purposes.

### Simulation results

We performed simulations using imputed genotype data from UKB to demonstrate that PUMAS and PUMAS-ensemble can fine-tune, combine, and benchmark PRS models. We included 100,000 independent individuals of European descent and 944,547 HapMap3 SNPs in the analysis. We simulated phenotypes with heritability of 0.2, 0.5, and 0.8 and randomly assigned causal variants under sparse and polygenic settings to mimic different types of genetic architecture (**Methods**). We performed GWAS and obtained marginal association statistics. We then implemented PUMAS and PUMAS-ensemble to conduct a 4-fold MCCV to train, optimize, and evaluate lassosum, PRS-CS, LDpred2, and an ensemble PRS which combines all three methods and SDPR(22, 25, 26, 43). For comparison, we also implemented a MCCV procedure using individual-level UKB data. We partitioned the UKB dataset into 4 mutually exclusive datasets. We used datasets 1 and 2 to train and fine-tune each PRS method, then used the third dataset to fit a regression to combine multiple PRS. We evaluated each PRS method in the fourth dataset and reported PRS prediction accuracy quantified by *R*^2^. We describe implementation details of both summary-statistics-based and individual-level-data-based MCCV in **Methods**.

Overall, we observed highly consistent results between PUMAS/ PUMAS-ensemble and MCCV for both quantitative and binary phenotypes (**Figure 2**; **Additional file 1**: **Fig. S1**-**S7**; **Additional file 2, 3, 4**, and **5**: **Table. S1**-**S4**). In addition, summary statistics-based approaches can closely approximate *R*^2^ values obtained from model-tuning and benchmarking techniques using individual-level data. PUMAS-ensemble also constructed scores that were highly concordant with ensemble PRS built from individual-level data which universally outperformed all PRS models used as input. During the revision, we also added simulation analysis for MegaPRS(40) which also yields similar ensemble score performance (**Additional file 1**: **Fig. S8**-**S10**; **Additional file 6** and **7**: **Table. S5**-**S6**). Computation-wise, PUMAS’s subsampling step can execute in parallel for each chromosome and our benchmarking results suggest that PUMAS/PUMAS-ensemble are scalable for GWAS summary statistics including millions of SNPs (**Additional file 8**: **Table. S7**).

**Figure 2.**
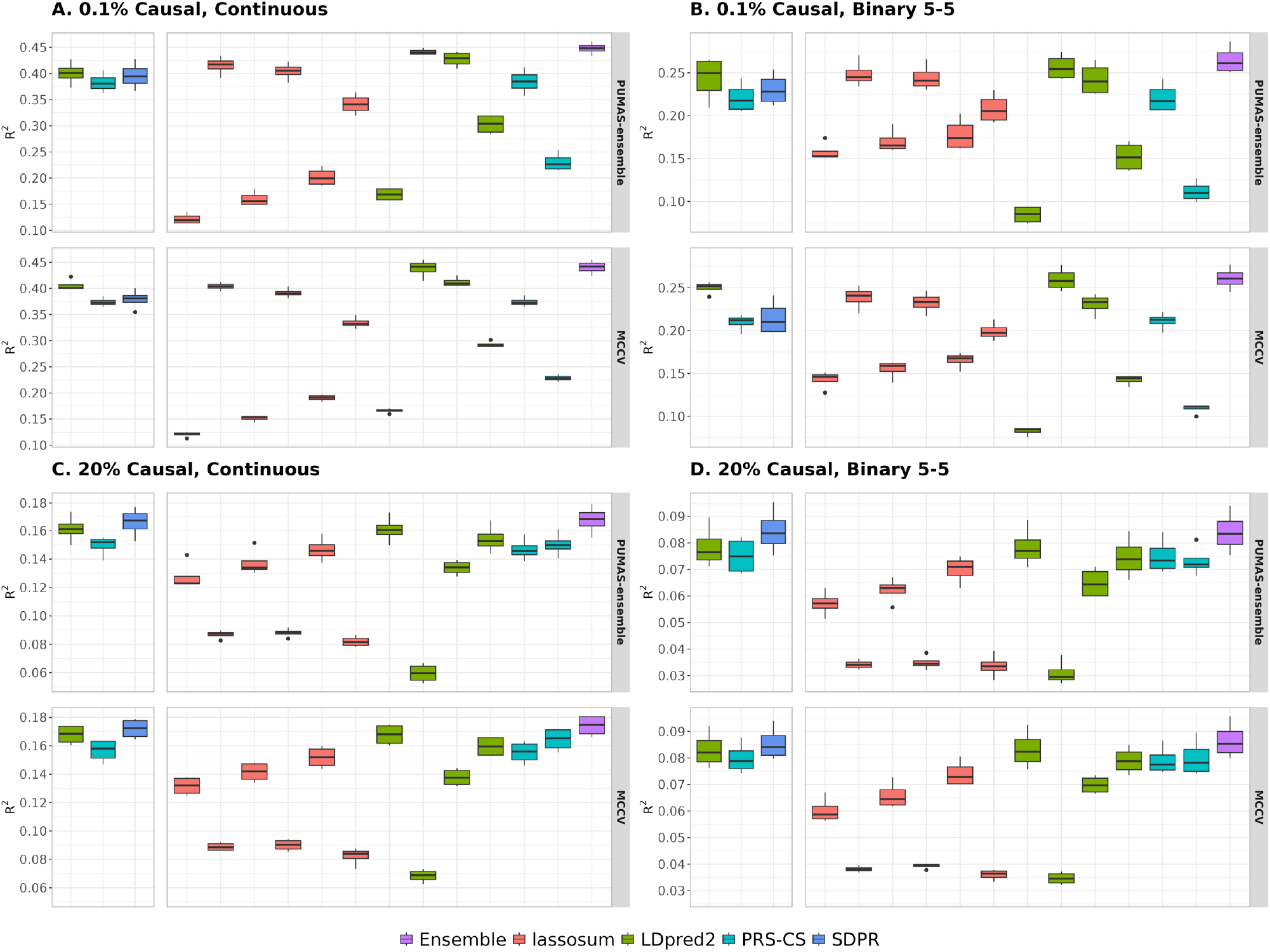
Comparison of PUMAS-ensemble and MCCV in UKB simulation. (**A** and **C**) Simulation results for quantitative traits. (**B** and **D**) Simulation results for binary traits with balanced case-control ratio. Proportion of causal variants is 0.1% in **A** and **B**, and 20% in **C** and **D**. The heritability is set to be 0.5 in all panels. Models that do not require fine-tuning are shown on the left side of each panel. Y-axis: predictive *R*^2^ across 4 repeats of MCCV; X-axis (left to right): tuning-free models: LDpred2-auto (green box), PRS-CS-auto (blue box), and SDPR (dark-blue box). lassosum models (red boxes) with tuning parameter settings: s=0.2 and λ=0.005, s=0.2 and λ=0.01, s=0.5 and λ=0.005, s=0.5 and λ=0.01, s=0.9 and λ=0.005, s=0.9 and λ=0.01. LDpred2 models (green boxes): non-infinitesimal with p=0.1, non-infinitesimal with p=0.01, non-infinitesimal with p=0.001, and infinitesimal model. PRS-CS (blue boxes): ϕ=0.01 and 0.0001. Finally, the purple box shows the results of ensemble PRS. Results for remaining simulation settings are summarized in **Additional file 1**: **Fig. S1**-**S10** and **Additional file 2, 3, 4, 5, 6**, and **7**: **Table. S1**-**S6**.

We conducted two additional analyses to demonstrate the validity of PUMAS subsampling framework. First, we benchmarked PUMAS against the subsampling approach implemented in MegaPRS(40), which uses ‘pseudo summary statistics’ for parameter tuning, and the gold standard approach MCCV based on individual-level data (**Methods**). We found consistent subsampling results from both approaches compared to MCCV (**Additional file 1**: **Fig. S11**; **Additional file 9**: **Table. S8**). In addition, while PUMAS’s subsampling framework assumes weak individual SNP effects, we observed robust performance of PUMAS under extremely sparse genetic architecture with large SNP effects (**Additional file 1**: **Fig. S12**; **Additional file 10**: **Table. S9**). These results highlight the robustness of PUMAS’s summary statistics subsampling scheme under different genetic architecture.

### PUMAS can fine-tune and benchmark PRS methods

Next, we demonstrate that PUMAS effectively fine-tunes PRS models and performs accordantly with the gold standard external validation approach based on individual-level data. We applied PUMAS to 16 quantitative traits, 4 diseases, and 1 ordinal trait in UKB(44) (**Additional file 11** and **12**: **Table. S10**-**S11**). After quality control, the UKB dataset contained 375,064 independent individuals and 1,030,187 SNPs (**Methods**). We applied a 9-to-1 data split to hold out 10% of the samples for external validation, and performed GWAS for all traits using 90% of the samples. We applied 4-fold MCCV implemented in PUMAS to train and fine-tune three PRS models (i.e., LDpred2, lassosum, and PRS-CS which have been demonstrated to achieve high prediction accuracy in a recent benchmark study(22, 25, 26, 29)) using only summary statistics. For external validation, we trained PRS models using the full summary statistics and calculated PRS prediction accuracy on the holdout dataset. We report the best tuning parameters for LDpred2, lassosum, and PRS-CS and corresponding *R*^2^ obtained from both PUMAS and external validation.

Our summary-statistics-based approach showed highly consistent model-tuning performance for all analyzed traits compared to external validation (**Figure 3, Additional file 1**: **Fig. S13**-**S33**; **Additional file 13** and **14**: **Table. S12**-**S13**). Among 21 traits, PUMAS and external validation selected the same best tuning parameters 21, 18, and 11 times for lassosum, LDpred2, and PRS-CS, respectively. When the model tuning results were different between PUMAS and external validation, both approaches still selected models with very similar prediction accuracy. Indeed, PUMAS provided precise R^2^ estimates for all models compared to external validation, advocating the use of our summary-statistics-based approach for PRS model benchmarking. In addition, it is noteworthy that empirical and full Bayesian approaches (i.e., LDpred2-auto and PRS-CS-auto) did not always outperform other fine-tuned PRS models even within the UKB cohort, demonstrating the necessity of PRS model tuning for optimizing out-of-sample prediction.

**Figure 3.**
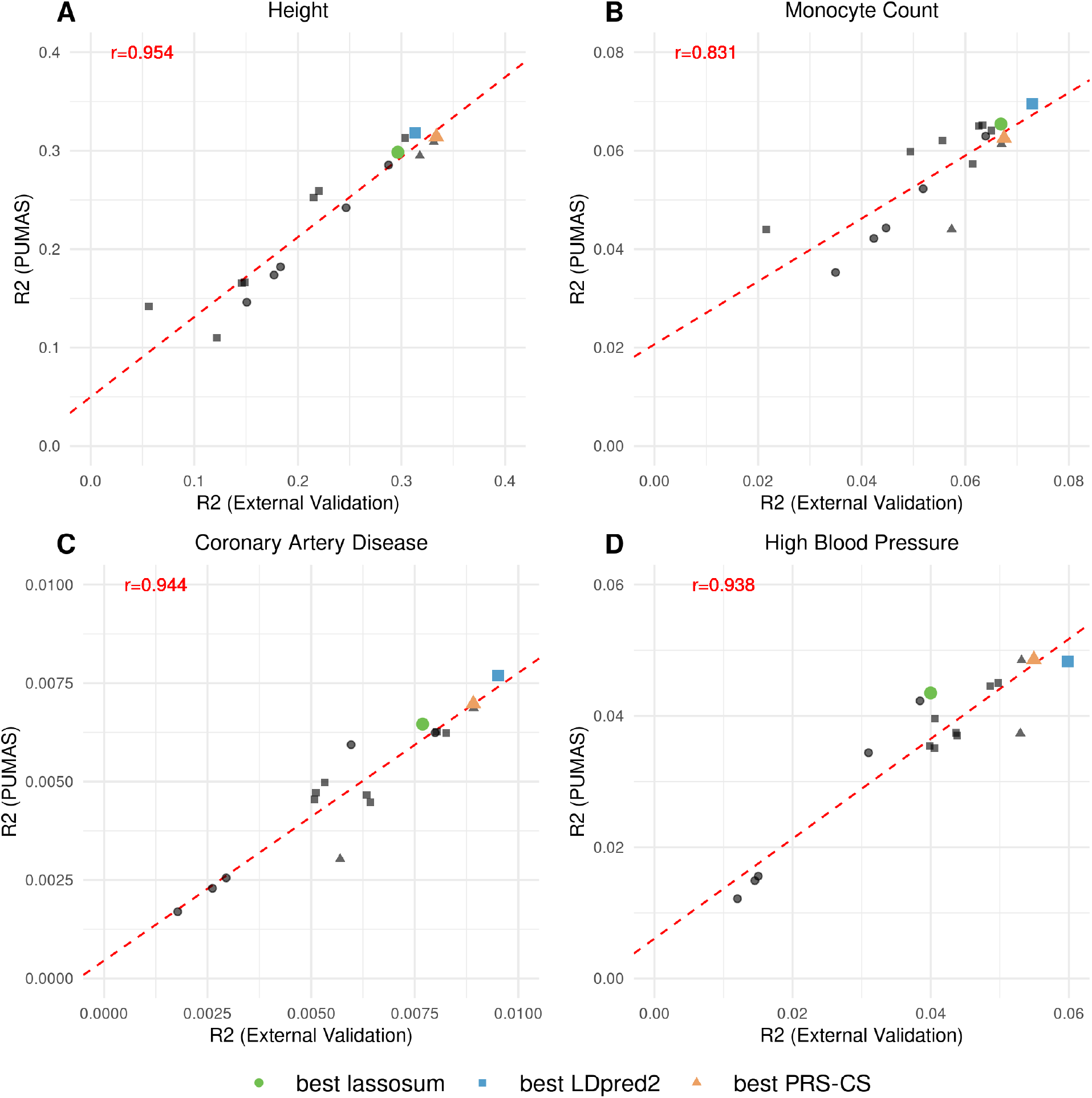
Comparing PUMAS results with external validation. Four panels show the model-tuning results for (**A**) height, (**B**) monocyte count, (**C**) coronary artery disease, and (**D**) high blood pressure. Y-axis: average predictive *R*^2^ across 4-fold replications from PUMAS; X-axis: predictive *R*^2^ evaluated by external validation on the holdout dataset. Each data points represents a PRS model with different tuning parameters and the shape of data points indicate three different PRS methods: LDpred2, PRS-CS, and lassosum. The best tuning parameter setting suggested by PUMAS for each PRS method is highlighted and colored. The dashed red line is fitted regression line between PRS *R*^2^ from PUMAS and external validation. Pearson correlations between two sets of results are shown in each panel. Detailed model-tuning results for all 21 traits are summarized in **Additional file 1**: **Fig. S13**-**S33** and **Additional file 13** and **14**: **Table. S12**-**S13**.

We also observed that the parameter-tuning results are accordant with the analyzed traits’ genetic architecture. For both height and monocyte count, PUMAS accurately selected the best tuning parameters based on external validation (**Figure 3A-B**), but the selected models were not the same between these two traits. Height is known to be extremely polygenic with more than 12,000 independent GWAS signals in the latest GWAS(45). In comparison, fewer loci have been found to significantly associate with monocyte count(46). Our model-tuning results suggest that polygenic prediction models fit best for height (e.g., LDpred2-Infinitesimal and PRS-CS with *ϕ* = 0.01) while sparser PRS models with stronger regularization (e.g., PRS-CS with *ϕ* = 0.0001) provide better prediction accuracy for monocyte count.

Finally, PUMAS can also effectively estimate predictive *R*^2^ for binary traits (**Figure 3C-D**). To calculate interpretable *R*^2^ for binary outcomes, PUMAS first transforms GWAS summary statistics obtained from logistic regressions to the linear regression scale, and then computes *R*^2^ on the observed scale(47-49). To show that such transformation is valid, we trained two sets of PRS models using both transformed and original logistic regression summary statistics for 4 disease traits and observed nearly identical PRS performance between two approaches (**Additional file 1**: **Fig. S29**-**S32**; **Additional file 14**: **Table. S13**). Details in the implementation of binary trait analysis and summary statistics transformation are presented in **Methods**.

### Ensemble learning via PUMAS-ensemble substantially improves PRS prediction accuracy

Here we apply PUMAS-ensemble, the ensemble learning extension of PUMAS, to UKB traits and show that ensemble PRS has superior prediction accuracy compared to each PRS method and our summary statistics-based approach is comparable to ensemble learning results based on individual-level data. We constructed linearly combined scores of lassosum, PRS-CS, LDpred2, and SDPR. Using individual-level data, we split the 10% UKB holdout dataset into two equally sized subsets. We fitted a multiple regression on the first holdout set to aggregate the best-performing PRS models trained and tuned from GWAS summary statistics, and then evaluated the ensemble score’s prediction accuracy using the second holdout set. For comparison, we implemented PUMAS-ensemble to conduct 4-fold MCCV to perform ensemble learning using summary statistics alone and assess its performance on the second holdout set.

Our approach showed almost identical performance compared to individual-level data results (**Figure 4A**), showcasing PUMAS-ensemble’s ability to benchmark and construct ensemble PRS without requiring additional datasets. In addition, ensemble PRS achieved the highest prediction accuracy compared with four input PRS models for all traits except diastolic blood pressure (**Additional file 1**: **Fig. S34**; **Additional file 15**: **Table. S14**). The ensemble PRS using individual-level data as input had an average 19.67% and 10.76% relative gain in *R*^2^ compared to PRS-CS-auto and LDpred2-auto while the PUMAS-ensemble ensemble PRS delivered a similar 17.68% and 8.93% *R*^2^ increase respectively (**Figure 4B**), highlighting the substantial gain in prediction accuracy from ensemble learning.

**Figure 4.**
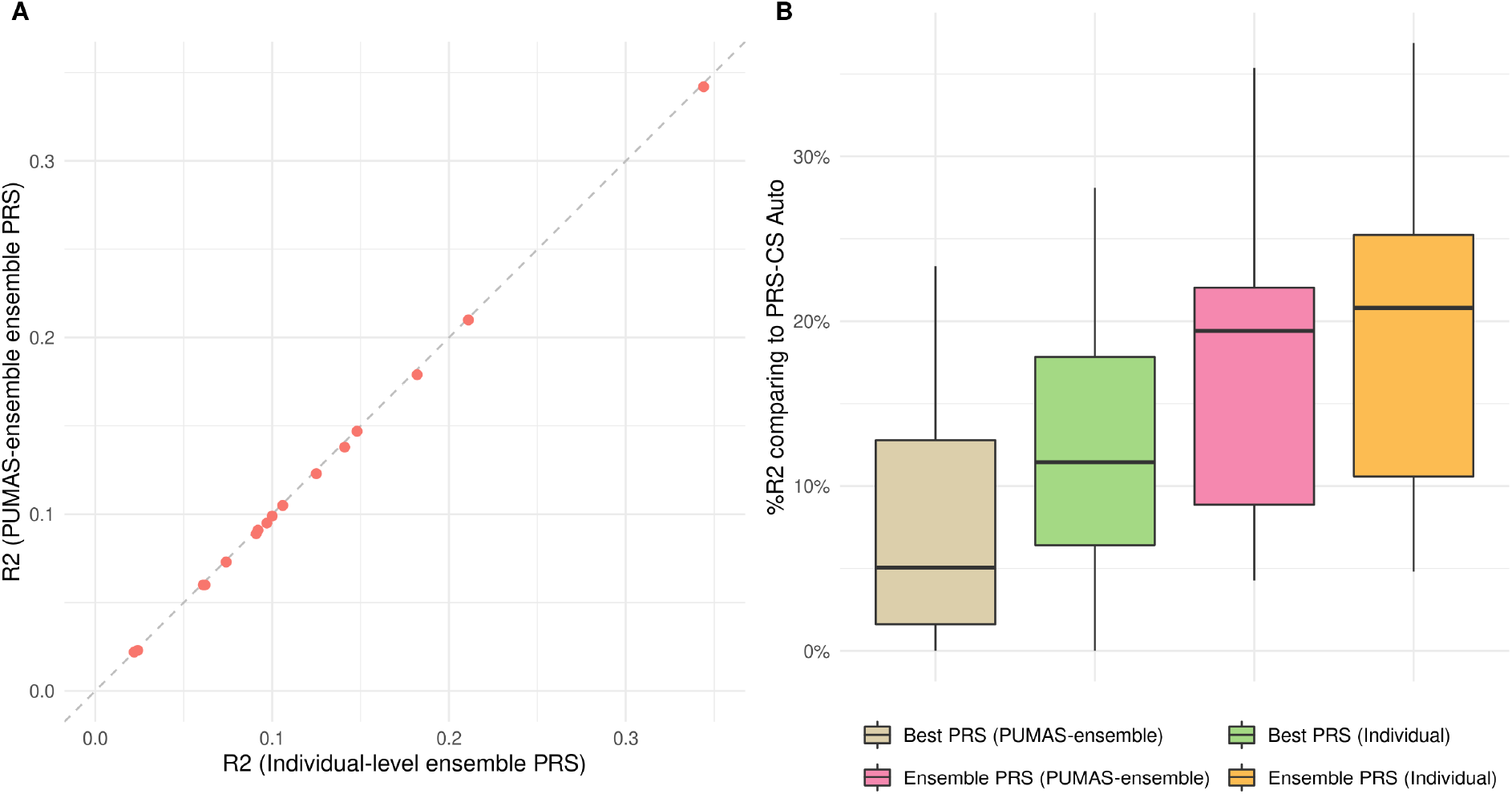
Constructing ensemble PRS for UKB traits. (**A**) Comparing two sets of ensemble PRS obtained from PUMAS-ensemble and individual-level data. The gray dashed line is the diagonal line. (**B**) Comparing ensemble PRS with input PRS methods. Y-axis: relative percentage increase in *R*^2^ compared to PRS-CS-auto; X-axis: 4 sets of PRS models, including the best single PRS suggested by PUMAS, the best single PRS selected based on the first individua-level holdout set, the ensemble PRS obtained from PUMAS-ensemble, and the ensemble PRS trained from individual-level data. All *R*^2^ values were computed using the second half of holdout dataset.

**Figure 5.**
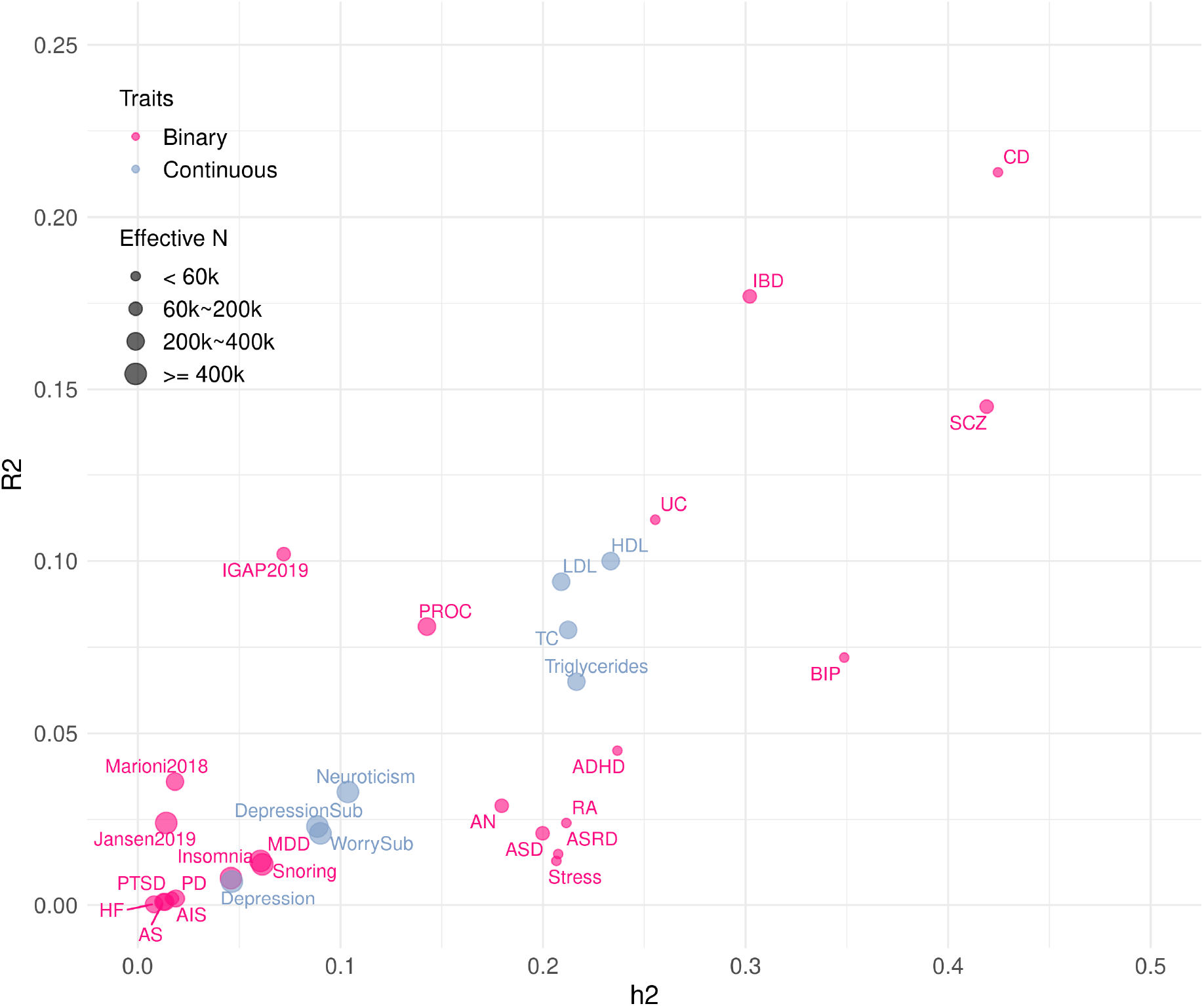
An ensemble PRS catalog for 31 complex traits. Y-axis: average predictive *R*^2^ of PUMAS-ensemble ensemble PRS; X-axis: heritability estimates from LD score regression(50). Size of data points indicates the effective sample size of each GWAS. Binary traits and continuous traits are highlighted with different colors. Detailed PRS benchmark results are presented in **Additional file 20**: **Table. S19**.

We sought to explore some properties of PUMAS-ensemble in real-world settings. A comparison between PUMAS and PUMAS-ensemble suggests that although ensemble learning requires additional splitting of data and reduces training sample size for individual PRS models, ensemble scores trained on smaller GWAS training subsets outperforms individual fine-tuned PRS, highlighting the benefit of ensemble learning (**Methods**; **Additional file 16**: **Table. S15**). Furthermore, we conducted sensitivity analyses to investigate the effect of LD misspecification on PUMAS-ensemble (**Methods**). We observed reduced ensemble PRS performance from PUMAS-ensemble when the LD reference data mismatches the ancestral population of GWAS samples (**Additional file 1**: **Fig. S35**-**S36**; **Additional file 17**: **Table. S16**).

### Constructing and benchmarking ensemble PRS for 31 complex traits

Finally, we applied PUMAS-ensemble to provide a comprehensive catalog of ensemble PRS for 31 publicly available GWAS summary statistics with varying sample size and genetic architecture. The detailed information and selecting criteria for GWAS summary-level data are summarized in **Methods** and **Additional file 18**: **Table. S17**. We employed extensive quality controls to pinpoint and calibrate misspecifications in GWAS summary statistics following a recent study(31) (**Additional file 19**: **Table. S18**). We also transformed logistic summary statistics to linear scale to produce interpretable *R*^2^ for binary traits(47-49). For each trait, we reported prediction accuracy of the best performing PRS model and ensemble PRS. The full results of the PRS catalog are presented in **Additional file 20**: **Table. S19**. The predictive performance of ensemble PRS is correlated with estimated trait heritability, and the predictive *R*^2^ ranged from 3E-4 to 0.213 across 31 traits, showing highly diverse predictive performance of genetic risk prediction. We also note that ensemble PRS improved predictive *R*^2^ for every trait in the analysis with an median increase of 26.36% compared to PRS-CS-auto (**Additional file 1**: **Fig. S37**). Among 31 complex diseases and traits, we observed the highest prediction improvement for rheumatoid arthritis (113.7%), Alzheimer’s disease (93.0%, 94.3%, and 109.5% on three datasets), and post-traumatic stress disorder (70.7%).

Another observation is that the ensemble PRS *R*^2^ exceeded the estimated trait heritability for all three Alzheimer’s disease GWAS. To demonstrate that this is not an artifact from overestimating predictive *R*^2^, we conducted additional analysis (**Methods**) using IGAP 2019 Alzheimer’s GWAS summary statistics(51) and compared our results with external validation based on 2,600 Alzheimer’s disease cases and 5,200 healthy controls in UKB (**Additional file 21**: **Table. S20**). The *R*^2^ of AD PRS obtained from external validation also exceeded estimated heritability (*h*^2^=0.072, SE=0.012) and the results were consistent with PUMAS *R*^2^ estimation (**Additional file 1**: **Fig. S38**; **Additional file 22**: **Table. S21**). We hypothesized that this is driven by the *APOE* region which contributes an unusually large fraction of AD risk(52-54). Indeed, after removing 383 SNPs in the *APOE* region from IGAP 2019 AD summary statistics (**Methods**), we observed a steep decline in *R*^2^ for both external validation and PUMAS. Both *R*^2^ values became substantially lower than the estimated *h*^2^ of 0.066 without *APOE* region (SE=0.009; **Additional file 22**: **Table. S21**).

## Discussion

Fine-tuning and benchmarking PRS models are challenging tasks due to the need of external individual-level datasets that are independent from the input GWAS. In this work, we extended our PUMAS approach to incorporate LD and fine-tune state-of-the-art PRS methods. In both simulations and analysis of UKB traits, we observed high concordance between PUMAS and results based on external validation using holdout samples. In addition, we presented a novel framework named PUMAS-ensemble to perform ensemble learning and create combined PRS using only GWAS summary statistics. We showed that ensemble PRS created by PUMAS-ensemble closely approximates scores built from holdout samples. Further, these ensemble scores substantially outperformed state-of-the-art PRS methods for complex traits we analyzed in the study. Finally, we applied PUMAS-ensemble to a collection of publicly available GWAS summary statistics and provided a comprehensive catalog of benchmarked and optimized PRS.

Our work presents several major advances that will impact future PRS applications. First, our method fills an important gap between PRS methodological research and its real-world applications. Currently, many PRS methods still have tuning parameters and grid search on external individual-level datasets remains the most common technique for fine-tuning these models. In practice, this kind of data can either be impossible to obtain, or need to be split from testing samples which could hurt statistical power in PRS applications(32). Our method provides a universal solution to PRS model fine-tuning. It is also noteworthy that some recent PRS methods such as MegaPRS(40) can also conduct model fine-tuning. MegaPRS bases its framework on GWAS z-scores and uses the LD matrix for summary statistics subsampling. On the other hand, PUMAS uses unstandardized SNP effect estimates and standard errors as inputs, and also considers GWAS regression residual variances in addition to LD for summary statistics partitioning. In practice, directly modeling GWAS standard errors and regression residual variances can be crucial when handling meta-analytic GWAS summary statistics(31) and when the trait of interest has a sparse genetic architecture. Second, model benchmarking is another major challenge in the field which conventionally relies on external validation data. Comprehensive and unbiased benchmarking allows researchers to compare the effectiveness of different PRS methods for particular traits of interest, and importantly, estimate PRS predictive accuracy without using testing samples. We note that although some advanced PRS approaches do not require model fine-tuning anymore, no existing methods could benchmark model performance using a single set of GWAS summary data, which is crucial for model selection, power calculation, and study design. Our approach now provides a solution to this problem. Third, the ensemble learning approach which combines multiple predictive models through a second level regression has been viewed as a highly effective but data-demanding approach(28, 29, 33). A major advance in this study is the introduction of PUMAS-ensemble which allows ensemble learning on GWAS summary statistics. We note that this approach not only showcased a substantial gain over existing PRS methods, but is generally applicable to future PRS developments. If a future PRS approach shows promising improvements compared to older methods, that new approach can also be incorporated into the ensemble PRS. In our view, PUMAS-ensemble is not a competing approach for any existing PRS model, but instead is a flexible and general modeling technique that combines the best-performing methods out there and should be applied to all future PRS applications. It is important to note that our implemented software allows users to specify how they wish to split the GWAS summary statistics into subsets. In practice, we recommend that researchers customize data partitioning of summary statistics tailored towards their analytical needs.

Our study has several limitations. First, we have constrained most statistical analysis in this study to the European ancestral population. PRS is known to transfer poorly in terms of prediction accuracy for non-European populations which could exacerbate the disparity in genomic medicine between ancestral groups(55, 56). While we conducted sensitivity analysis to demonstrate less robust ensemble model training due to LD misspecification, it is an important future direction to systematically optimize and benchmark PRS for diverse ancestral populations which would require incorporation of multiple sets of ancestry-specific GWAS and LD references. Although we did not extensively explore this topic in this paper, our recent work introduced parallel ideas to tackle the challenges in multi-ancestry genetic risk prediction(42). Second, we did not investigate the effect of assortative mating on PUMAS and PUMAS-ensemble in this study. Assortative mating is known to affect LD structure in human genome, bias heritability estimation(57), and affect PRS accuracy(58). The extent to which assortative mating influences our results requires further investigation. Third, analyses in this study were limited to GWAS summary statistics computed from independent samples. It remains to be investigated whether application of these approaches will be affected if the input GWAS summary statistics were obtained from linear mixed models with related samples or family-based designs(59-61). Future work will focus on developing statistical methods to correct for sample relatedness or demonstrate robustness to these issues. That said, we expect PRS model-tuning to remain valid even with sample relatedness since the inflation in *R*^2^ should be uniform across various tuning parameter settings, although biases may be introduced to the predictive *R*^2^ which could affect benchmarking efforts. Fourth, PUMAS/PUMAS-ensemble uses *R*^2^ on the observed scale(49) to evaluate PRS accuracy for binary traits but AUC is adopted for classification more frequently. Although we have shown in an earlier work(38) that AUC and *R*^2^ demonstrated highly consistent performance for PRS model fine-tuning, it remains future work to incorporate summary-statistics-based AUC estimator(62) into the PUMAS framework. Furthermore, our current analyses focused only on PRS derived from lassosum, PRS-CS, LDpred2, SDPR, and MegaPRS based on HapMap3 SNPs. While it serves to support the superiority of ensemble PRS as a proof of concept, more genetic variants and more PRS methods need to be jointly modeled and evaluated in the future, including scores that leverage auxiliary information including functional annotation(13, 14) or multiple phenotypes(15, 17, 63). Particularly for multi-trait PRS models, extending PUMAS to conduct multi-GWAS subsampling while modeling sample overlap between these GWAS summary statistics may be necessary. Finally, collinearity among PRS models could arise when using multiple regression to combine a large number of scores since some PRS methods tend to yield similar results.

Therefore, another future direction is to incorporate variable selection strategies into our ensemble learning framework, including penalized regression that has been employed in ensemble models based on individual-level data(35-37). It also remains an interesting but challenging task to fit non-linear ensemble PRS models using only GWAS summary statistics for incorporating machine learning ensemble methods such as XGBoost(64) used in Multi-PGS(63).

## Conclusions

We presented a sophisticated statistical framework to fine-tune, combine, and benchmark PRS methods using only GWAS summary statistics. This is a statistically novel and computationally efficient approach with flexible implementation that can handle a variety of applications. We have demonstrated its performance through careful and comprehensive analyses, and we argue that this framework presents highly innovative and generally applicable features that should become the default in many future PRS studies.

## Methods

### Sampling distribution of summary statistics

We adopt a commonly used linear model framework to quantify the relationship between a quantitative trait and SNP genotypes:

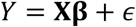

Here, *Y* denotes the trait, **X** = (**X**_1_, …, **X**_*p*_) denotes the genotypes of *p* SNPs, **β** ∈ ℝ^*p*^ denotes their true effect sizes, and *ϵ* denotes the random error that is independent from **X** and follows a normal distribution with mean zero and some variance 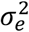. Let **y** and **x** = (**x**_1_, …, **x**_*p*_) denote the observed values for *Y* and **X** from *N* independent individuals. For simplicity, we assume both **y** and **x**_*j*_ (*j* = 1, …, *p*) are centered. Then, GWAS summary statistics can be denoted as:

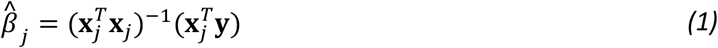

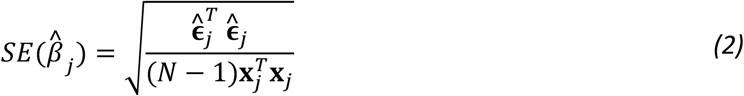

where 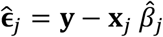 are the residuals from the marginal linear regression between the trait and the *j*-th SNP. To train, fine-tune, combine, and benchmark PRS models, independent datasets are required to avoid overfitting. We have previously proposed a flexible statistical framework to generate training and fine-tuning datasets when only GWAS summary statistics are available(38). Here, we generalize this statistical framework in two different directions. First, we allow our method to incorporate LD information. We note that this extension is similar to some recent work built on our initial PUMAS paper(40, 42). Second, we allow the method to partition full GWAS summary statistics into more than two datasets for various analytical purposes. Let *y*^*(s)*^ and *x*^*(s)*^ denote phenotype and genotype data for any arbitrary subset of *N* individuals with sample size *N*^*(s)*^. When *N* is large enough, we have previously shown that by central limit theorem(38):

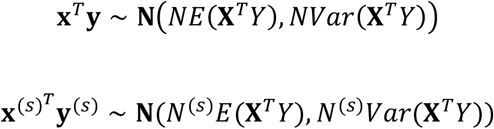

where **X**^*T*^*Y* = (*X*_1_*Y*, …, *X*_*p*_*Y*)^*T*^. Then, given the observed summary-level data from GWAS, the conditional distribution of summary statistics of a subset of GWAS samples is

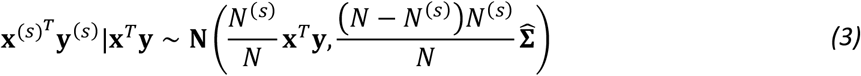

where 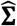 is the observed variance-covariance matrix for **X**^*T*^*Y*. To subsample summary statistics 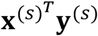, we need to estimate **x**^*T*^**y** and 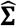 first. Recall formula *(1)* for marginal regression coefficient estimation, 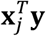 can be calculated using 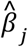 and 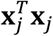 which is proportional to SNP variance and can be estimated by minor allele frequency (MAF) reported from GWAS or imputed from LD reference panel. On the other hand, deriving **Σ** is more complicated and we discuss how 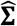 is estimated using summary statistics and an LD reference panel in the following section.

### Estimate variance-covariance matrix of summary statistics

Let *D* denote the SNP correlation matrix and *d*_*jk*_ denote the correlation between the *j*-th and the *k*-th SNPs. Let **Σ** be the true covariance matrix of summary statistics with diagonal and off-diagonal elements denoted as **Σ**_*j*_ and **Σ**_*jk*_, respectively. For convenience, we write *Y* = **Xβ** + *ϵ* = *X*_1_*β*_1_+… +*X*_*p*_*β*_*p*_ + *ϵ* = *X*_*j*_*β*_*j*_ + *ϵ*_*j*_, where *ϵ*_*j*_ = Σ_*i:i* ≠ *j*_ *X*_*i*_*β*_*i*_ + *ϵ*. Then the diagonal terms of the **Σ** can be written as

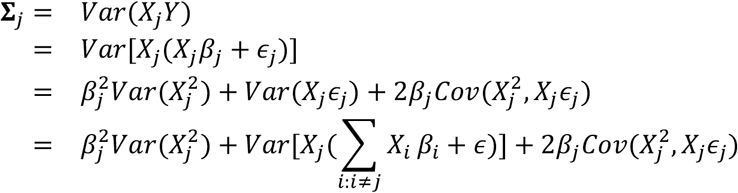

We partition all SNPs in the genome into 2 sets. Let *S*_1_ be the index set that contains all SNPs that are independent from the *j*-th SNP and *S*_2_ be the set with all remaining SNPs that are in LD with the *j*-th SNP. Then we can further expand **Σ**_*j*_ by

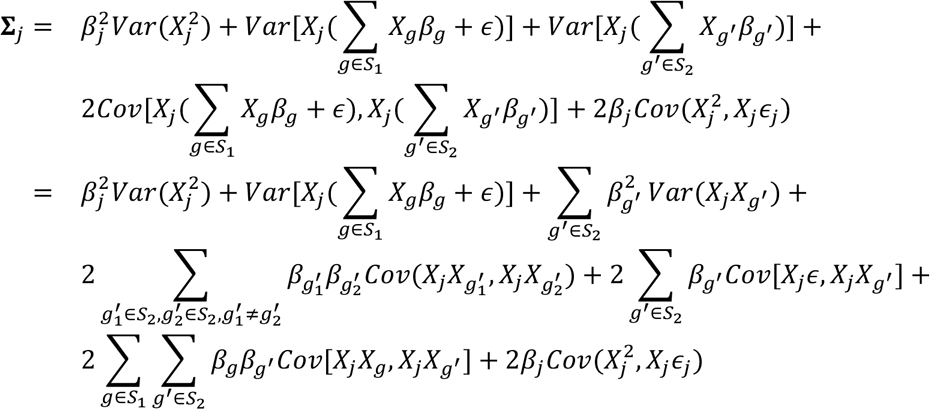

We can simplify **Σ**_*j*_ based on two commonly made assumptions. First, any given SNP should be in linkage equilibrium with the vast majority of SNPs in the genome. Therefore, we can safely assert ∣*S*_1_∣ ≫ ∣*S*_2_∣. Second, each individual SNP’s effect on the phenotype is typically very small such that the products of any effect sizes are negligible in practice. Taken together, we can reduce the expansion of **Σ**_*j*_ by discarding SNPs in *S*_2_ which eventually allows us to treat *X*_*j*_ and *ϵ*_*j*_ as independent in practice:

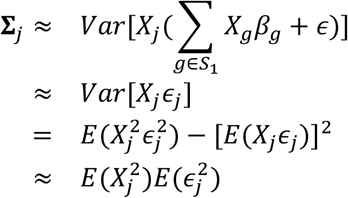

Note that 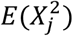 can be easily approximated using an MAF-based estimator, denoted as 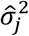, that may be obtained either from the full GWAS summary statistics or the LD reference data. For 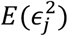, we can estimate its value by standard error of effect size estimation from GWAS summary data using formula *(2)*. In this way we can obtain an estimator of **Σ**_*j*_ as

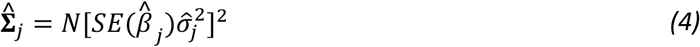

To estimate off-diagonal terms **Σ**_*jk*_, we now write *Y* = **Xβ** + *ϵ* = *X*_1_*β*_1_+… +*X*_*p*_*β*_*p*_ + *ϵ* = *X*_*j*_*β*_*j*_ + *X*_*k*_*β*_*k*_ + *ϵ*_*jk*_, where *ϵ*_*jk*_ = Σ_*i*:*i*∉{*j,k*}_ *X*_*i*_*β*_*i*_ + *ϵ*. Under the same assumption where the magnitude of SNP effects is very small, we can simplify **Σ**_*jk*_ by:

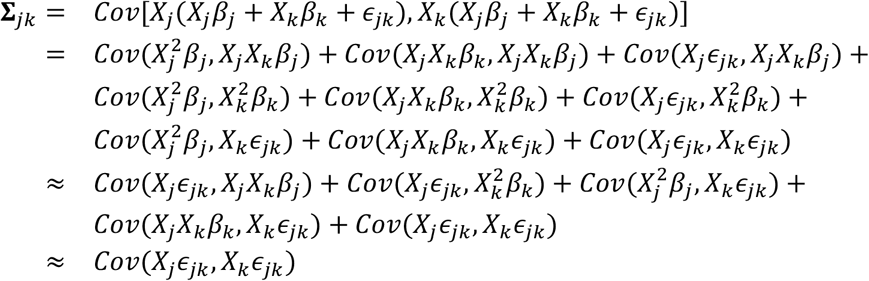

In a similar fashion, we further partition all SNPs in the genome other than the *j*-th and the *k*-th SNP into two sets. Let *S*_3_ denote the collection of SNPs that are independent from both the *j*-th and the *k*-th SNPs, and *S*_7_ includes the remaining SNPs that are in LD with either the *j*-th or the *k*-th SNP. Based on a similar rationale, we can safely assume that ∣*S*_3_∣ ≫ ∣*S*_4_∣. Then, by ignoring SNPs in *S*_4_ and thus treating *X*_*j*_ and *X*_*k*_ as being independent from *ϵ*_*jk*_, we express **Σ**_*jk*_ as:

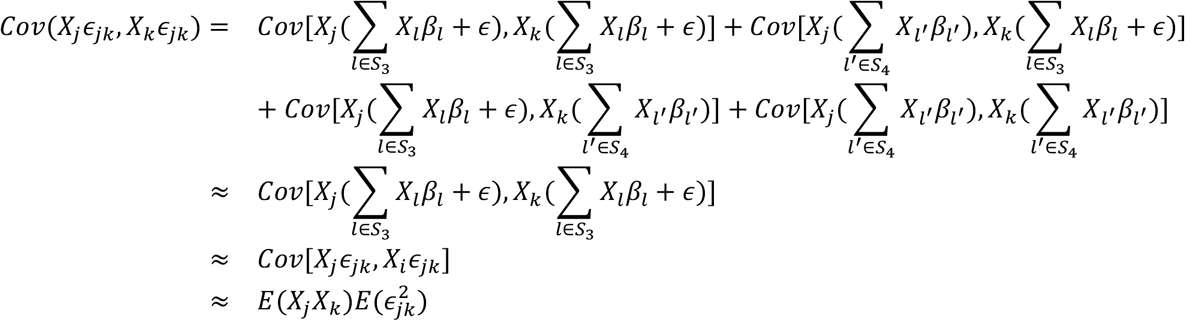

where *E*(*X*_*j*_*X*_*k*_) can be directly estimated by the LD correlation matrix and MAF-based SNP variance estimator. For 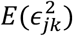, it is the residual variance from a two-SNP regression model and should be smaller than both 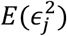 and 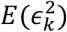. In practice, we can approximate it by the smaller value between 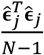 and 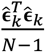 Therefore, the numerical approximation for **Σ**_*jk*_ becomes

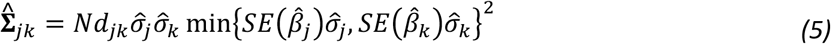

Now we can then generate summary statistic from the multivariate normal distribution in formula *(3)*. Note that our earlier subsampling framework is a special case where SNPs are independent and its only difference with the current method is the estimation of 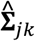. In the next session we will discuss how to subsample summary statistics efficiently from a multivariate normal distribution.

### Strategy for subsampling summary statistics

Next, we discuss how to partition full GWAS summary statistics into *K* independent subsets of GWAS samples, denoted as **x**^(1)T^**y**^(1)^, …, **x**^(<)T^**y**^(<)^ for *K* > 2. When *K* = 2, formula *(3)* can be directly applied to divide GWAS summary statistics into two independent sets. Otherwise, let *N*^(1)^, …, *N*^(*K*)^ denote the corresponding sample size for each subset of individuals and 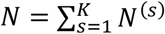. By formula *(3)*, we can subsample 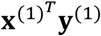 from **x**^*T*^**y** observed in the complete GWAS summary data. After that, we calculate summary statistics excluding *N*^(1)^ individuals from the first subset as 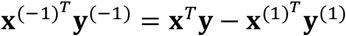. This technique of combining or subtracting independent sets of summary statistics has been commonly utilized by methods such as METAL and Metasubtract(65, 66). To generate summary statistics for any following subset numbered *t* + 1 (i.e., 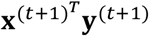) for *t* = 1, …, *K* − 2, we update the conditional distribution in *(3)* with the new “full” GWAS summary statistics and correspondent total sample size:

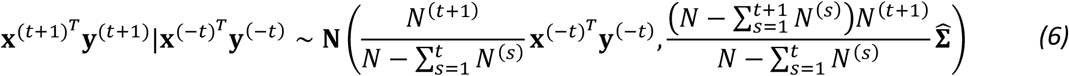

where 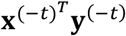 represents summary statistics excluding first *t* subsets of individuals. This subsampling strategy guarantees that every subset is independent from each other and avoids overfitting when *K* > 2. Finally, for the last subset *K*, we can directly calculate its summary statistics by 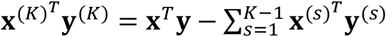. Together, this is a flexible framework for generating summary statistics and can be used for various types of PRS analyses as we discuss in later sections.

It is a difficult task to subsample summary statistics for all SNPs in the genome simultaneously given the large dimension of genotype and imputed data. Even if PRS modeling is restricted to HapMap3 SNPs, it remains challenging to subsample 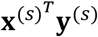 for more than one million SNPs altogether(26). To efficiently generate data, we partition the whole genome into approximately independent LD blocks and subsample summary statistics for SNPs in each LD block separately(67, 68). Then 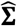 becomes a sparse block-diagonal matrix, i.e., 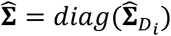. Within each LD block, the empirical SNP correlation matrix may not always be positive-definite and thus making it impossible to randomly generate data from that LD block. A straightforward remedy is to conduct eigen decomposition for any 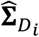 that is negative definite, manually change negative eigenvalues to 0’s, and obtain an approximation of 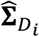 that is positive semi-definite. Note that this may not be the best approach and other methods for estimating LD blocks can also be applied(69, 70).

### Evaluate predictive performance of PRS

Here, we generalize the summary-statistics-based PRS evaluation scheme proposed in our previous work to incorporate LD. We denote PRS as a weighted sum of allele counts across many SNPs:

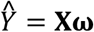

where **ω** ∈ ℝ^*p*^ is a vector of SNP weights, which can be marginal regression coefficients from GWAS or post-hoc effect size estimates. If individual-level data is available, then *R*^2^ evaluated on any holdout dataset (**y**^(*s*)^, **x**^(*s*)^) can be calculated as

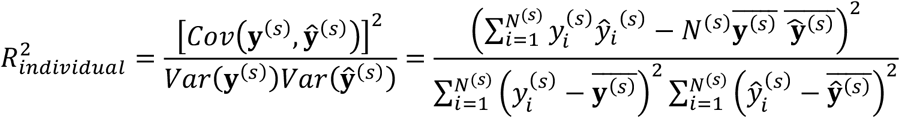

where *ŷ*_*i*_ is the PRS for the *i*-th person, 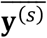 is the mean phenotypic value, and 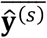 is the mean PRS value in holdout dataset *s*. On the other hand, we have shown that when only summary statistics of the holdout dataset is available and SNPs are independent, 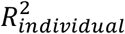 can be approximated by(38):

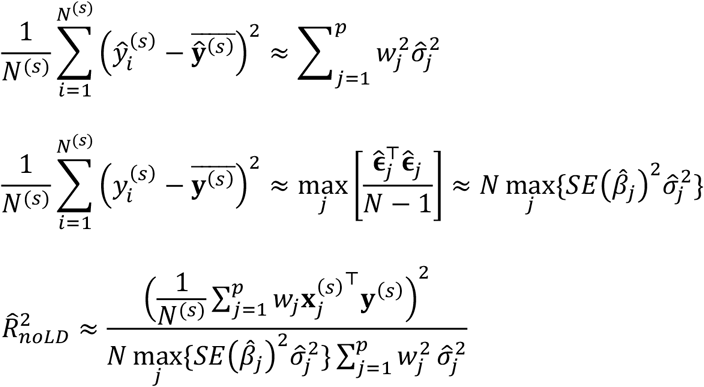

given that **x**^(*s*)^, **y**^(*s*)^, and ŷ^(*s*)^ are centered. In practice, we use the 90% quantile instead of 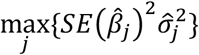 to get a robust estimate of *Var*(**y**^(*s*)^). When LD is present, the approximations for *Cov*(**y**^(*s*)^, ŷ^(*s*)^) and *Var*(**y**^(*s*)^) remain the same. For *Var*(ŷ^(*s*)^), it can now be approximated by **ω**^*T*^*Var*(**x**^(*s*)^)**ω**, with *Var*(**x**^(*s*)^) estimated using the LD correlation matrix and MAF calculated from the reference panel. Taken together, we have

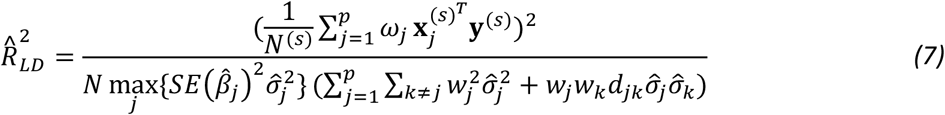

Note that similar versions of this formula have been tested and applied in the literature(22, 40, 41). In practice, we can directly calculate PRS on the LD reference genotype data and use the sample variance of PRS to replace 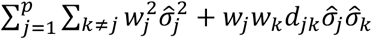 for optimal computational efficiency.

### The PUMAS framework

Given the flexible framework we introduced for subsampling GWAS summary data and evaluating PRS based on summary statistics, PUMAS becomes a special case where the entire GWAS summary-level data is partitioned into a training and a tuning dataset, denoted as 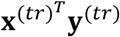 and 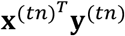. PUMAS first draws 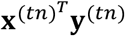 from *(3)* and then calculates 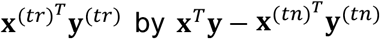. For each SNP, the marginal effect size and its standard error from the training set can be calculated as

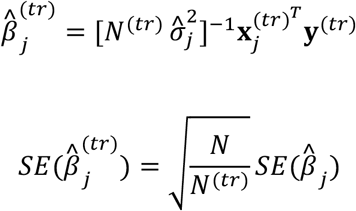

Then these summary statistics from the training dataset can be used to train any PRS methods that use GWAS summary statistics as input. *R*^2^ of the PRS model assessed on the fine-tuning dataset can be approximated by replacing 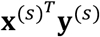 with 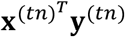 and changing the corresponding sample size in formula *(7)*. This procedure can be repeated *k* times to implement a *k*-fold Monte Carlo cross-validation (MCCV) to select the best-performing tuning parameter. When there is a set of tuning parameters **λ** in a PRS framework, that is, *Ŷ*(*λ*) = **Xω**(*λ*), PUMAS chooses the optimal tuning parameter 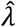 by

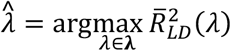

where 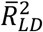 denotes the mean 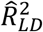 across *k* -fold MCCV. This cross-validation technique also applies to models that are hyperparameter-free or fine-tuned in advance. When the goal is to pick the best PRS model among a total of *M* PRS methods, the best model 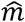 can be selected by

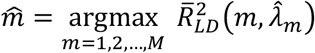

where 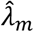 is the besting tuning parameter for PRS framework *m*.

### Combining multiple PRSs with PUMAS-ensemble

Next, we introduce PUMAS-ensemble, an extension of PUMAS that applies ensemble learning to combine multiple PRS using GWAS summary statistics. To do this, PUMAS-ensemble further partitions the full GWAS association results to 4 independent sets of summary statistics corresponding to training 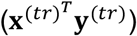, tuning 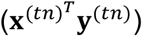, ensemble training 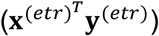, and testing 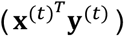 summary statistics. Using formula *(6)*, we subsample summary statistics iteratively and compute 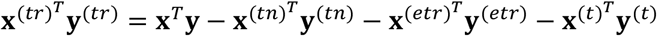. Like PUMAS, PUMAS-ensemble first conducts *k*-fold MCCV using training and tuning summary statistics to pick the best tuning parameter for each PRS method. Then, it trains each optimal PRS model’s weight on the ensemble training data and evaluates the combined PRS on the testing summary statistics. A straightforward and intuitive way of combining PRS is through multiple linear regression. However, if individual-level genotype and phenotype data is not available, we cannot fit the regression in the conventional way. Below we illustrate how to calculate regression coefficients using summary-level data alone. We define the multiple linear regression model on the ensemble training dataset as:

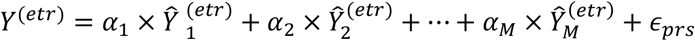

where **α** = [*α*_1_ *α*_2_ … *α*_*M*_]^*T*^ are PRS weights for *M* PRS methods. We also define

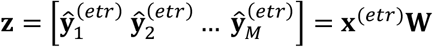

as the observed PRS matrix with dimension *N*^(*ert*)^ × *M*, and **W** = [**w**_1_ **w**_2_ … **w**_*M*_] are a *p* × *M* SNP weights matrix for *p* SNPs from *M* methods. To obtain the least squares estimator of **α**, that is 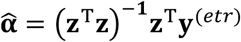, we need to estimate **z**^*T*^**z** and **z**^T^**y**^(*ert*)^ separately. In fact, under the assumption that genotype and phenotype are both centered, we can show that

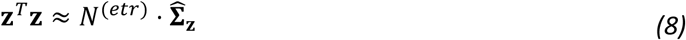

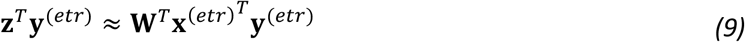

where 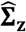 is the empirical covariance matrix of the PRS matrix **z**. In practice, we can estimate 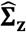 by calculating PRSs and their sample covariance matrix on a reference LD genotype dataset or approximate it by computing **W**^*T*^**DW**. Taken *(8)* and *(9)* together, we can estimate PRS weights using only summary statistics. Then we take the average PRS weights across *k* folds, i.e., 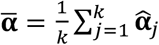, and report it as the PRS weight to combine optimized PRSs. Finally, we modify equation *(7)* to calculate predictive *R*^2^ for ensemble PRS on the testing summary-level data:

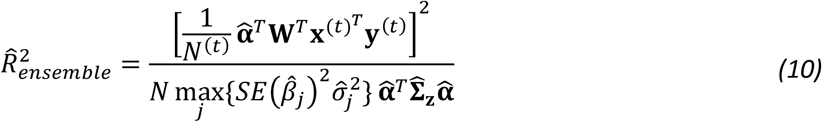

In the end, PUMAS-ensemble reports the average prediction accuracy of ensemble PRS across *k* folds. Note that PUMAS-ensemble can benchmark all PRS models in addition to the ensemble PRS on the testing summary statistics since it is independent from training and tuning datasets. Given sufficient data for model training, the ensemble learning model should always outperform fine-tuned PRS identified through grid-search. This is because grid search result is a special case in the ensemble learning, with the weight set to be 1 for a particular PRS and 0 for all other PRS models. Instead, the ensemble learning approach fits a regression to identify optimal weight values to maximize the predictive performance of ensemble score. Taken together, PUMAS-ensemble is a highly flexible framework to train, fine-tune, combine, and evaluate PRS models based on GWAS summary statistics.

### Binary phenotypes

There are two challenges when applying PUMAS and PUMAS-ensemble to binary phenotypes. First, summary statistics obtained from logistic regression frameworks violate the linear regression model assumption in our derivation. Therefore equations *(3)* and *(6)* are not directly appliable to subsampling summary statistics for binary traits because **X**^*T*^Y calculation is non-trivial for log odds ratios. Second, squared Pearson correlation between a binary outcome and PRS using logistic regression coefficients as input is less interpretable and rarely reported. On the other hand, area under the ROC curve (AUC) is often the preferred metric to quantify PRS accuracy for binary outcome. AUC calculation based on summary statistics has been developed but is not yet generalized to handle whole genome data, making it difficult to evaluate more sophisticated PRS methods that leverage contributions from millions of SNPs when individual-level data is not accessible(71). Here we propose a simple solution that allows us to apply PUMAS and PUMAS-ensemble to binary phenotypes and report interpretable *R*^2^. For binary traits, *R*^2^ on the observed scale (i.e., 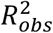) has been defined and discussed in the literature as an alternative metric for evaluating PRS prediction accuracy(49). 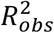 is the squared correlation between PRS and 0-1 status where PRS uses effect sizes estimated from linear probability model (LPM, i.e., linear regression between the binary response and SNP allele counts) as inputs(72). If GWAS summary-level data is acquired from linear probability model, then PUMAS and PUMAS-ensemble can be directly applied to calculate 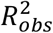 for binary traits(60). When LPM summary statistics are not available, since a single SNP has very weak effect on the phenotypic outcome in practice, we can still safely approximate LMP coefficient estimations using Z-score from logistic regression(47, 48). Specifically, we can calculate 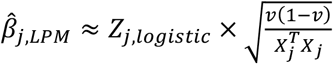 where *Zj,logistic* is Z-score for the *j*-th SNP from logistic summary statistics and *v* is the sample prevalence. Then, we can use 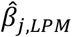 and correspondent standard error 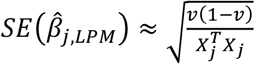 to apply PUMAS and PUMAS-ensemble to dichotomous phenotypes. Eventually, if it is preferred to transform 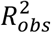 to *R*^2^ on the liability scale 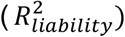 which can be comparable across different studies and phenotypes, such transformation has been developed using sample and population prevalence(49).

### Sample size imputation

In this section, we discuss how to handle sample size misspecification in GWAS summary statistics when applying our approach. Sample size misspecification is common in published GWAS datasets since many studies often do not report SNP-specific sample size and only provide a maximum sample size for the entire study. This is sub-optimal for PRS training if variant-level samples sizes differ substantially (e.g., in meta-analysis). A recent study has extensively investigated sample size misspecification in marginal association statistics and observed consistently decreased PRS prediction accuracy when the issue is not properly addressed(31). For PUMAS and PUMAS-ensemble, incorrect sample sizes will both affect the quality of subsampled summary statistics and bias the estimation of predictive *R*^2^. To address this issue, we employed the approach proposed in Privé et al. to impute and conduct quality control on variant-specific sample size(31). Specifically, when the summary-level data does not provide sample size information for each SNP, we first impute sample size and remove SNPs with imputed sample size smaller than 70% and larger than 110% of reported maximum sample size. For summary statistics that provides per-SNP sample sizes, we simply removed variants with sample size smaller than 70% of the largest sample size. On the other hand, to make sure formula *(7)* and *(10)* work for summary statistics with varying SNP-specific sample sizes, we enforce all summary statistics other than training summary statistics to have the same sample size for every SNP. We achieve this by subsampling all other summary statistics first where we can specify subset size and calculate 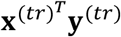 at last.

### PRS training

We trained lassosum, PRS-CS, and LDpred2 models for all PRS analyses in this study(22, 25, 26). lassosum is a penalized regression framework that trains lasso regression coefficients for SNPs in each LD blocks with tuning parameters *s* and *λ*, where *s* controls the sparsity of LD matrix and *λ* is the penalty term that regularizes shrinkage of effect sizes. PRS-CS and LDpred2 are both Bayesian PRS frameworks with different prior assumptions for the SNP effect size distribution. PRS-CS has a global shrinkage parameter *ϕ* that uniformly shrinks its continuous prior distribution for each SNP and includes a fully Bayesian approach that automatically learns *ϕ* during model fitting. LDpred2 is an extension of LDpred that places a point normal prior on SNP effects based on tuning parameter *p* that represents the proportion of causal variants in the genome (LDpred non-inf and LDpred2_grid) or a univariate normal prior on all SNPs that doesn’t require model-tuning (LDpred/LDpred2-Inf)(12). Like PRS-CS, LDpred2 can also employ an empirical Bayesian approach to optimize *p* on the training summary statistics. For implementation, we trained PRS-CS (v1.0.0) models using UKB European LD reference for simulation study and 1000 Genomes European LD reference for real data analysis. We followed PGS server pipeline to implement lassosum (R package ‘lassosum’ v0.4.5) and LDpred2 (R package ‘bigsnpr’ v1.9.11)(22, 28, 73). Due to larger computational burden, we implemented LDpred2 on each chromosome separately and only used the estimated heritability from LD-score regression as the tuning parameter *h*^2^ in LDpred2(50). For real data analysis in UKB we constructed both non-sparse and sparse versions of LDpred2 models. We employed more shrinkage on LDpred2-auto model (shrink_corr = 0.5) and LDpred2_grid models (low_h^2^=0.1*h^2^) when analyzing publicly available GWAS summary statistics to ensure model convergence. The best tuning parameter for lassosum was obtained through grid search. For LDpred2 and PRS-CS, we compared grid search with empirical Bayesian models to find the best parameter.

In addition, we trained SDPR(43) and MegaPRS(40) models for all ensemble PRS analyses and PUMAS-ensemble simulation, respectively. SDPR is a recently developed Bayesian nonparametric PRS model that is computationally efficient and tuning-free. We fitted SDPR models using its latest v0.9.1 release on github with provided 1000 Genomes Project EUR LD reference. MegaPRS is a flexible Bayesian PRS framework that can employ multiple different prior specifications. The MegaPRS model with BayesR prior includes 84 sets of tuning parameters that determine the relative weights of various Gaussian components. We fitted MegaPRS models using the LDAK software (v5.2) with LDAK-thin heritability estimation(74). We only trained PRS models on HapMap3 SNPs in all analyses throughout this study.

### Simulation settings

We conducted simulations using UKB genotype data(44) imputed to the Haplotype Reference Consortium reference. We removed samples who are not of European ancestry and genetic variants with MAF below 0.01, imputation *R*^2^ below 0.9, Hardy-Weinberg equilibrium test p-value below 1e-6, or missing genotype call rate greater than 2%. We further extracted variants in the HapMap3 SNP list and 1000 Genomes Project Phase III LD reference data for European ancestry from PRS-CS. 377,509 samples and 944,547 variants remained after quality control. Then, we randomly selected 100,000 samples to be the training dataset and 1,000 samples as the LD genotype reference for our summary-statistics-based approach. To generate trait values, we simulated true effect sizes from a point normal distribution, i.e., 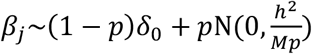 where *p* is the proportion of causal variants, *δ*_0_ is point mass at 0, *h*^2^ is the total heritability of the phenotype, and *M* is the total number of SNPs(7, 12). We did not simulate associations between SNP true effects on the allelic scale and MAF since previous analysis has shown minimal difference in performance between PUMAS and PRS validation using individual-level data(38, 74). We chose *p* to be 0.1% and 20% corresponding to sparse and polygenic genetic models, and *h*^2^ = 0.2, 0.5, 0.8 to create a total of 6 simulation settings with various types of genetic architecture. Within each setting, we randomly selected causal variants across the whole genome. Then we simulated quantitative traits by adding up the SNP allele counts weighted by their true effect sizes and randomly generated gaussian noises scaled based on trait heritability. We fitted marginal linear regression in PLINK to obtain GWAS summary statistics in each setting(75).

We compared PUMAS-ensemble with 4-fold MCCV. To implement 4-fold MCCV, in each fold we randomly selected 60% of all samples to form the training dataset (N=60,000), 20% as the tuning dataset (N=20,000), 10% as the ensemble training dataset (N=10,000), and the remaining 10% as the testing dataset (N=10,000). We conducted GWAS on the training data and used summary statistics to train PRS models, fine-tuned PRS methods on the tuning data, obtained optimized PRSs’ weights in the ensemble score by fitting multiple linear regression on the ensemble training data, and finally evaluated each PRS model’s predictive *R*^2^ on the testing data. For PUMAS-ensemble, we first used all samples (N=100,000) to fit marginal linear regression and obtained the full summary statistics. In a similar fashion, we partitioned the full summary statistics to training summary data (N=60,000), tuning summary data (N=20,000), ensemble learning summary data (N=10,000) and testing summary data (N=10,000) for corresponding PRS analysis. Similarly, we compared PUMAS with 4-fold MCCV by using only the training and tuning summary-level and individual-level data for two approaches, respectively. We included lassosum, LDpred2, and PRS-CS in all simulation analysis, and added SDPR and MegaPRS in PUMAS-ensemble simulations. In all simulations, we used 1000 Genomes Project European LD dataset provided by the PRS-CS software to subsample summary statistics. lassosum, LDpred2, and MegaPRS model training used the holdout UKB LD genotype data (N=1,000) as the LD reference. We implemented SDPR, lassosum with *s* = 0.2, 0.5, 0.9 and *λ* = 0.005, 0.01, PRS-CS with *ϕ* = 0.0001, 0.01, *auto*, and LDpred2 with *p* = 0.001, 0.01, 0.1, *auto*, and the infinitesimal model. For MegaPRS, to ensure robust model convergence in all simulation settings, we included 9 models with distinct tuning parameters {p1,p2,p3,p4} = {0.99,0.01,0,0}, {0.95,0.05,0,0}, {0.9,0.1,0,0}, {0.8,0.1,0.05,0.05}, {0.7,0.1,0.1,0.1}, {0.6,0.2,0.1,0.1}, {0.5,0.2,0.2,0.1}, {0.4,0.2,0.2,0.2}, {0,0,0,1}. We repeated this procedure four times and calculated average *R*^2^ to pick the best set of tuning parameters for both approaches.

We conducted additional simulations to demonstrate that PUMAS and PUMAS-ensemble can be applied to binary traits. For each setting in the quantitative simulation study, we dichotomized the continuous phenotype (i.e., true liability value under a liability threshold model) using either the median or 90% quantile to acquire balanced (5-to-5) and unbalanced (1-to-9) case-control ratios. Therefore, we have a total of 12 binary simulation settings. We fitted logistic regressions in PLINK to obtain GWAS summary statistics in each setting and transformed logistic regression summary statistics to the linear scale(47, 48, 75). We then compared PUMAS/PUMAS-ensemble with MCCV using *R*^2^ computed on the observed scale (i.e., *R*^2^ between PRS and 0-1 status).

We conducted two additional simulation analyses for PUMAS’s subsampling scheme. First, we investigated the similarity between PUMAS and MegaPRS under two simulation settings with heritability of 0.5 from six quantitative trait simulation scenarios described above. Since MegaPRS only uses z-scores as its input and output, we focused on z-scores in this simulation. As a benchmark, we compared both approaches with MCCV where we randomly selected a subset of individuals and obtained SNP association statistics from regression analysis. For each approach, we subsampled SNP z-scores based on 75% of samples (total N = 100,000) and repeated this procedure 100 times. We summarized the results for randomly selected 5 causal variants and 5 non-causal variants in each simulation. Second, we investigated the robustness of PUMAS under an extremely sparse simulation setting. We followed the same simulation strategy and set number of causal variants to be 10 on chromosome 1 and heritability to be 0.1. For both approaches, we subsampled summary statistics based on 75% of individuals and repeated this procedure 100 times. We compared the distribution of summary statistics generated from PUMAS and MCCV for each causal SNP.

### UKB data analysis

We applied our approach to 16 quantitative traits, 4 diseases, and 1 ordinal trait in UKB. The list of UKB phenotypes is presented in **Additional file 11** and **12**: **Table. S10**-**S11**. The imputed UKB genotype data consists of 375,064 independent individuals of European ancestry and 1,030,187 variants after quality control. We used Hail (v0.2.57) to perform linear regression for quantitative and ordinal traits while adjusting for sex, age polynomials to the power of two, interactions between sex and age polynomials, and top 20 principal components. For 4 disease outcomes, we obtained GWAS summary statistics via regenie (v3.0.3) accounting for sex, age polynomials to the power of 3, interactions between sex and age polynomials, and top 10 principal components as recommended(76).

We compared PUMAS with external validation using a holdout subset of UKB samples. For external validation of quantitative traits, we randomly selected 38,521 samples with non-missing phenotypic measurements for all traits to form the holdout dataset. The remaining samples for each phenotype were used as training data. In this way, we implemented an approximately 9-to-1 training-testing split. Similarly for each binary and ordinal outcome, we continued to employ a 9-to-1 sample partition while matching the case-control ratio between the training and holdout datasets. Detailed sample size information for all traits is included in **Additional file 11** and **12**: **Table. S10**-**S11**. Then, we conducted GWAS on the training data and obtained summary statistics. For quantitative and ordinal traits, we computed and evaluated PRS models on the entire holdout set and reported predictive *R*^2^ between PRS and phenotypes with covariates regressed out. For disease traits, we constructed PRS models and calculated *R*^2^ on the observed scale using both linear probability model summary statistics and logistic model summary statistics. For all phenotypes, the holdout set of quantitative traits (N=38,521) was also used as LD reference data for PRS model training. For comparison, we applied PUMAS to partition the same GWAS summary-level data used in MCCV to 75% training summary statistics and 25% tuning summary statistics. We used the holdout dataset (N=38,521) for summary statistics subsampling(67) and as the LD reference for lassosum and LDpred2 model training. We estimated variance of PRS models based on a smaller subset (N=1,000) of the holdout data when evaluating PRS performance. This procedure was repeated 4 times and we reported the average *R*^2^ for each PRS model. In all analyses, we implemented lassosum with *s* = 0.2, 0.5, 0.9 and *λ* = 0.005, 0.01, PRS-CS with *ϕ* = 0.0001, 0.01, *auto*, LDpred2 with *p* = 0.001, 0.01, 0.1, *auto* and the infinitesimal model.

Next, we compared PUMAS-ensemble with the training-testing split approach for ensemble learning on the holdout dataset. For PUMAS-ensemble, we partitioned full GWAS summary statistics into training (60%), tuning (20%), and ensemble training (10%) summary statistics to train PRS models based on a grid of tuning parameters, select the best tuning parameter setting for each PRS method, and fit a second level regression to obtain regression weights for fine-tuned PRS models. We then randomly partitioned the holdout dataset into two equally sized subsets. We used PUMAS-ensemble to obtain PRS models’ regression weights and then constructed and evaluated the ensemble PRS on the second half of the holdout set. PRS models with negative weights were removed from linear combination. In comparison, for the training-testing split approach based on individual-level data, we used the first half of the holdout set to fit multiple linear regression to obtain regression coefficients for SDPR and fine-tuned lassosum, LDpred2, and PRS-CS scores. Then we computed and evaluated the ensemble PRS models on the second half of the holdout data. In all analyses, we trained SDPR, lassosum with *s* = 0.2, 0.9 and *λ* = 0.001, 0.01, 0.1, PRS-CS with *ϕ* = 0.0001, 0.01, *auto*, LDpred2 with *p* = 0.001, 0.01, 0.1, *auto* and the infinitesimal model. As a secondary analysis, we compared performance of PUMAS (70% training, 20% tuning, 10% testing) and PUMAS-ensemble (50% training, 20% tuning, 20% ensemble learning, 10% testing) on 16 quantitative traits in UKB. We benchmarked the best PRS model chosen by PUMAS and the ensemble score trained by PUMAS-ensemble on the second half of UKB holdout dataset.

To investigate how sensitive PUMAS-ensemble is to LD misspecification, we repeated PUMAS-ensemble analysis on 16 complex traits in UKB with different LD references. Previously, we used UKB LD panel; in this sensitivity analysis, we explored the impact of using 1000 Genomes Project Phase III European samples (1KG EUR) and East Asian samples (1KG EAS) as the LD reference panel, while keeping everything else unchanged. The 1KG LD reference data were prepared from our earlier work(42). We trained ensemble scores by PUMAS-ensemble using different LD reference panels and evaluated these scores on UKB holdout dataset. Results based on individual-level data were used as a benchmark of performance. In addition, we meta-analyzed UKB and Biobank Japan(77-79) (BBJ) GWAS summary statistics for 16 complex traits using METAL(65) and applied PUMAS-ensemble using either 1KG EUR or 1KG EAS data as the LD reference. Sample size information for BBJ GWAS summary statistics is included in **Additional file 17**: **Table. S16**. Similarly, we compared ensemble scores from PUMAS-ensemble and individual-level ensemble learning on the UKB holdout dataset. lassosum with *s* = 0.2, 0.9 and *λ* = 0.001, 0.01, 0.1, PRS-CS with *ϕ* = 0.0001, 0.01, *auto*, and LDpred2 with *p* = 0.001, 0.01, 0.1, *auto* and the infinitesimal model were considered for ensemble PRS training in this analysis.

### Building a catalog of PUMAS-ensemble ensemble scores

We applied PUMAS-ensemble to a collection of publicly available GWAS summary statistics. We selected complex diseases and traits with a minimal case sample size of 5,000 and a total sample size of 50,000, respectively. We excluded studies that performed GWAS on related samples and retained traits with significant heritability estimation (p-value below 0.05) from LD score regression(50). In the end, we obtained a list of 31 GWAS summary statistics including 23 binary outcomes and 8 complex traits as summarized in **Additional file 18**: **Table. S17**. For each summary statistics, we kept HapMap 3 SNPs that passed a series of quality control criteria listed in **Additional file 19**: **Table. S18**, including transformation of logistic summary statistics and imputation of per-SNP sample size. Then we applied PUMAS-ensemble to each phenotype to implement 4-fold MCCV by partitioning the summary statistics to training (60%), tuning (20%), ensemble training (10%), and testing (10%) datasets. We used 1000 Genomes Project Phase III European samples as the LD panel for summary statistics subsampling, PRS model fitting and benchmarking. We implemented SDPR, lassosum with *s* = 0.2, 0.5, 0.9 and *λ* = 0.005, 0.01, PRS-CS with *ϕ* = 0.0001, 0.01, *auto*, LDpred2 with *p* = 0.001, 0.01, 0.1, *auto* and the infinitesimal model. We reported average predictive *R*^2^ of ensemble PRS, the best single PRS model, PRS-CS-auto and LDpred2-auto on the testing summary statistics.

We conducted additional analysis to investigate the validity of predictive *R*^2^ of ensemble PRS for Alzheimer’s disease. We used IGAP 2019 Alzheimer’s GWAS summary statistics to train PRS models and included 2,600 Alzheimer’s disease cases of European ancestry from the UKB cohort in the external validation dataset(51). The data fields used for Alzheimer’s cases extraction are presented in **Additional file 21**: **Table. S20**. We randomly selected 5,200 independent UKB samples not diagnosed with Alzheimer’s disease to use as healthy controls to match the case-control ratio in the IGAP 2019 study. Together, we obtained a UKB external validation dataset with 7,800 samples in total. We applied PUMAS to IGAP 2019 GWAS summary-level data and compared its performance with external validation. We compared *R*^2^ from both approaches with and without removing the *APOE* region from GWAS summary statistics. We excluded the *APOE* region from PRS analysis by removing variants between base pairs 45,116,911 and 46,318,605 (hg19) on chromosome 19.

## Declarations

### Ethics approval

Not applicable.

## Availability of data and materials

The UKB data were downloaded from UK Biobank Resource (https://www.ukbiobank.ac.uk) under application number 42148(44). PUMAS/ PUMAS-ensemble software is freely available at https://github.com/qlu-lab/PUMAS(80). The source code for PUMAS/ PUMAS-ensemble used in this study is deposited at https://zenodo.org/records/13826837 with DOI: 10.5281/zenodo.13826837. The PUMAS/ PUMAS-ensemble package is under MIT license.

## Competing interests

The authors declare no competing interests.

## Funding

The authors gratefully acknowledge research support from National Institutes of Health (NIH) grants U01 HG012039 and R21 AG067092, and support from the University of Wisconsin-Madison Office of the Chancellor and the Vice Chancellor for Research and Graduate Education with funding from the Wisconsin Alumni Research Foundation (WARF). We also acknowledge use of the facilities of the Center for Demography of Health and Aging at the University of Wisconsin-Madison, funded by NIA Center Grant P30 AG017266.

## Author’s contributions

Z.Z. and Q.L. conceived and design the study.

Z.Z. developed the statistical framework.

Z.Z., T.G., and M.Y. performed statistical analyses.

Z.Z. and Y.W. wrote the software.

S.Z. assisted in preparing and curating summary statistics.

J.M. assisted in developing ensemble PRS approach.

J.M. and Y.W. assisted in UKB data preparation.

J.S. assisted in developing statistical method for subsampling summary statistics.

Q.L. advised on statistical and genetic issues.

Z.Z. and Q.L. wrote the manuscript.

All authors contributed to manuscript editing and approved the manuscript.

## Acknowledgements

We thank members of the Social Genomics Working Group at University of Wisconsin for helpful comments. This research has been conducted using the UK Biobank Resource under Application 42148. This study makes use of summary statistics from many GWAS consortia. We thank many GWAS investigators for providing publicly available GWAS summary-level datasets.

## Supplementary information

Additional file 1:

Supplementary figures.

Additional file 2: Table S1

PUMAS-ensemble quantitative simulation results in UKB.

Additional file 3: Table S2

PUMAS-ensemble binary simulation results in UKB.

Additional file 4: Table S3

PUMAS quantitative simulation results in UKB.

Additional file 5: Table S4

PUMAS binary simulation results in UKB.

Additional file 6: Table S5

PUMAS-ensemble quantitative simulation results in UKB (including MegaPRS).

Additional file 7: Table S6

PUMAS-ensemble binary simulation results in UKB (including MegaPRS).

Additional file 8: Table S7

Benchmark computational requirement at different stages of PUMAS-ensemble.

Additional file 9: Table S8

Simulated Z scores from different approaches in quantitative trait simulation.

Additional file 10: Table S9

Simulated summary statistics from PUMAS and MCCV under extremely sparse simulation setting.

Additional file 11: Table S10

Description of UKB quantitative traits.

Additional file 12: Table S11

Description of UKB binary and ordinal traits.

Additional file 13: Table S12

Compare PRS prediction accuracy between PUMAS and external validation for quantitative traits in UKB.

Additional file 14: Table S13

Compare PRS prediction accuracy between PUMAS and external validation for binary traits in UKB.

Additional file 15: Table S14

Compare PRS prediction accuracy between PUMAS-ensemble and external validation in UKB.

Additional file 16: Table S15

Compare fine-tuned PRS and ensemble PRS based on different data partitioning on UKB holdout data.

Additional file 17: Table S16

Compare PRS prediction accuracy between PUMAS-ensemble (with LD misspecification) and external validation in UKB.

Additional file 18: Table S17

Description of 31 GWAS summary statistics used in PUMAS-ensemble catalog.

Additional file 19: Table S18

PUMAS-ensemble catalog GWAS summary statistics quality controls.

Additional file 20: Table S19

PRS prediction accuracy in PUMAS-ensemble catalog.

Additional file 21: Table S20

UKB data fields for AD cases extraction.

Additional file 22: Table S21

IGAP 2019 AD PRS prediction accuracy.

## Notes

### Competing Interest Statement

The authors have declared no competing interest.

### Summary of Updates

Manuscript revised and supplemental materials updated

