## Supplementary figures for "Optimizing and benchmarking polygenic risk scores with GWAS summary statistics"

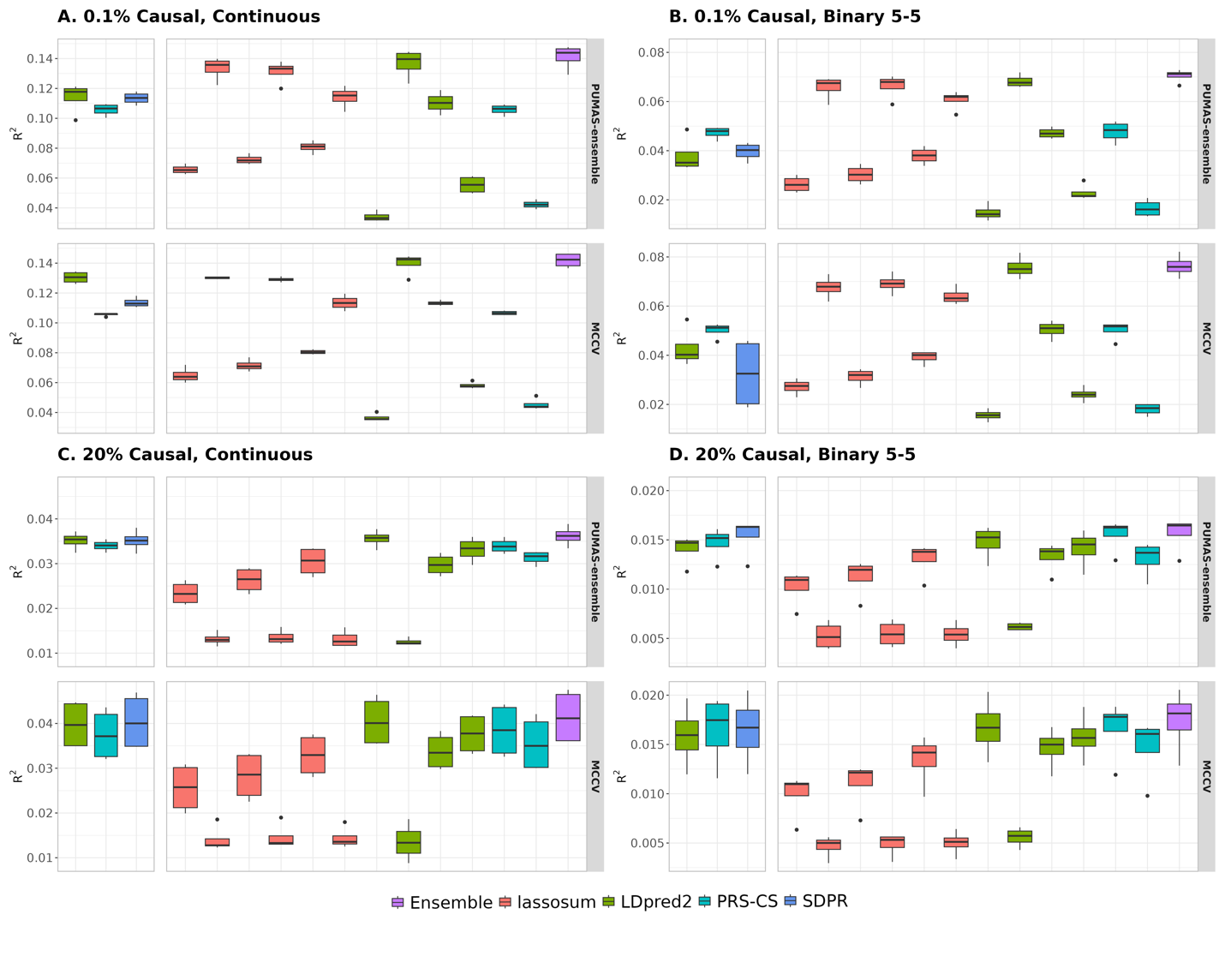

**Fig S1. Comparison of PUMAS-ensemble and MCCV in UKB simulation when heritability is 0.2**. (**A** and **C**) Simulation results for quantitative traits. (**B** and **D**) Simulation results for binary traits with balanced case-control ratio. Proportion of causal variants is 0.1% in **A** and **B**, and 20% in **C** and **D**. Models that do not require fine-tuning are shown on the left side of each panel. Y-axis: predictive $R^{2}$ across 4 repeats of MCCV; X-axis (left to right): tuning-free models: LDpred2-auto (green box), PRS-CS-auto (blue box), and SDPR (dark-blue box). lassosum models (red boxes) with tuning parameter settings: s=0.2 and λ=0.005, s=0.2 and λ=0.01, s=0.5 and λ=0.005, s=0.5 and λ=0.01, s=0.9 and λ=0.005, s=0.9 and λ=0.01. LDpred2 models (green boxes): non-infinitesimal with p=0.1, non-infinitesimal with p=0.01, non-infinitesimal with p=0.001, and infinitesimal model. PRS-CS (blue boxes): $\phi$=0.01 and 0.0001. Finally, the purple box shows the results of ensemble PRS.

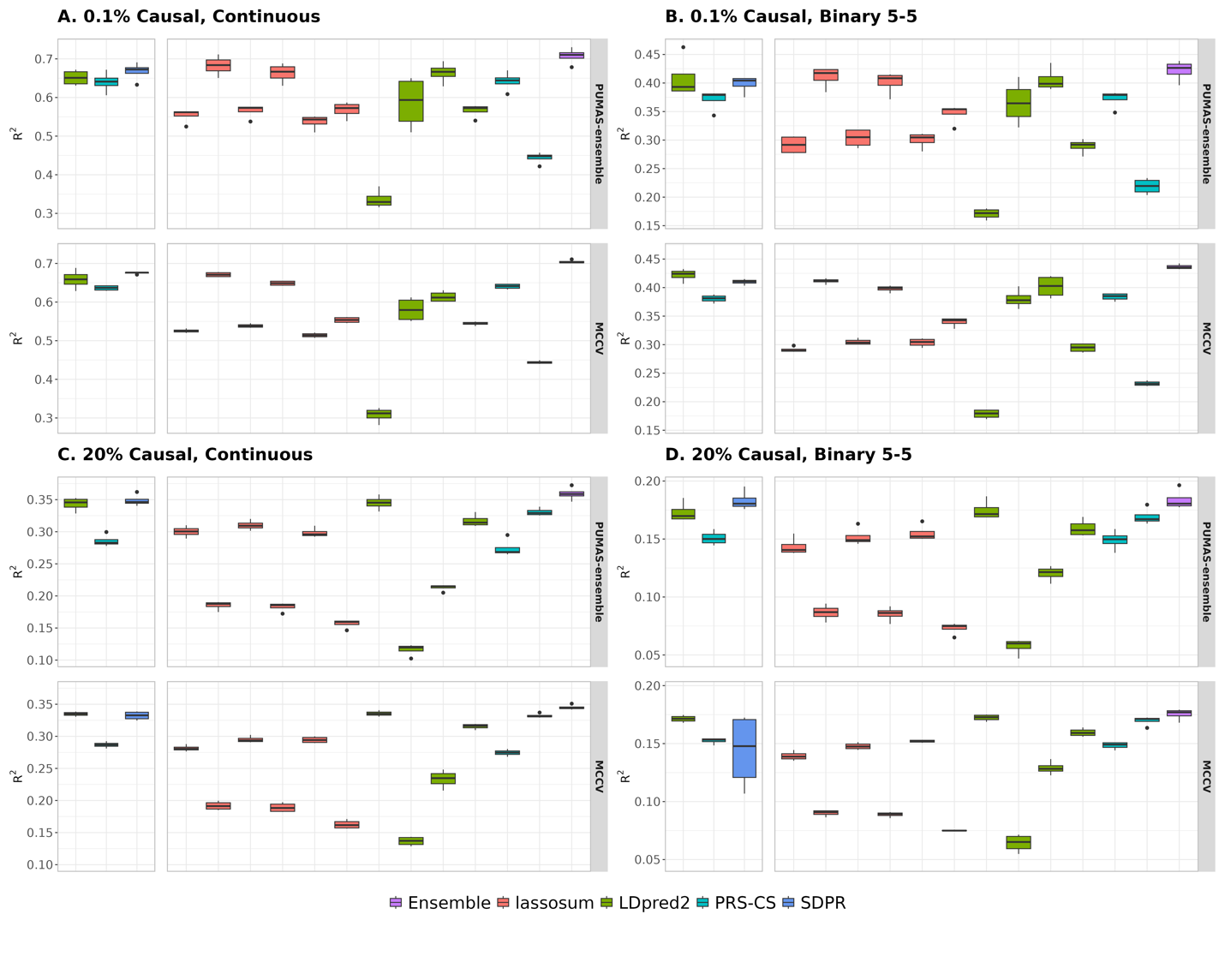

**Fig S2. Comparison of PUMAS-ensemble and MCCV in UKB simulation when heritability is 0.8**. (**A** and **C**) Simulation results for quantitative traits. (**B** and **D**) Simulation results for binary traits with balanced case-control ratio. Proportion of causal variants is 0.1% in **A** and **B**, and 20% in **C** and **D**. Models that do not require fine-tuning are shown on the left side of each panel. Y-axis: predictive $R^{2}$ across 4 repeats of MCCV; X-axis (left to right): tuning-free models: LDpred2-auto (green box), PRS-CS-auto (blue box), and SDPR (dark-blue box). lassosum models (red boxes) with tuning parameter settings: s=0.2 and λ=0.005, s=0.2 and λ=0.01, s=0.5 and λ=0.005, s=0.5 and λ=0.01, s=0.9 and λ=0.005, s=0.9 and λ=0.01. LDpred2 models (green boxes): non-infinitesimal with p=0.1, non-infinitesimal with p=0.01, non-infinitesimal with p=0.001, and infinitesimal model. PRS-CS (blue boxes): $\phi$=0.01 and 0.0001. Finally, the purple box shows the results of ensemble PRS.

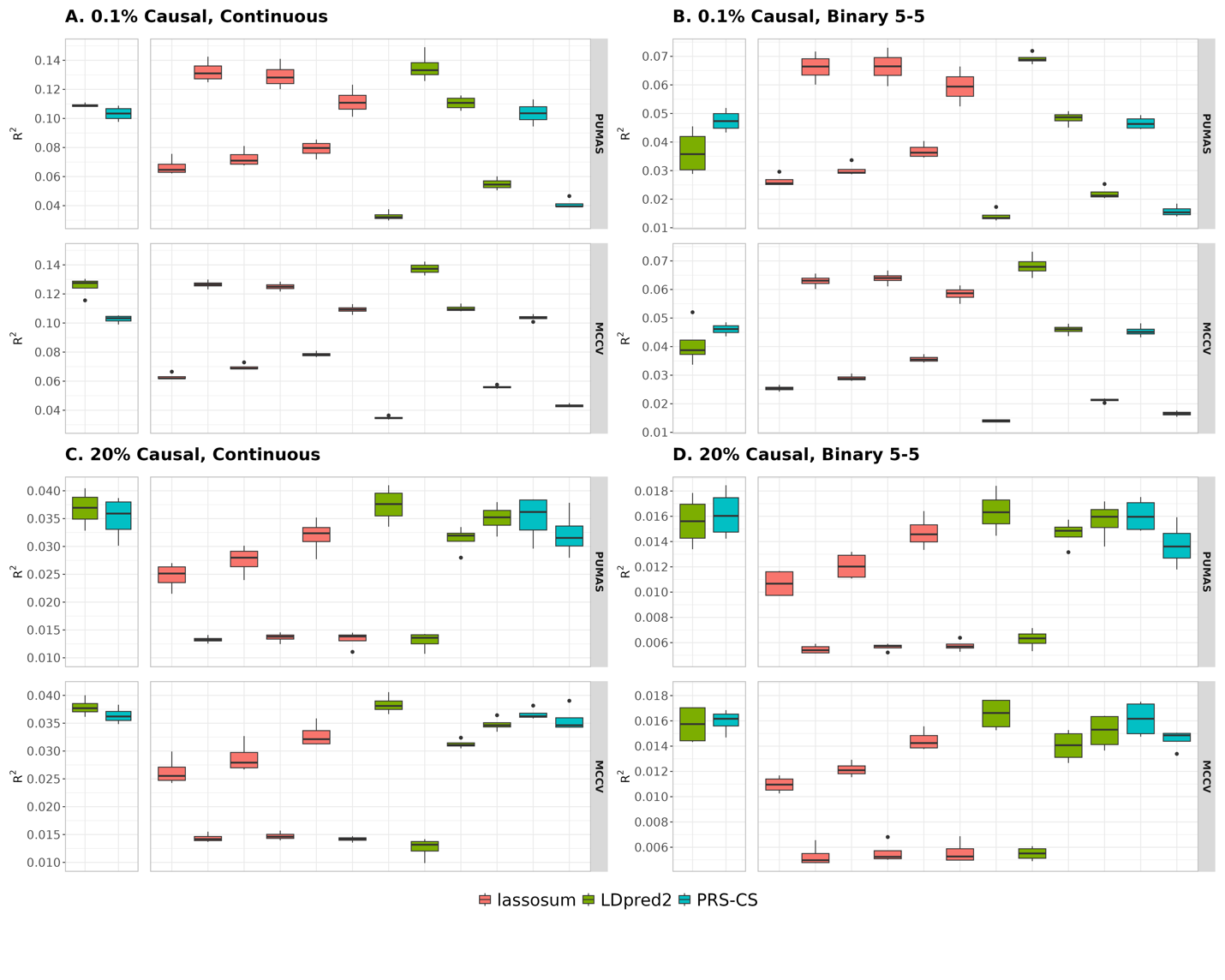

**Fig S3. Comparison of PUMAS and MCCV in UKB simulation when heritability is 0.2**. (**A** and **C**) Simulation results for quantitative traits. (**B** and **D**) Simulation results for binary traits with balanced case-control ratio. Proportion of causal variants is 0.1% in **A** and **B**, and 20% in **C** and **D**. Models that do not require fine-tuning are shown on the left side of each panel. Y-axis: predictive $R^{2}$ across 4 repeats of MCCV; X-axis (left to right): tuning-free models: LDpred2-auto (green box) and PRS-CS-auto (blue box). lassosum models (red boxes) with tuning parameter settings: s=0.2 and λ=0.005, s=0.2 and λ=0.01, s=0.5 and λ=0.005, s=0.5 and λ=0.01, s=0.9 and λ=0.005, s=0.9 and λ=0.01. LDpred2 models (green boxes): non-infinitesimal with p=0.1, non-infinitesimal with p=0.01, non-infinitesimal with p=0.001, and infinitesimal model. PRS-CS (blue boxes): $\phi$=0.01 and 0.0001.

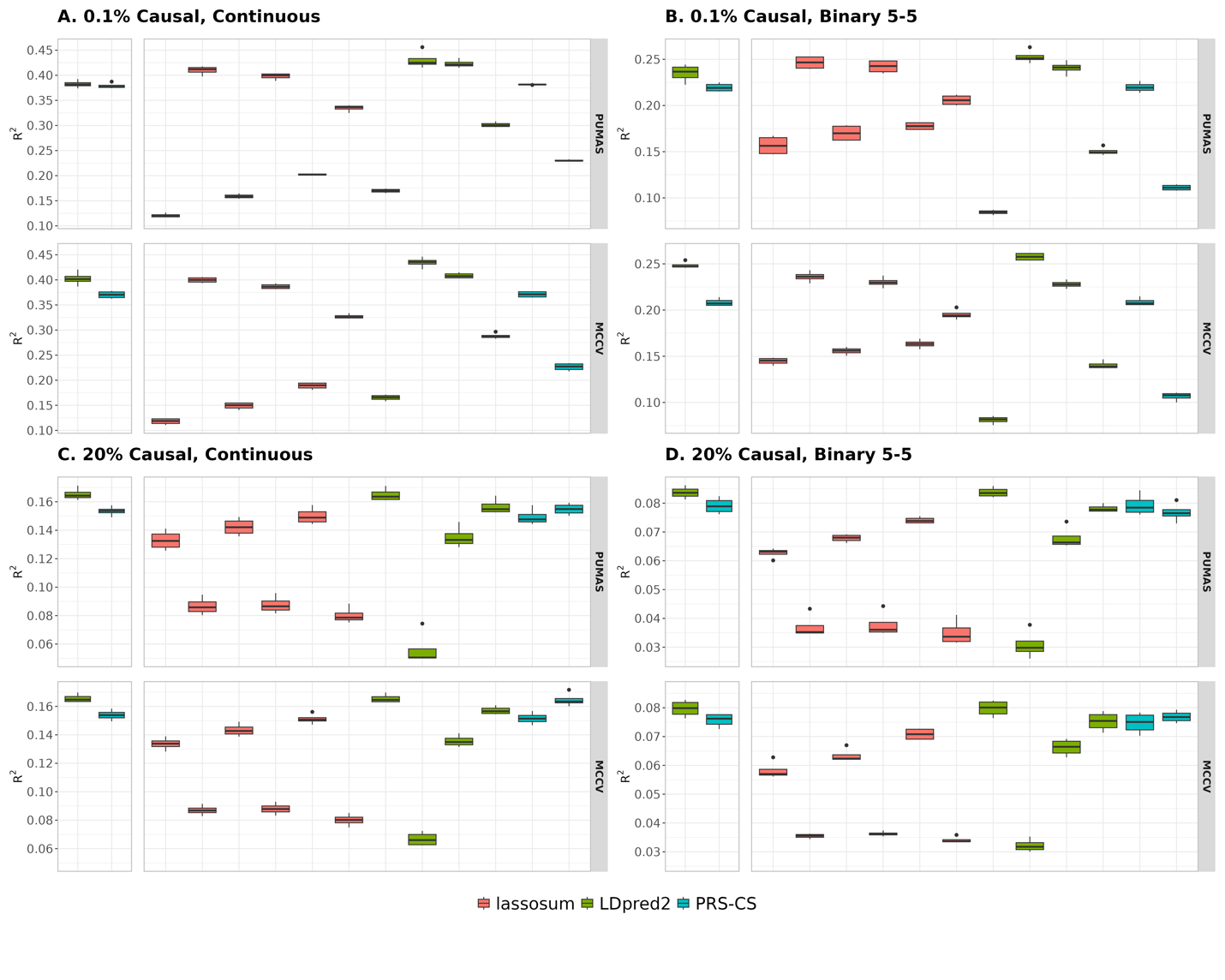

**Fig S4. Comparison of PUMAS and MCCV in UKB simulation when heritability is 0.5**. (**A** and **C**) Simulation results for quantitative traits. (**B** and **D**) Simulation results for binary traits with balanced case-control ratio. Proportion of causal variants is 0.1% in **A** and **B**, and 20% in **C** and **D**. Models that do not require fine-tuning are shown on the left side of each panel. Y-axis: predictive $R^{2}$ across 4 repeats of MCCV; X-axis (left to right): tuning-free models: LDpred2-auto (green box) and PRS-CS-auto (blue box). lassosum models (red boxes) with tuning parameter settings: s=0.2 and λ=0.005, s=0.2 and λ=0.01, s=0.5 and λ=0.005, s=0.5 and λ=0.01, s=0.9 and λ=0.005, s=0.9 and λ=0.01. LDpred2 models (green boxes): non-infinitesimal with p=0.1, non-infinitesimal with p=0.01, non-infinitesimal with p=0.001, and infinitesimal model. PRS-CS (blue boxes): $\phi$=0.01 and 0.0001.

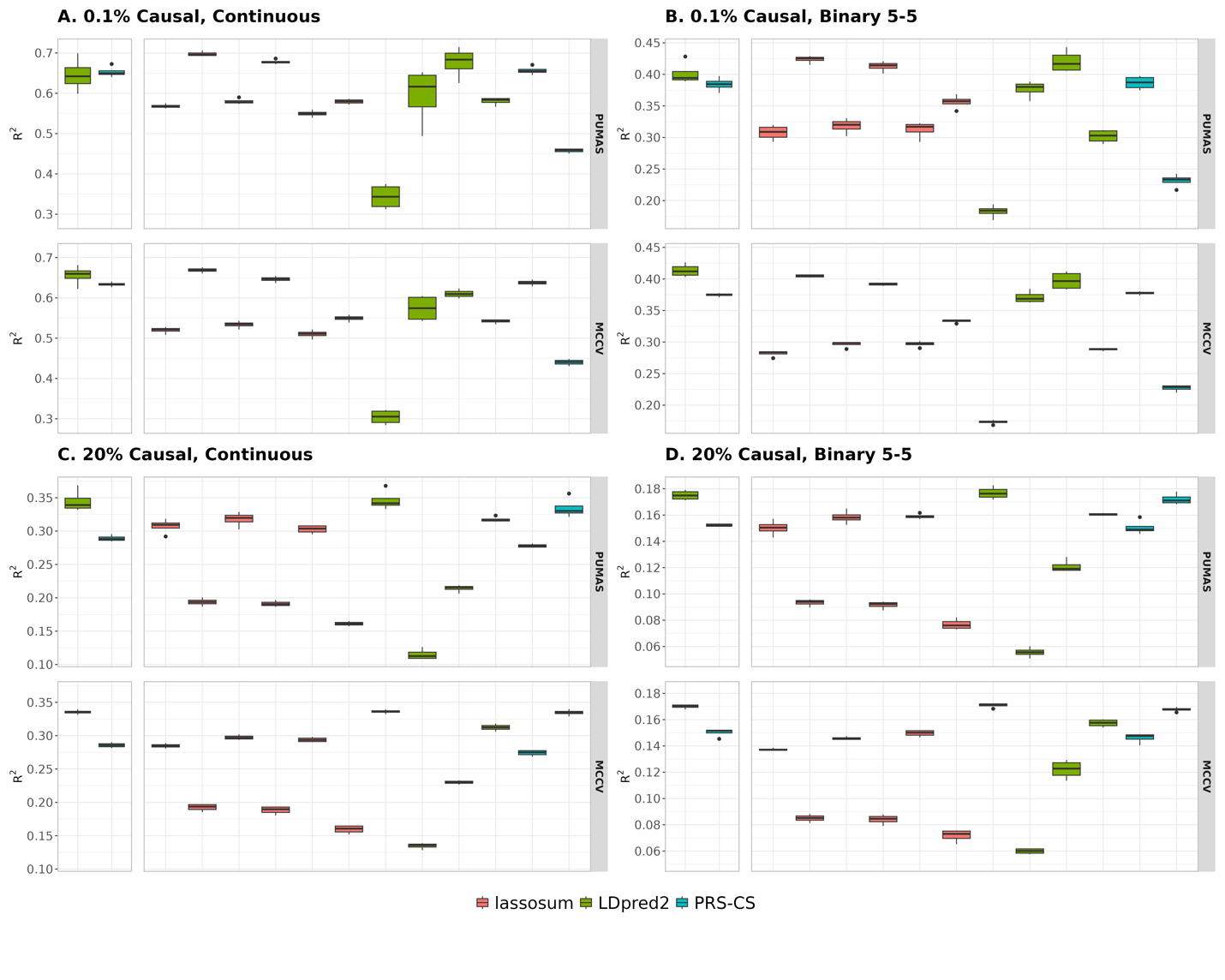

**Fig S5. Comparison of PUMAS and MCCV in UKB simulation when heritability is 0.8**. (**A** and **C**) Simulation results for quantitative traits. (**B** and **D**) Simulation results for binary traits with balanced case-control ratio. Proportion of causal variants is 0.1% in **A** and **B**, and 20% in **C** and **D**. Models that do not require fine-tuning are shown on the left side of each panel. Y-axis: predictive $R^{2}$ across 4 repeats of MCCV; X-axis (left to right): tuning-free models: LDpred2-auto (green box) and PRS-CS-auto (blue box). lassosum models (red boxes) with tuning parameter settings: s=0.2 and λ=0.005, s=0.2 and λ=0.01, s=0.5 and λ=0.005, s=0.5 and λ=0.01, s=0.9 and λ=0.005, s=0.9 and λ=0.01. LDpred2 models (green boxes): non-infinitesimal with p=0.1, non-infinitesimal with p=0.01, non-infinitesimal with p=0.001, and infinitesimal model. PRS-CS (blue boxes): $\phi$=0.01 and 0.0001.

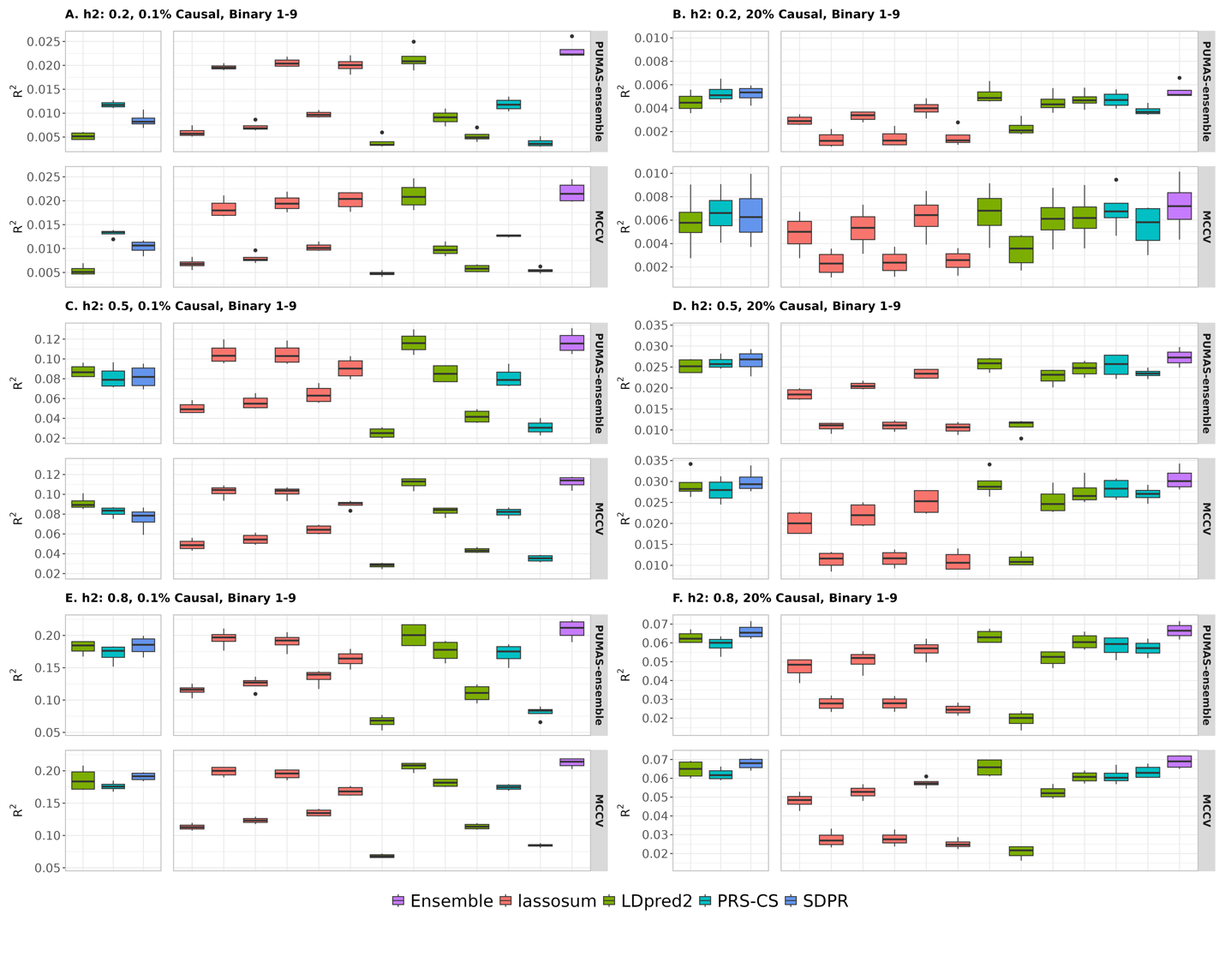

**Fig S6. Comparison of PUMAS-ensemble and MCCV in UKB binary simulation with unbalanced case-control ratio (1:9)**. (**A** and **B**) Heritability is 0.2. (**C** and **D**) Heritability is 0.5. (**E** and **F**) Heritability is 0.8. Proportion of causal variants is 0.1% in **A**, **C** and **E**, and 20% in **B**, **D** and **F**. Models that do not require fine-tuning are shown on the left side of each panel. Y-axis: predictive $R^{2}$ across 4 repeats of MCCV; X-axis (left to right): tuning-free models: LDpred2-auto (green box), PRS-CS-auto (blue box), and SDPR (dark-blue box). lassosum models (red boxes) with tuning parameter settings: s=0.2 and λ=0.005, s=0.2 and λ=0.01, s=0.5 and λ=0.005, s=0.5 and λ=0.01, s=0.9 and λ=0.005, s=0.9 and λ=0.01. LDpred2 models (green boxes): non-infinitesimal with p=0.1, non-infinitesimal with p=0.01, non-infinitesimal with p=0.001, and infinitesimal model. PRS-CS (blue boxes): $\phi$=0.01 and 0.0001. Finally, the purple box shows the results of ensemble PRS.

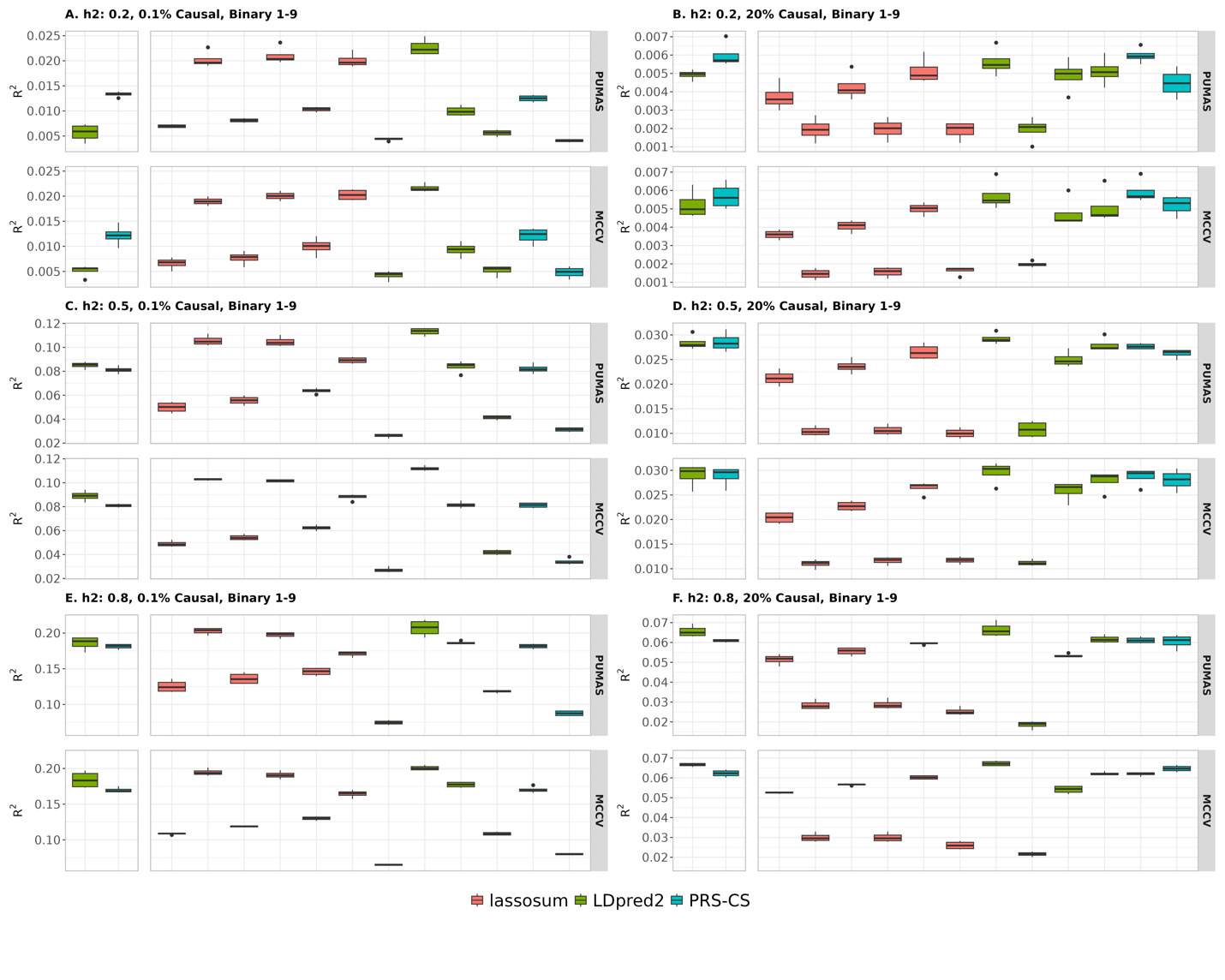

**Fig S7. Comparison of PUMAS and MCCV in UKB binary simulation with unbalanced case-control ratio (1:9)**. (**A** and **B**) Heritability is 0.2. (**C** and **D**) Heritability is 0.5. (**E** and **F**) Heritability is 0.8. Proportion of causal variants is 0.1% in **A**, **C** and **E**, and 20% in **B**, **D** and **F**. Models that do not require fine-tuning are shown on the left side of each panel. Y-axis: predictive $R^{2}$ across 4 repeats of MCCV; X-axis (left to right): tuning-free models: LDpred2-auto (green box) and PRS-CS-auto (blue box). lassosum models (red boxes) with tuning parameter settings: s=0.2 and λ=0.005, s=0.2 and λ=0.01, s=0.5 and λ=0.005, s=0.5 and λ=0.01, s=0.9 and λ=0.005, s=0.9 and λ=0.01. LDpred2 models (green boxes): non-infinitesimal with p=0.1, non-infinitesimal with p=0.01, non-infinitesimal with p=0.001, and infinitesimal model. PRS-CS (blue boxes): $\phi$=0.01 and 0.0001.

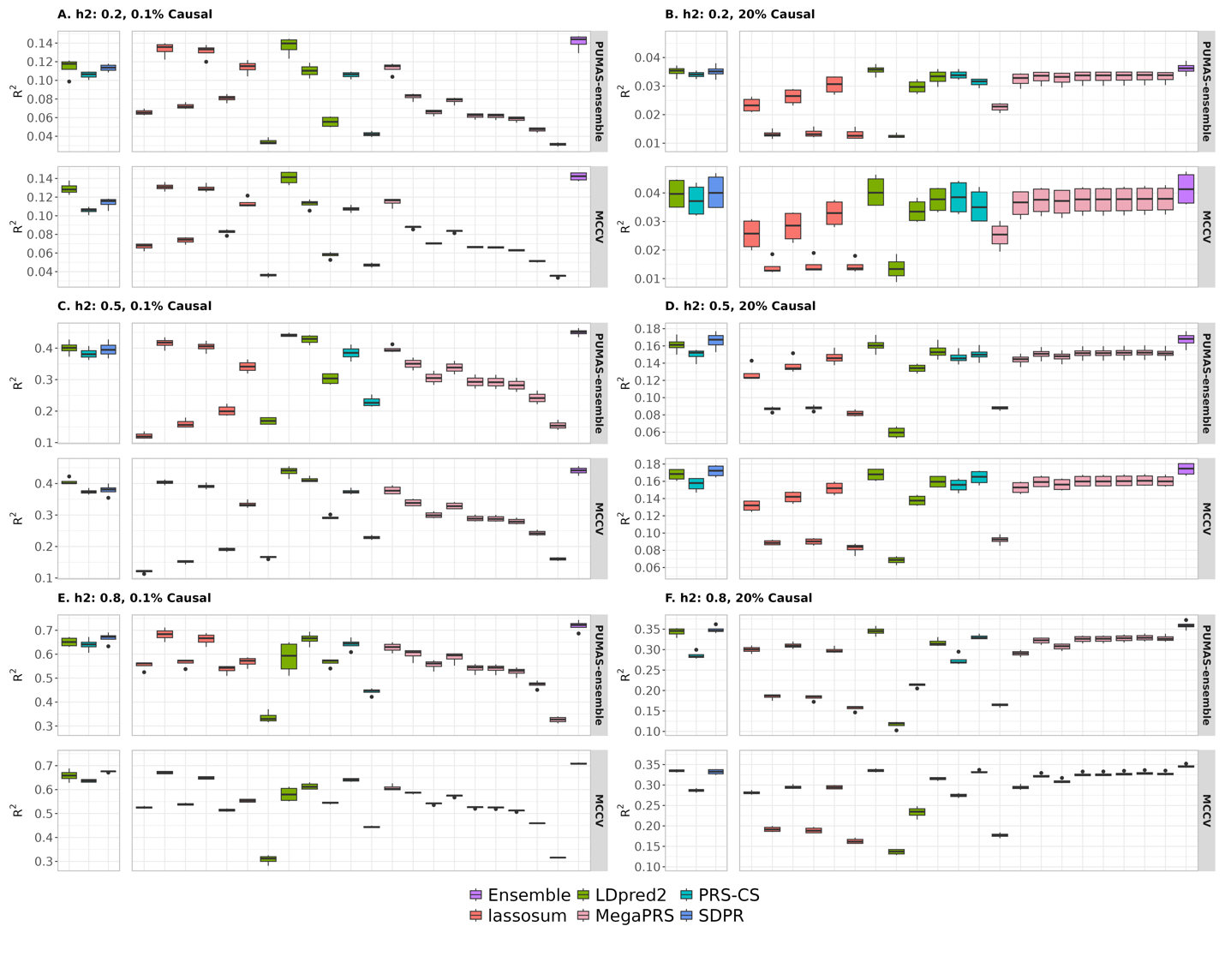

**Fig S8. Comparison of PUMAS-ensemble and MCCV in UKB quantitative simulation including MegaPRS**. (**A** and **B**) Heritability is 0.2. (**C** and **D**) Heritability is 0.5. (**E** and **F**) Heritability is 0.8. Proportion of causal variants is 0.1% in **A**, **C** and **E**, and 20% in **B**, **D** and **F**. Models that do not require fine-tuning are shown on the left side of each panel. Y-axis: predictive $R^{2}$ across 4 repeats of MCCV; X-axis (left to right): tuning-free models: LDpred2-auto (green box), PRS-CS-auto (blue box), and SDPR (dark-blue box). lassosum models (red boxes) with tuning parameter settings: s=0.2 and λ=0.005, s=0.2 and λ=0.01, s=0.5 and λ=0.005, s=0.5 and λ=0.01, s=0.9 and λ=0.005, s=0.9 and λ=0.01. LDpred2 models (green boxes): non-infinitesimal with p=0.1, non-infinitesimal with p=0.01, non-infinitesimal with p=0.001, and infinitesimal model. MegaPRS (pink boxes) with tuning parameter settings: {p1,p2,p3,p4} = {0.99,0.01,0,0}, {0.95,0.05,0,0}, {0.9,0.1,0,0}, {0.8,0.1,0.05,0.05}, {0.7,0.1,0.1,0.1}, {0.6,0.2,0.1,0.1}, {0.5,0.2,0.2,0.1}, {0.4,0.2,0.2,0.2}, {0,0,0,1}. PRS-CS (blue boxes): $\phi$=0.01 and 0.0001. Finally, the purple box shows the results of ensemble PRS.

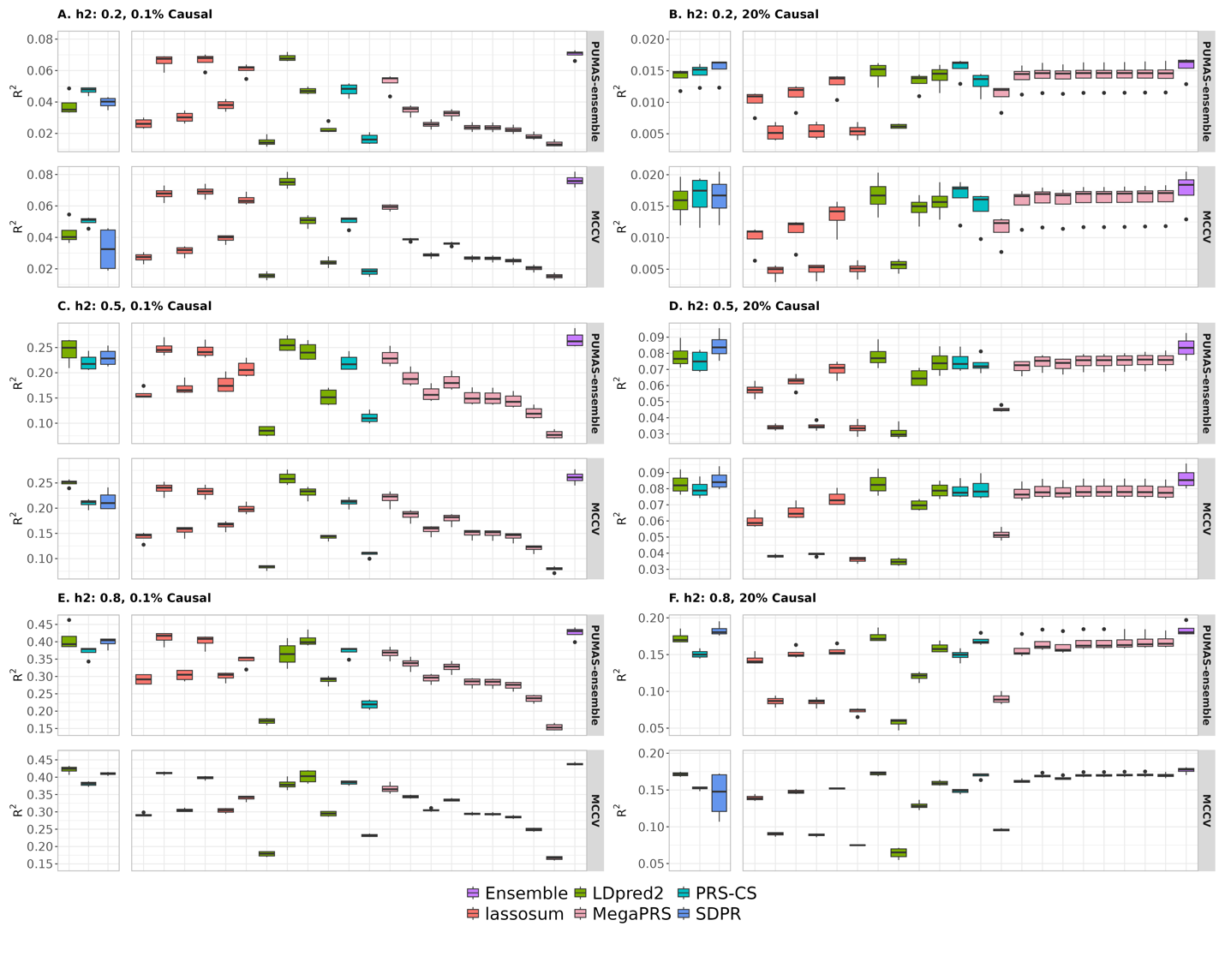

**Fig S9. Comparison of PUMAS-ensemble and MCCV in UKB binary simulation with balanced case-control ratio (5:5) including MegaPRS**. (**A** and **B**) Heritability is 0.2. (**C** and **D**) Heritability is 0.5. (**E** and **F**) Heritability is 0.8. Proportion of causal variants is 0.1% in **A**, **C** and **E**, and 20% in **B**, **D** and **F**. Models that do not require fine-tuning are shown on the left side of each panel. Y-axis: predictive $R^{2}$ across 4 repeats of MCCV; X-axis (left to right): tuning-free models: LDpred2-auto (green box), PRS-CS-auto (blue box), and SDPR (dark-blue box). lassosum models (red boxes) with tuning parameter settings: s=0.2 and λ=0.005, s=0.2 and λ=0.01, s=0.5 and λ=0.005, s=0.5 and λ=0.01, s=0.9 and λ=0.005, s=0.9 and λ=0.01. LDpred2 models (green boxes): non-infinitesimal with p=0.1, non-infinitesimal with p=0.01, non-infinitesimal with p=0.001, and infinitesimal model. MegaPRS (pink boxes) with tuning parameter settings: {p1,p2,p3,p4} = {0.99,0.01,0,0}, {0.95,0.05,0,0}, {0.9,0.1,0,0}, {0.8,0.1,0.05,0.05}, {0.7,0.1,0.1,0.1}, {0.6,0.2,0.1,0.1}, {0.5,0.2,0.2,0.1}, {0.4,0.2,0.2,0.2}, {0,0,0,1}. PRS-CS (blue boxes): $\phi$=0.01 and 0.0001. Finally, the purple box shows the results of ensemble PRS.

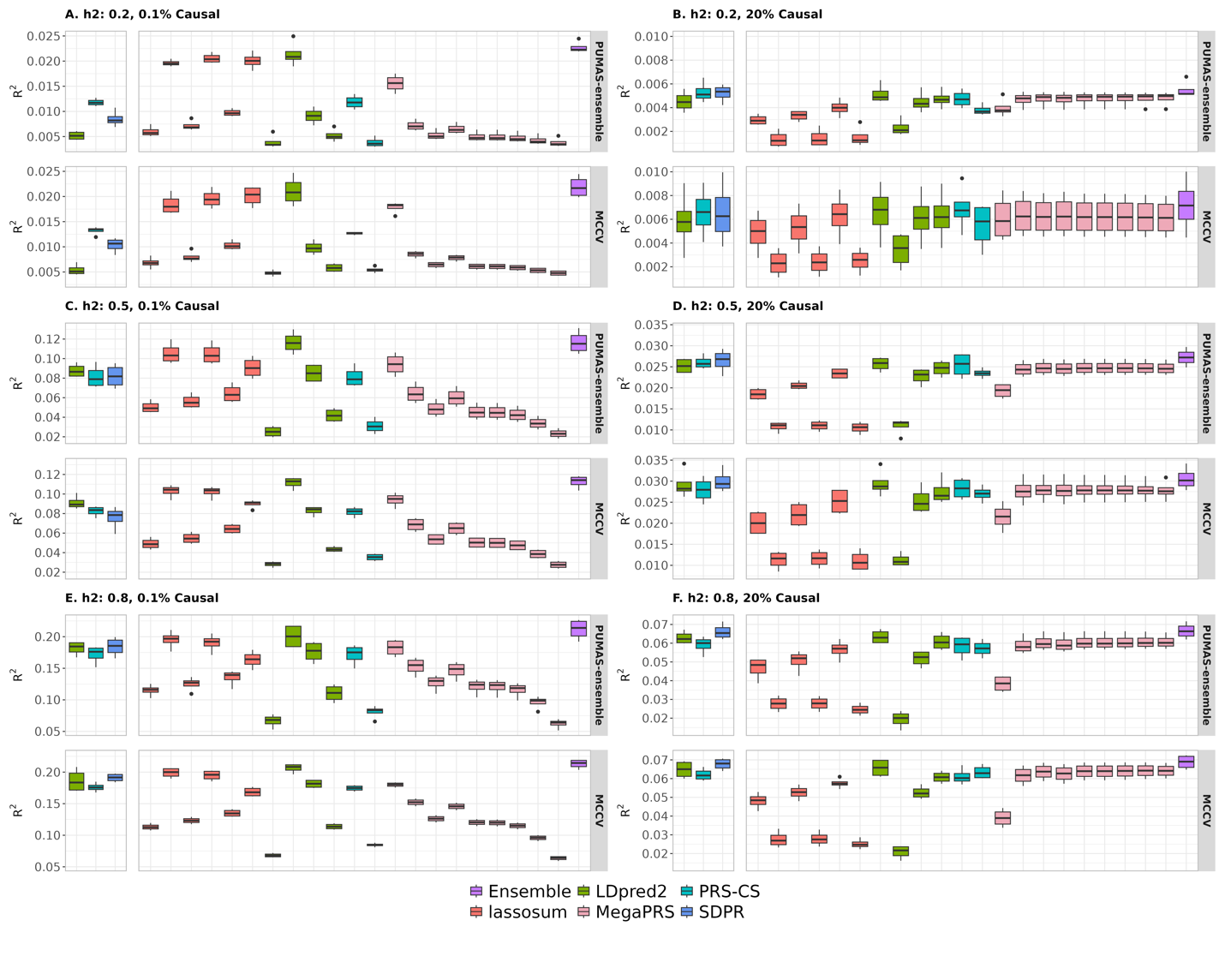

**Fig S10. Comparison of PUMAS-ensemble and MCCV in UKB binary simulation with unbalanced case-control ratio (1:9) including MegaPRS**. (**A** and **B**) Heritability is 0.2. (**C** and **D**) Heritability is 0.5. (**E** and **F**) Heritability is 0.8. Proportion of causal variants is 0.1% in **A**, **C** and **E**, and 20% in **B**, **D** and **F**. Models that do not require fine-tuning are shown on the left side of each panel. Y-axis: predictive $R^{2}$ across 4 repeats of MCCV; X-axis (left to right): tuning-free models: LDpred2-auto (green box), PRS-CS-auto (blue box), and SDPR (dark-blue box). lassosum models (red boxes) with tuning parameter settings: s=0.2 and λ=0.005, s=0.2 and λ=0.01, s=0.5 and λ=0.005, s=0.5 and λ=0.01, s=0.9 and λ=0.005, s=0.9 and λ=0.01. LDpred2 models (green boxes): non-infinitesimal with p=0.1, non-infinitesimal with p=0.01, non-infinitesimal with p=0.001, and infinitesimal model. MegaPRS (pink boxes) with tuning parameter settings: {p1,p2,p3,p4} = {0.99,0.01,0,0}, {0.95,0.05,0,0}, {0.9,0.1,0,0}, {0.8,0.1,0.05,0.05}, {0.7,0.1,0.1,0.1}, {0.6,0.2,0.1,0.1}, {0.5,0.2,0.2,0.1}, {0.4,0.2,0.2,0.2}, {0,0,0,1}. PRS-CS (blue boxes): $\phi$=0.01 and 0.0001. Finally, the purple box shows the results of ensemble PRS.

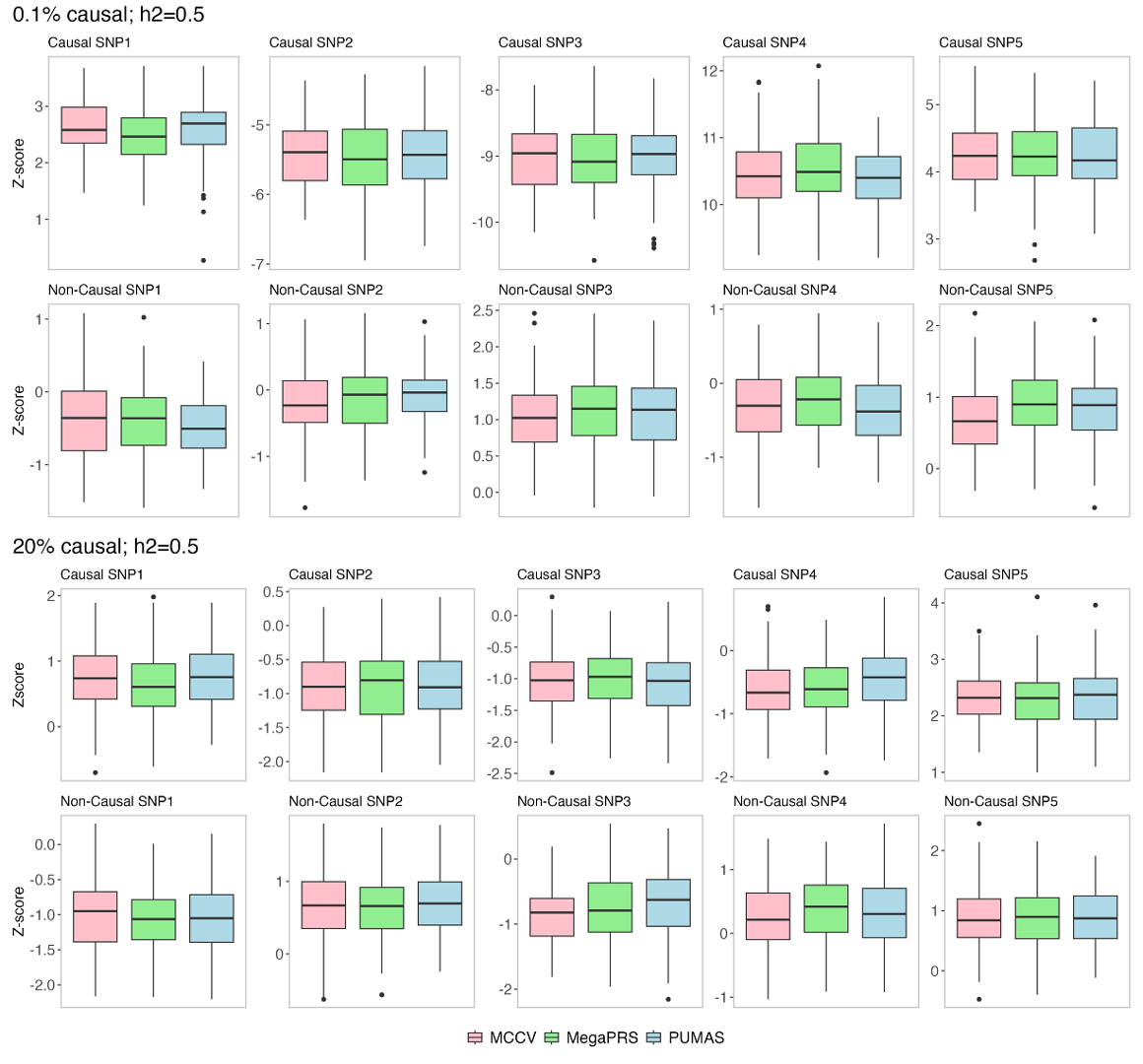

**Fig S11. Distributions of simulated Z scores from PUMAS, MegaPRS, and MCCV in simulations**. UKB simulation data (N=100,000) were used to simulate quantitative phenotype. Three set s of linear regression Z scores from 75,000 samples were simulated by PUMAS and MegaPRS and calculated by MCCV. Results are shown for 5 randomly selected causal and non-causal variants in each setting, with 100 replications.

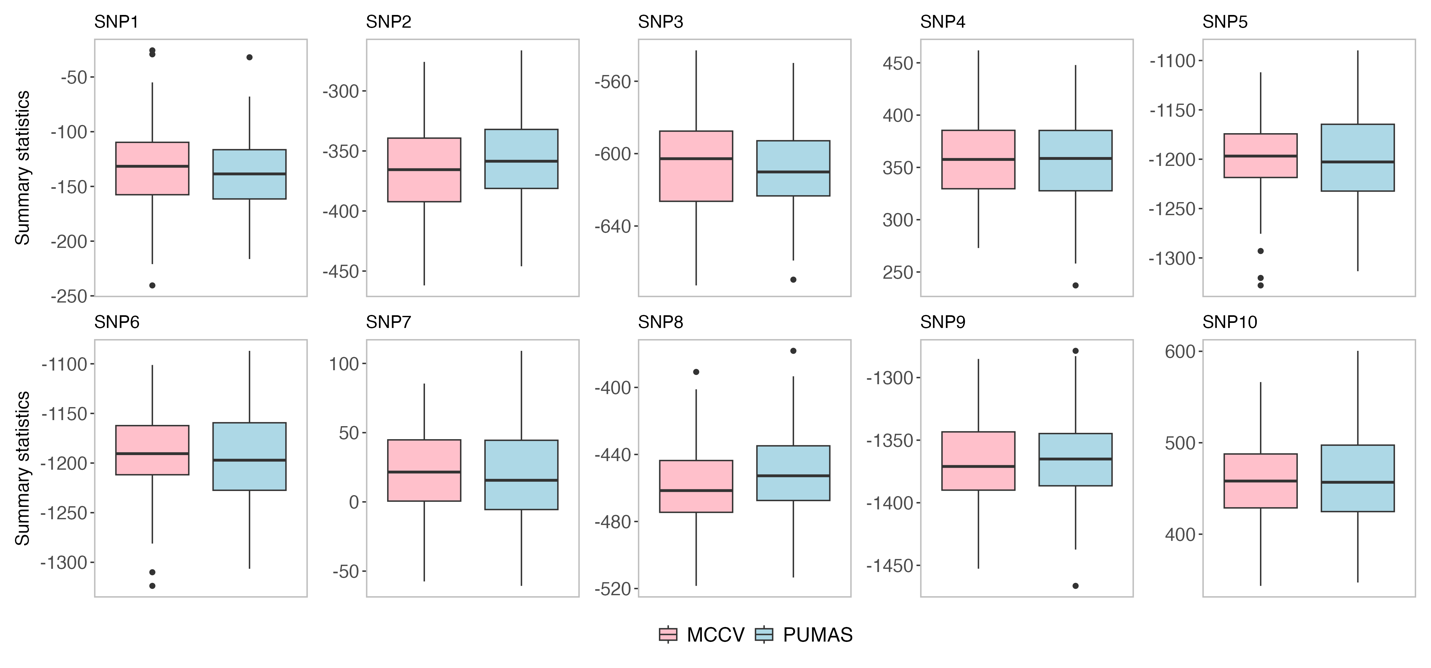
 **Fig S12. Distributions of simulated summary statistics from PUMAS and MCCV in extremely sparse simulation setting**. UKB simulation data (N=100,000) were used to simulate 10 causal SNPs that together explain 10% of trait heritability. Results are shown for each causal variant from 100 repeats.

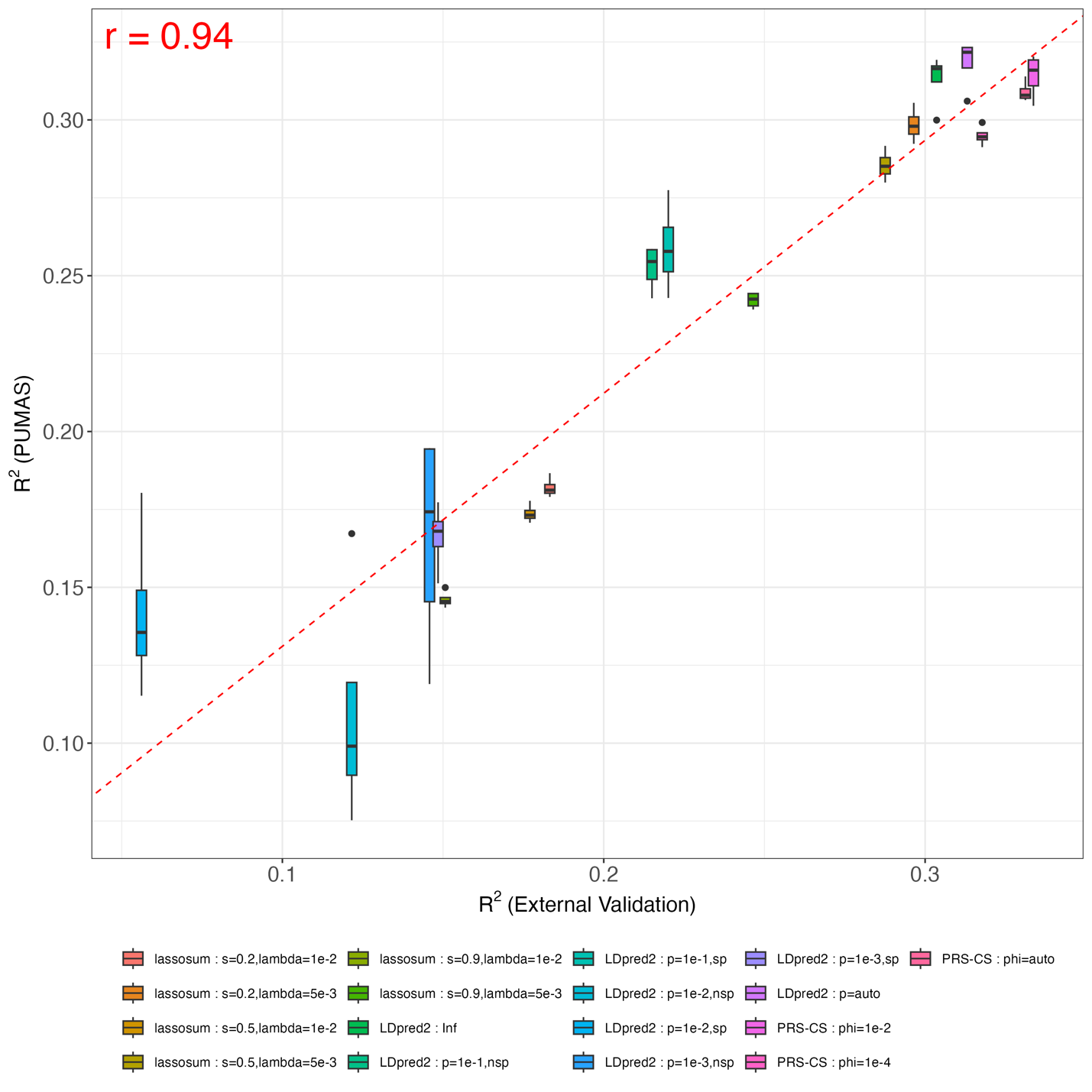

**Fig S13. Comparing PUMAS result for Height with external validation in UKB**. Y-axis: predictive $R^{2}$ across 4-fold replications from PUMAS; X-axis: predictive $R^{2}$ evaluated by external validation on the holdout dataset. The dashed red line is fitted regression line between averaged PUMAS $R^{2}$ and PRS $R^{2}$. Pearson correlation between averaged PUMAS $R^{2}$ and PRS $R^{2}$ is shown in the plot.

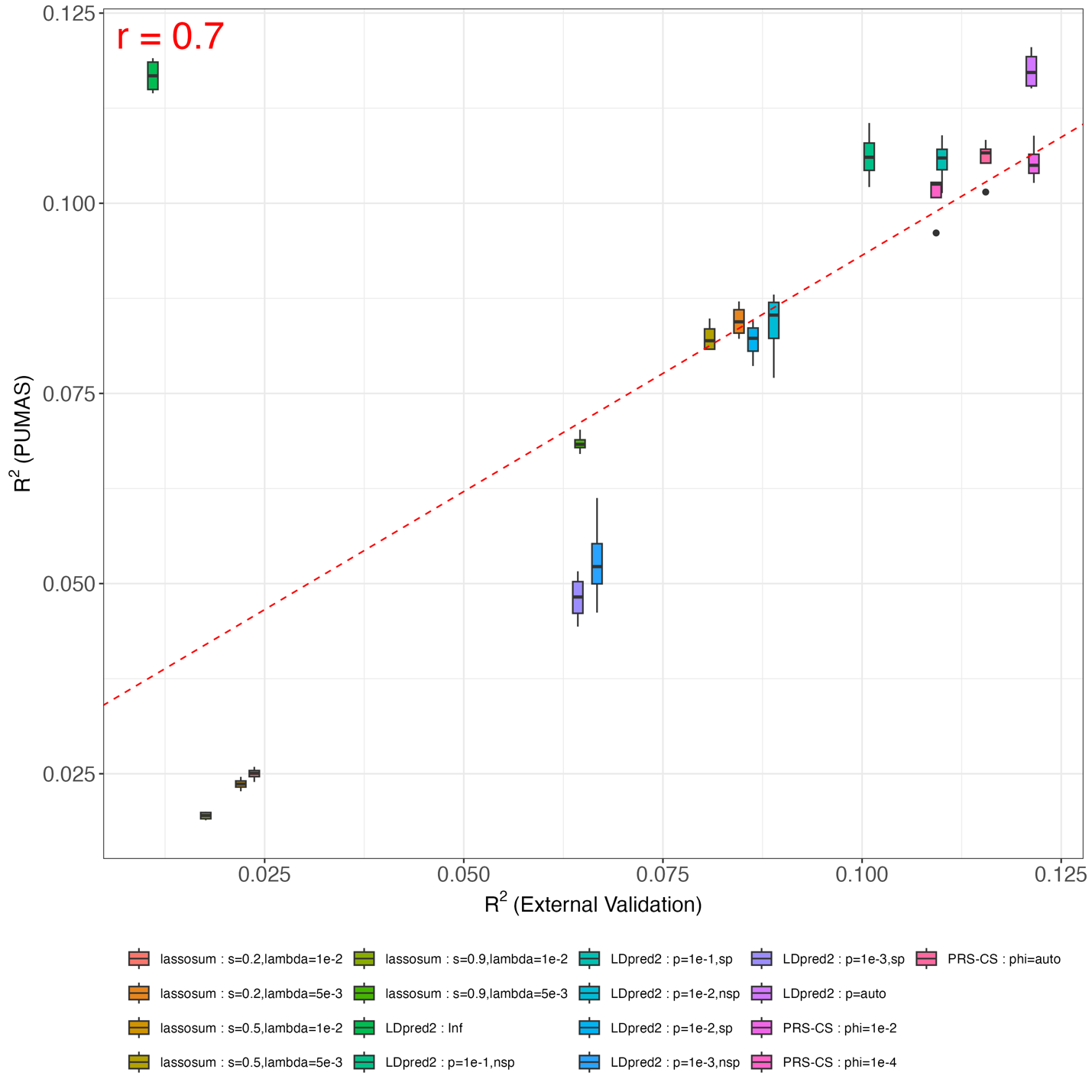

**Fig S14. Comparing PUMAS result for BMI with external validation in UKB**. Y-axis: predictive $R^{2}$ across 4-fold replications from PUMAS; X-axis: predictive $R^{2}$ evaluated by external validation on the holdout dataset. The dashed red line is fitted regression line between averaged PUMAS $R^{2}$ and PRS $R^{2}$. Pearson correlation between averaged PUMAS $R^{2}$ and PRS $R^{2}$ is shown in the plot.

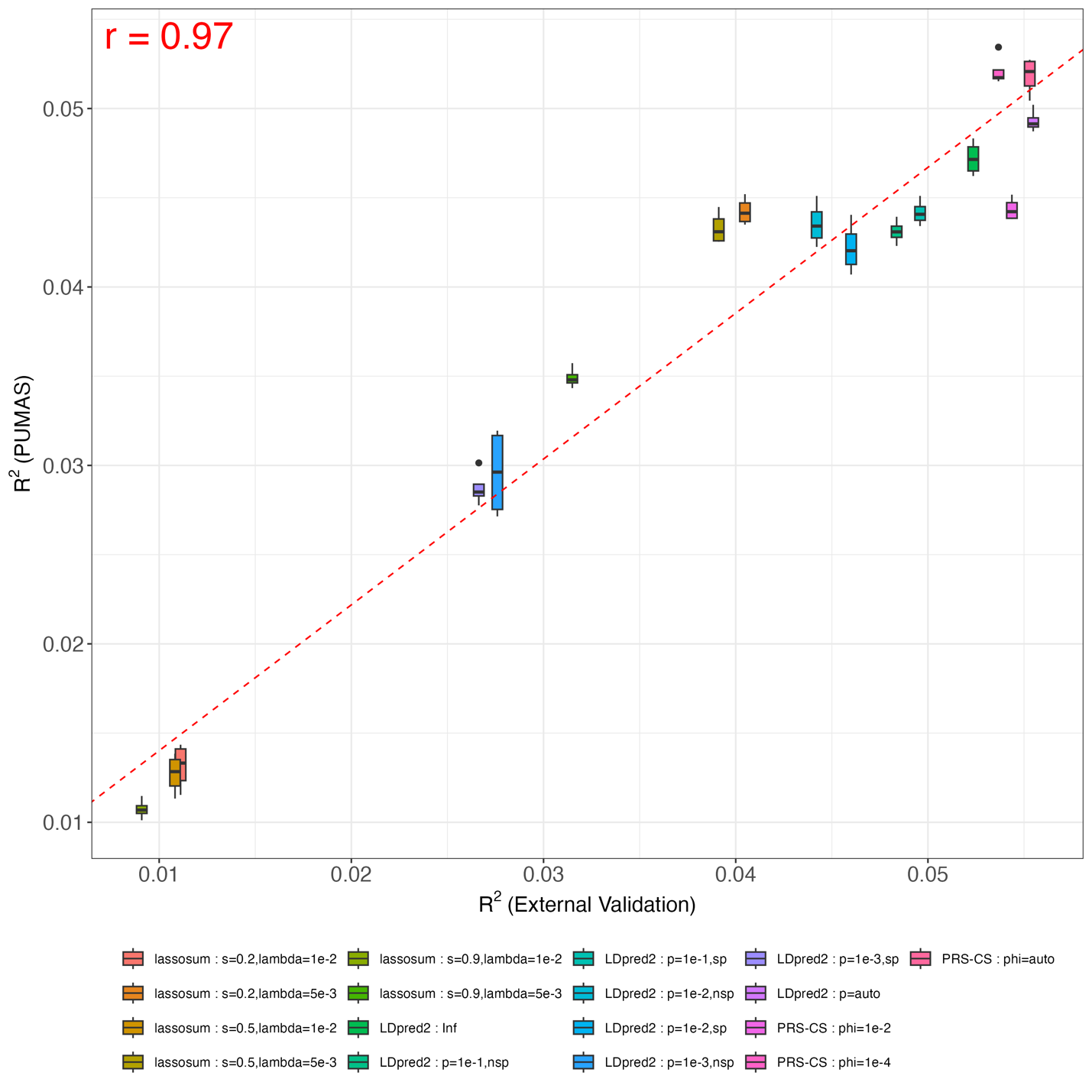

**Fig S15. Comparing PUMAS result for SBP with external validation in UKB**. Y-axis: predictive $R^{2}$ across 4-fold replications from PUMAS; X-axis: predictive $R^{2}$ evaluated by external validation on the holdout dataset. The dashed red line is fitted regression line between averaged PUMAS $R^{2}$ and PRS $R^{2}$. Pearson correlation between averaged PUMAS $R^{2}$ and PRS $R^{2}$ is shown in the plot.

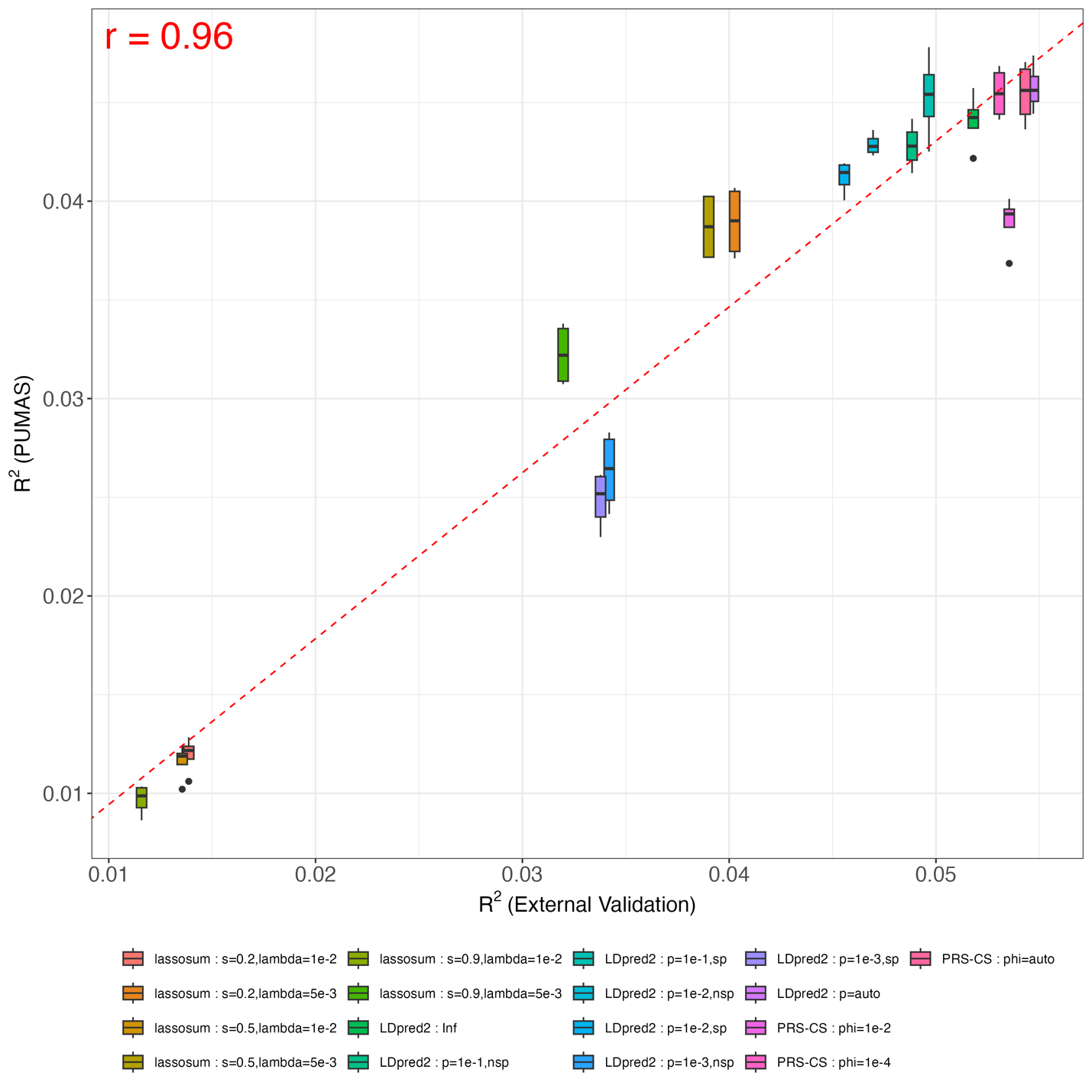

**Fig S16. Comparing PUMAS result for DBP with external validation in UKB**. Y-axis: predictive $R^{2}$ across 4-fold replications from PUMAS; X-axis: predictive $R^{2}$ evaluated by external validation on the holdout dataset. The dashed red line is fitted regression line between averaged PUMAS $R^{2}$ and PRS $R^{2}$. Pearson correlation between averaged PUMAS $R^{2}$ and PRS $R^{2}$ is shown in the plot.

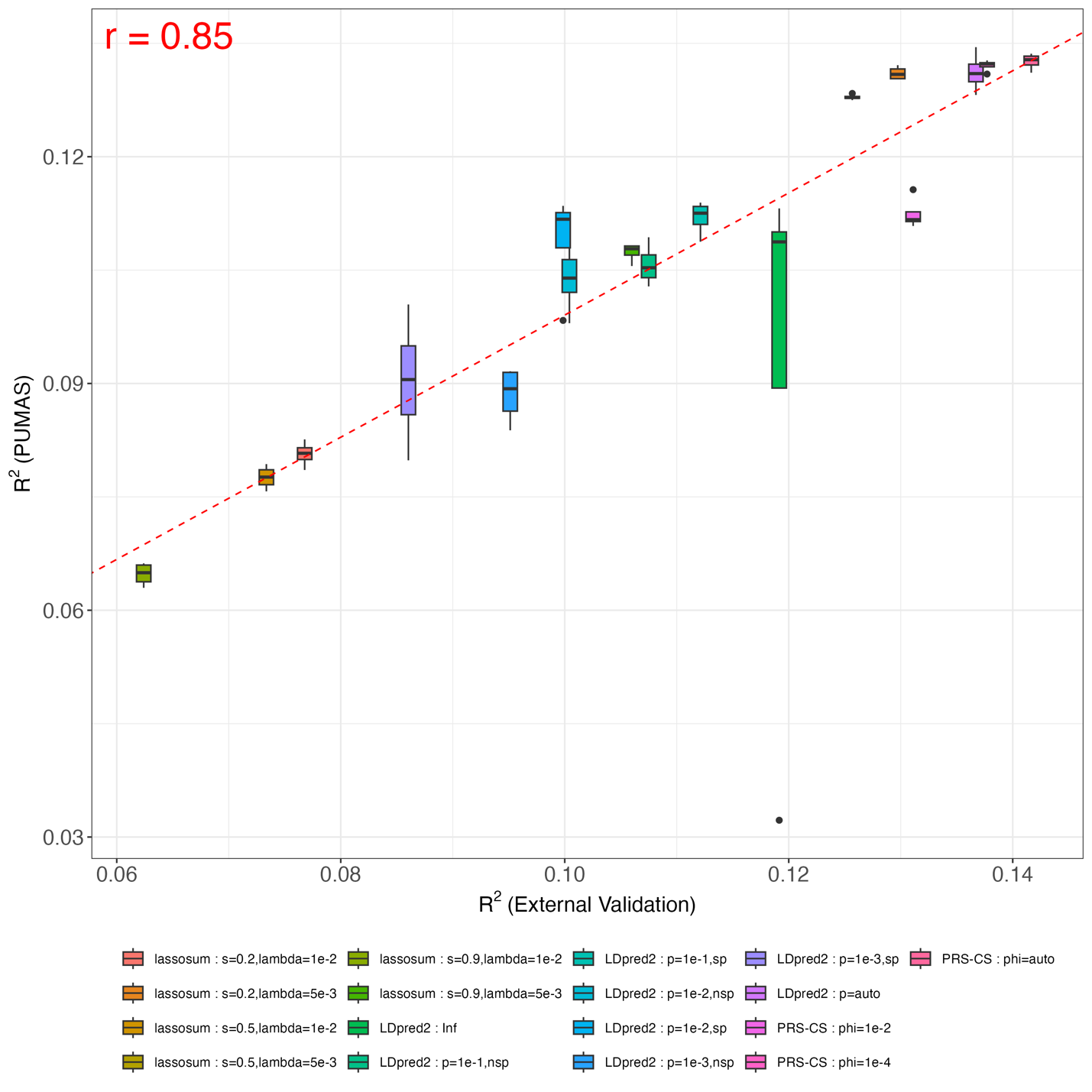

**Fig S17. Comparing PUMAS result for RBCC with external validation in UKB**. Y-axis: predictive $R^{2}$ across 4-fold replications from PUMAS; X-axis: predictive $R^{2}$ evaluated by external validation on the holdout dataset. The dashed red line is fitted regression line between averaged PUMAS $R^{2}$ and PRS $R^{2}$. Pearson correlation between averaged PUMAS $R^{2}$ and PRS $R^{2}$ is shown in the plot.

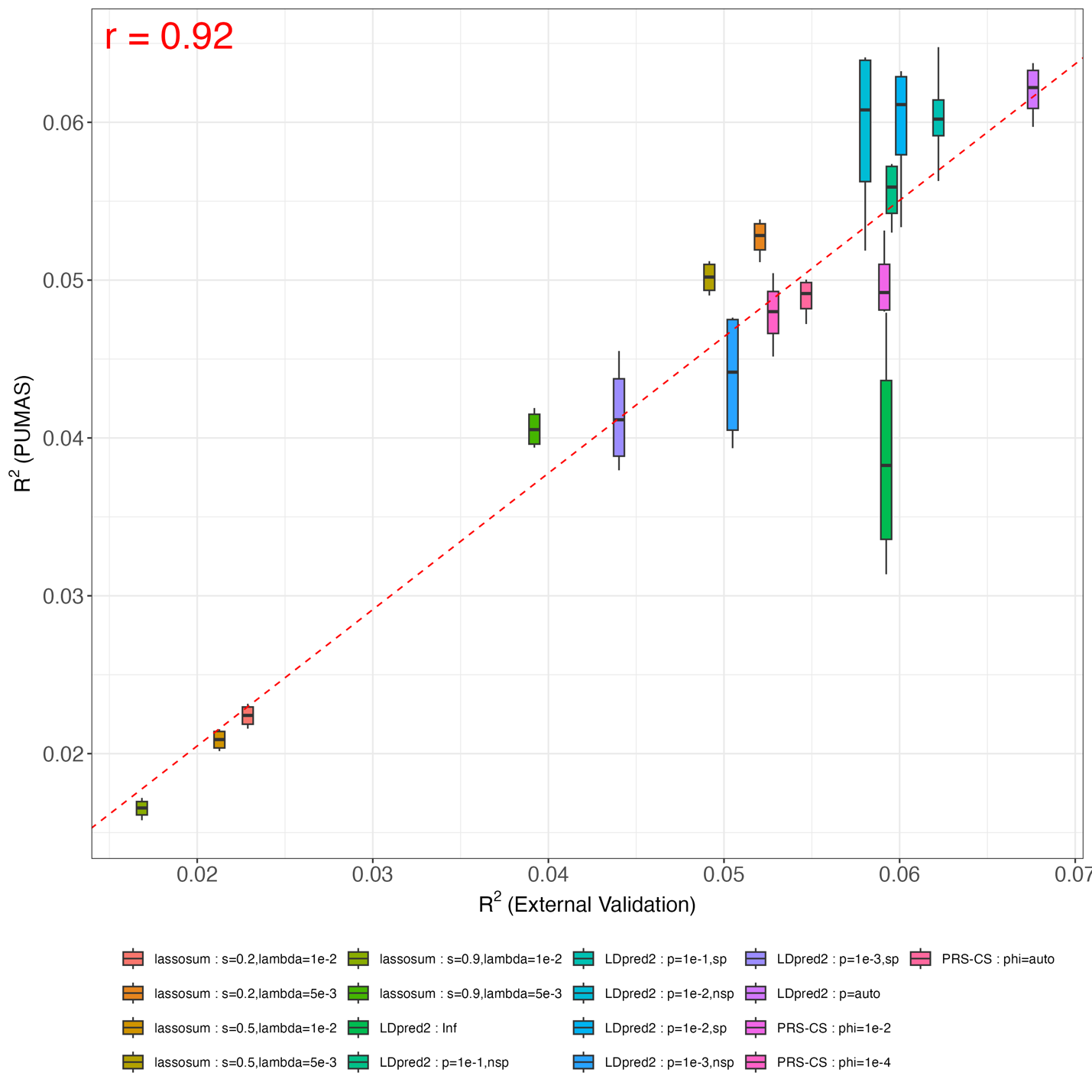

**Fig S18. Comparing PUMAS result for WBCC with external validation in UKB**. Y-axis: predictive $R^{2}$ across 4-fold replications from PUMAS; X-axis: predictive $R^{2}$ evaluated by external validation on the holdout dataset. The dashed red line is fitted regression line between averaged PUMAS $R^{2}$ and PRS $R^{2}$. Pearson correlation between averaged PUMAS $R^{2}$ and PRS $R^{2}$ is shown in the plot.

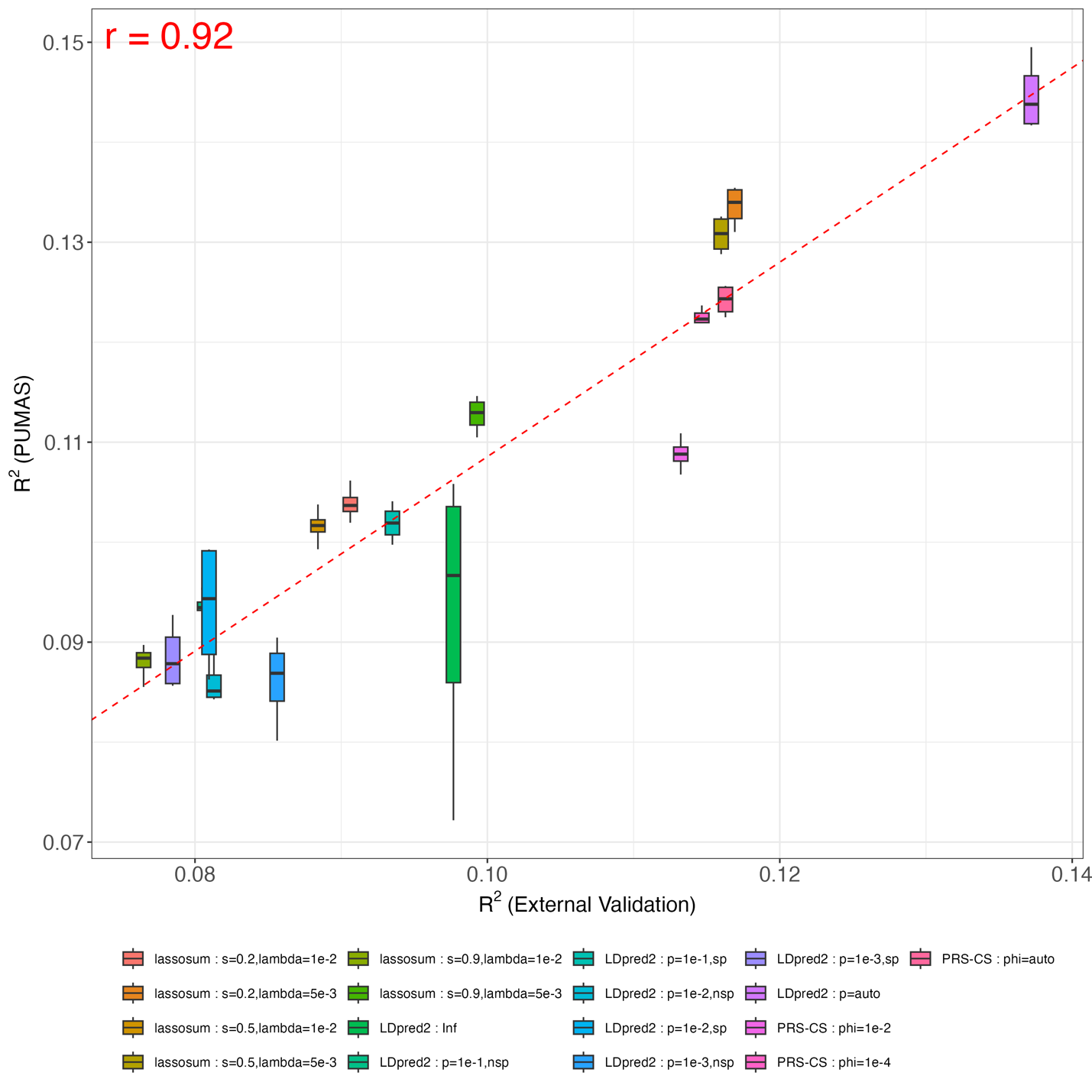

**Fig S19. Comparing PUMAS result for MCH with external validation in UKB**. Y-axis: predictive $R^{2}$ across 4-fold replications from PUMAS; X-axis: predictive $R^{2}$ evaluated by external validation on the holdout dataset. The dashed red line is fitted regression line between averaged PUMAS $R^{2}$ and PRS $R^{2}$. Pearson correlation between averaged PUMAS $R^{2}$ and PRS $R^{2}$ is shown in the plot.

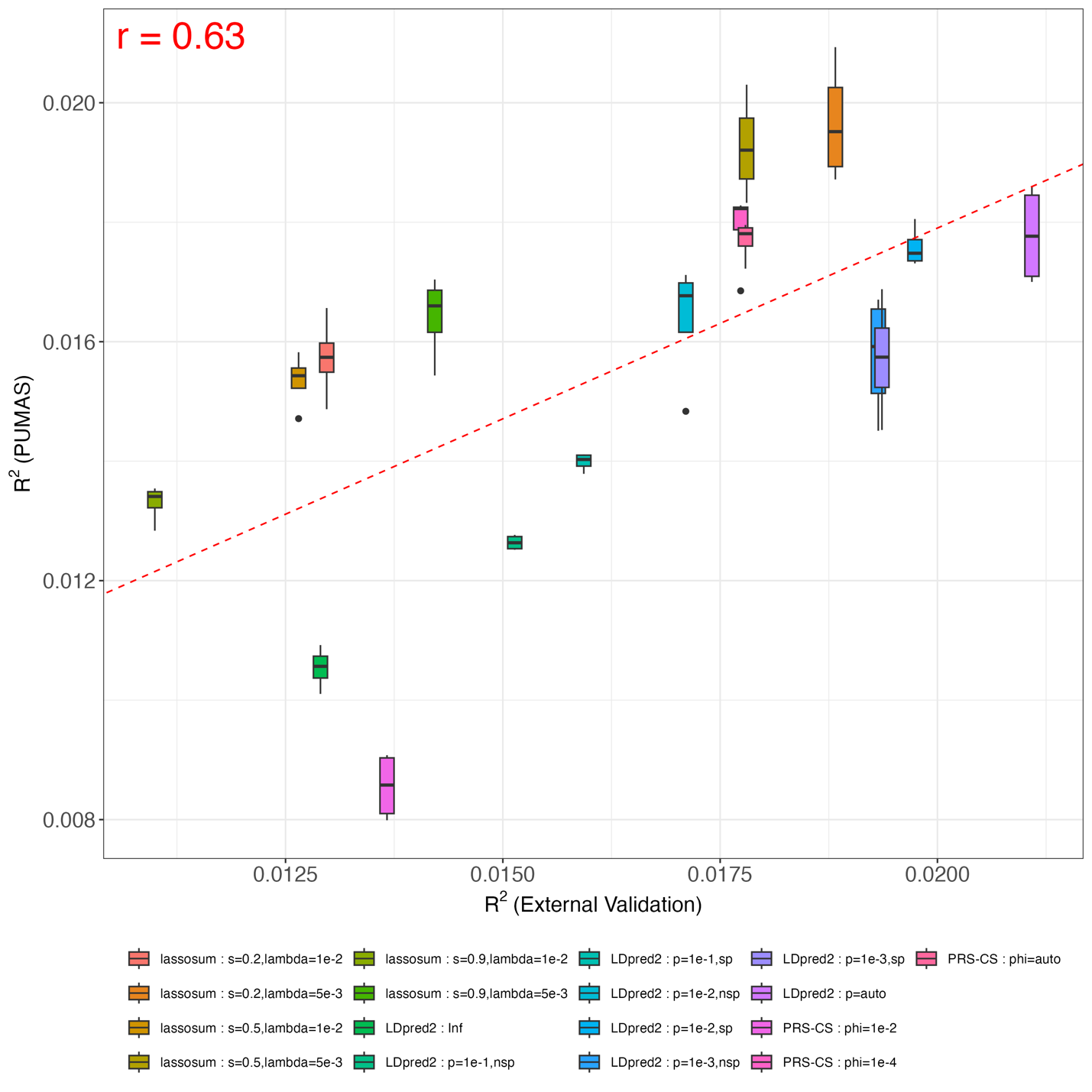

**Fig S20. Comparing PUMAS result for MCHC with external validation in UKB**. Y-axis: predictive $R^{2}$ across 4-fold replications from PUMAS; X-axis: predictive $R^{2}$ evaluated by external validation on the holdout dataset. The dashed red line is fitted regression line between averaged PUMAS $R^{2}$ and PRS $R^{2}$. Pearson correlation between averaged PUMAS $R^{2}$ and PRS $R^{2}$ is shown in the plot.

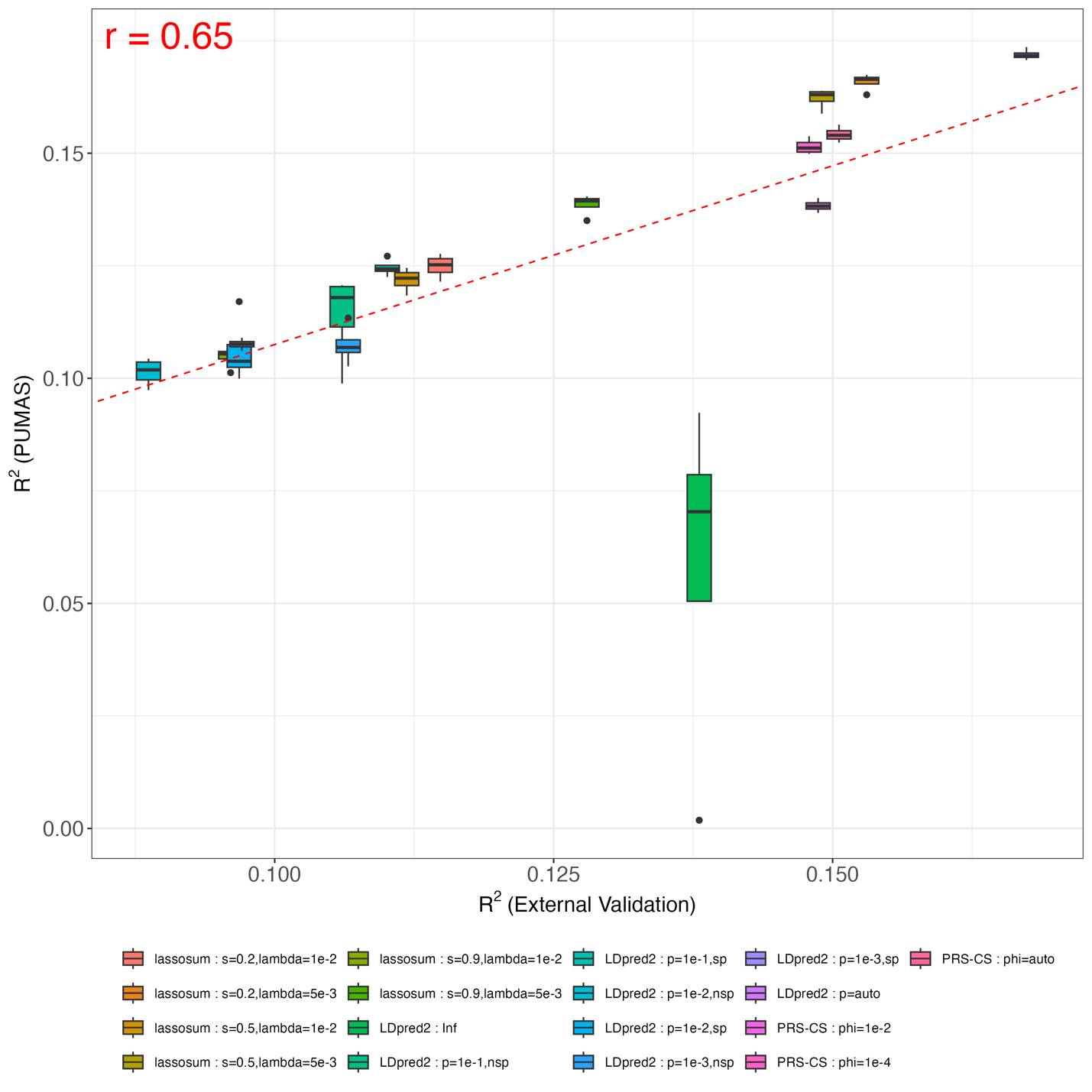

**Fig S21. Comparing PUMAS result for MCV with external validation in UKB**. Y-axis: predictive $R^{2}$ across 4-fold replications from PUMAS; X-axis: predictive $R^{2}$ evaluated by external validation on the holdout dataset. The dashed red line is fitted regression line between averaged PUMAS $R^{2}$ and PRS $R^{2}$. Pearson correlation between averaged PUMAS $R^{2}$ and PRS $R^{2}$ is shown in the plot.

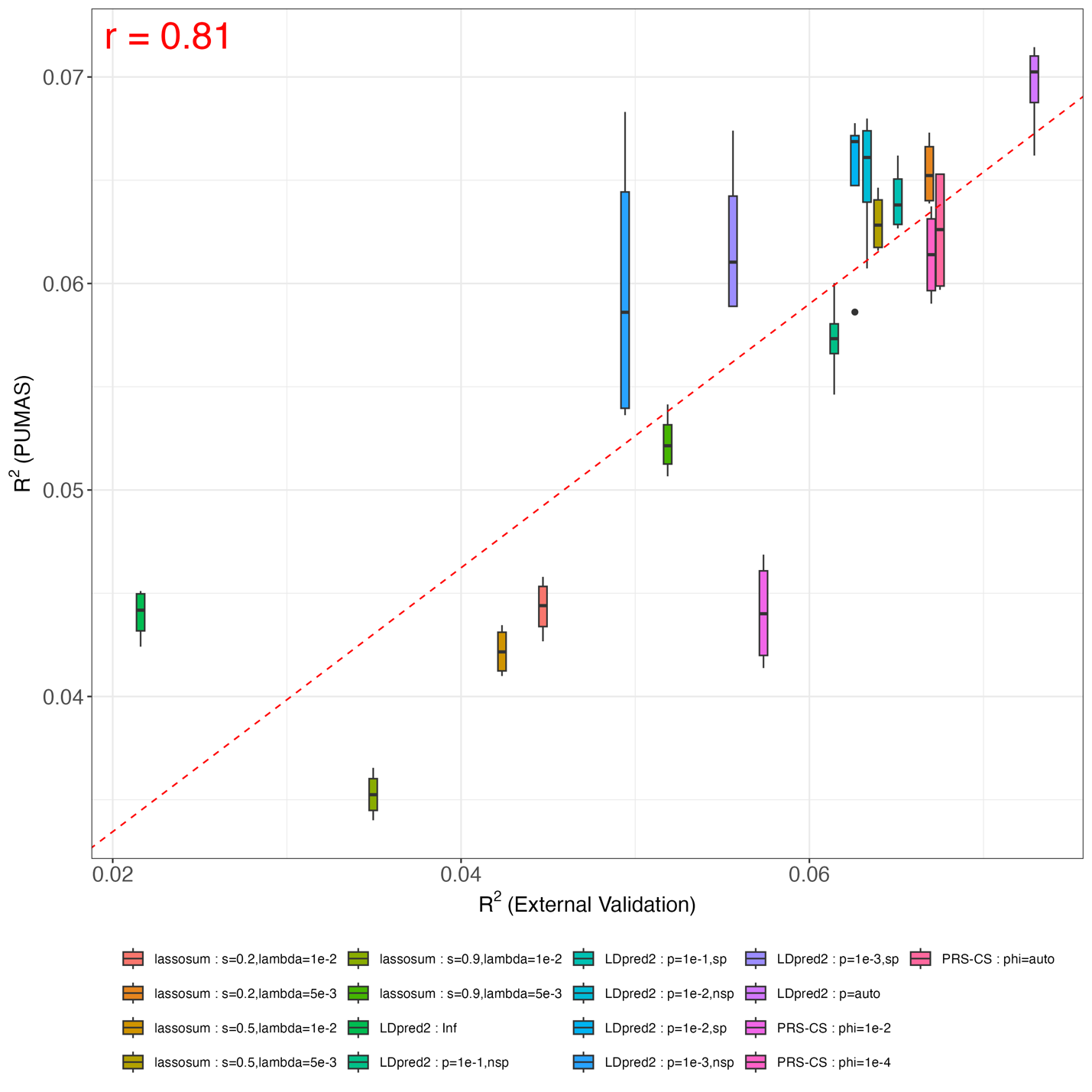

**Fig S22. Comparing PUMAS result for MC with external validation in UKB**. Y-axis: predictive $R^{2}$ across 4-fold replications from PUMAS; X-axis: predictive $R^{2}$ evaluated by external validation on the holdout dataset. The dashed red line is fitted regression line between averaged PUMAS $R^{2}$ and PRS $R^{2}$. Pearson correlation between averaged PUMAS $R^{2}$ and PRS $R^{2}$ is shown in the plot.

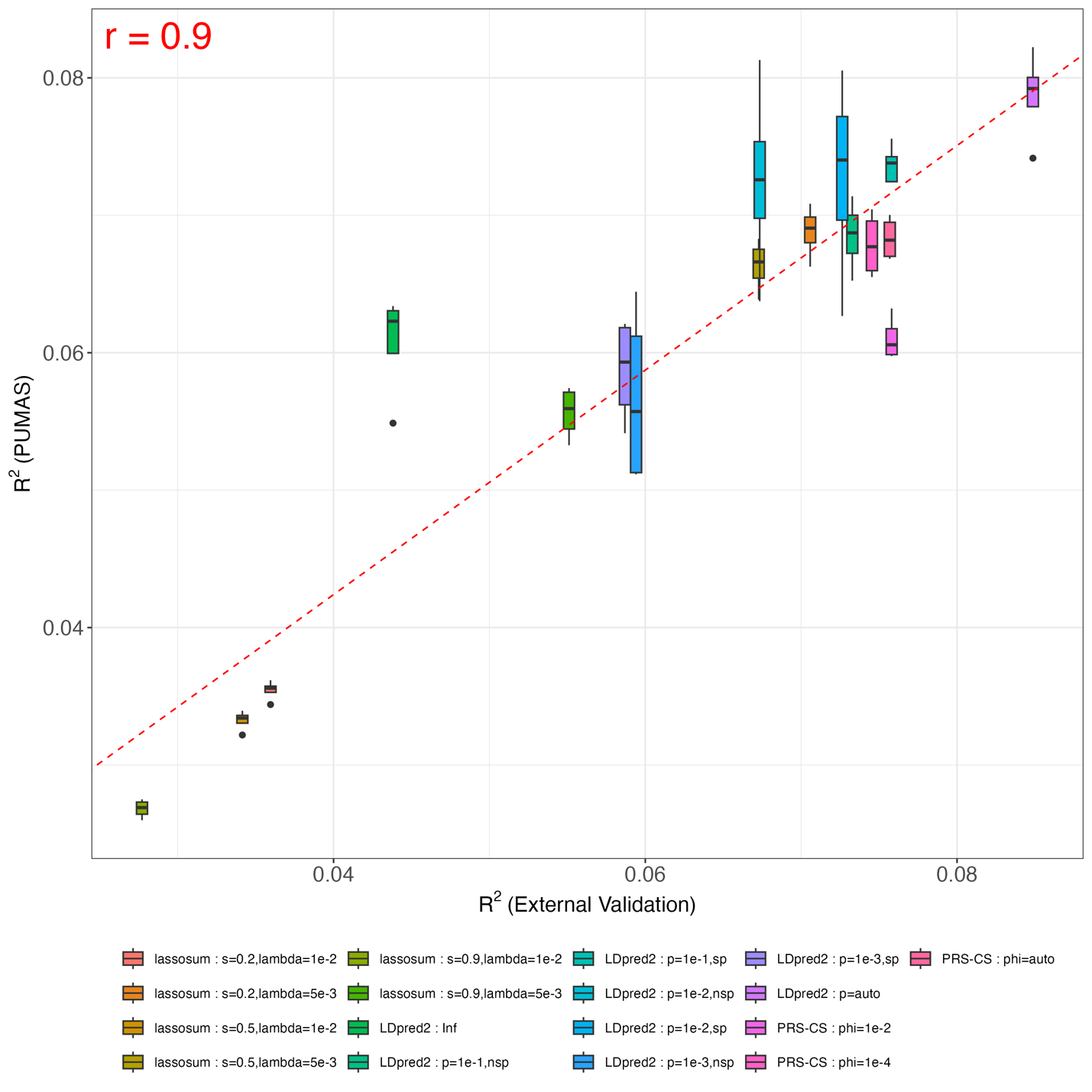

**Fig S23. Comparing PUMAS result for NC with external validation in UKB**. Y-axis: predictive $R^{2}$ across 4-fold replications from PUMAS; X-axis: predictive $R^{2}$ evaluated by external validation on the holdout dataset. The dashed red line is fitted regression line between averaged PUMAS $R^{2}$ and PRS $R^{2}$. Pearson correlation between averaged PUMAS $R^{2}$ and PRS $R^{2}$ is shown in the plot.

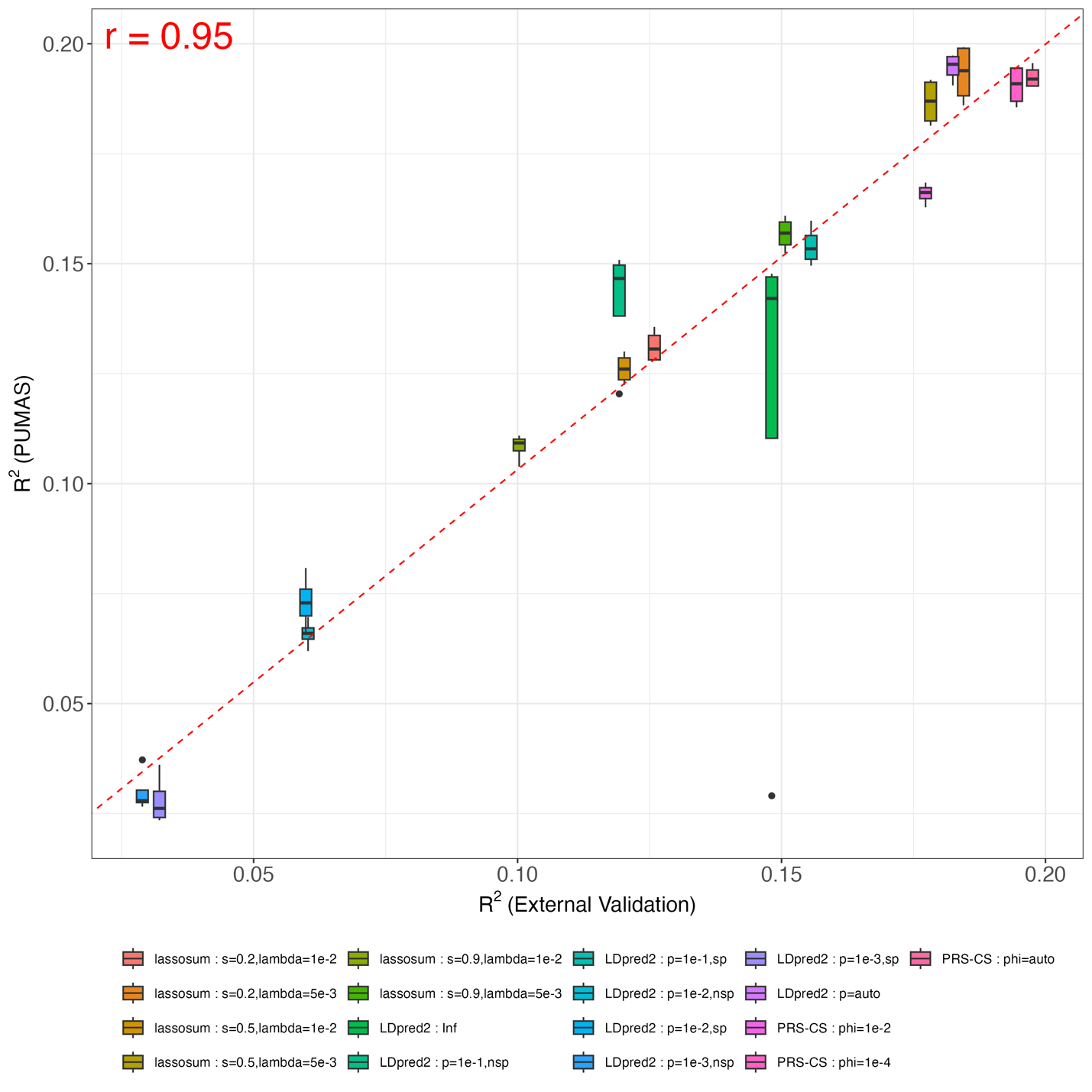

**Fig S24. Comparing PUMAS result for PC with external validation in UKB**. Y-axis: predictive $R^{2}$ across 4-fold replications from PUMAS; X-axis: predictive $R^{2}$ evaluated by external validation on the holdout dataset. The dashed red line is fitted regression line between averaged PUMAS $R^{2}$ and PRS $R^{2}$. Pearson correlation between averaged PUMAS $R^{2}$ and PRS $R^{2}$ is shown in the plot.

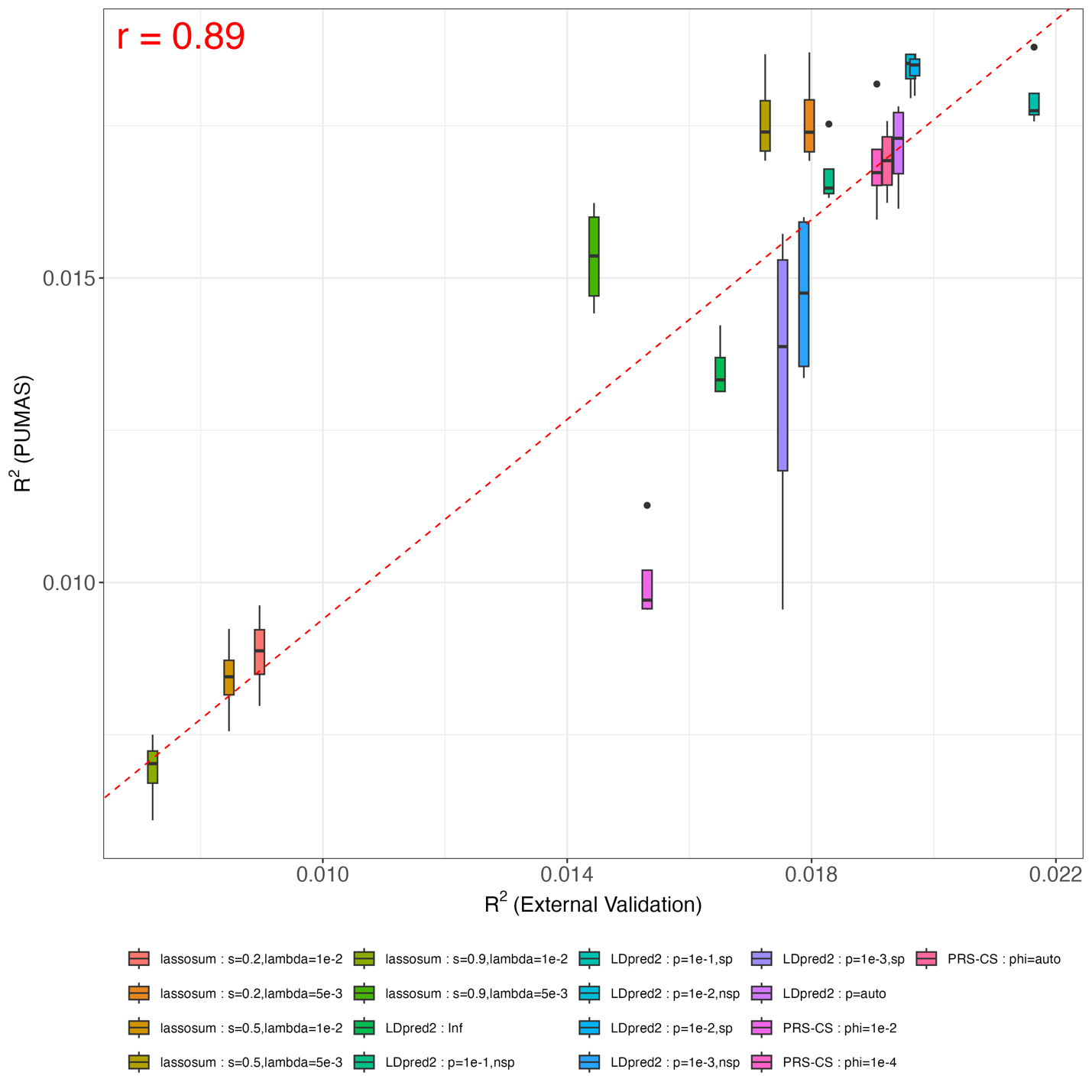

**Fig S25. Comparing PUMAS result for LC with external validation in UKB**. Y-axis: predictive $R^{2}$ across 4-fold replications from PUMAS; X-axis: predictive $R^{2}$ evaluated by external validation on the holdout dataset. The dashed red line is fitted regression line between averaged PUMAS $R^{2}$ and PRS $R^{2}$. Pearson correlation between averaged PUMAS $R^{2}$ and PRS $R^{2}$ is shown in the plot.

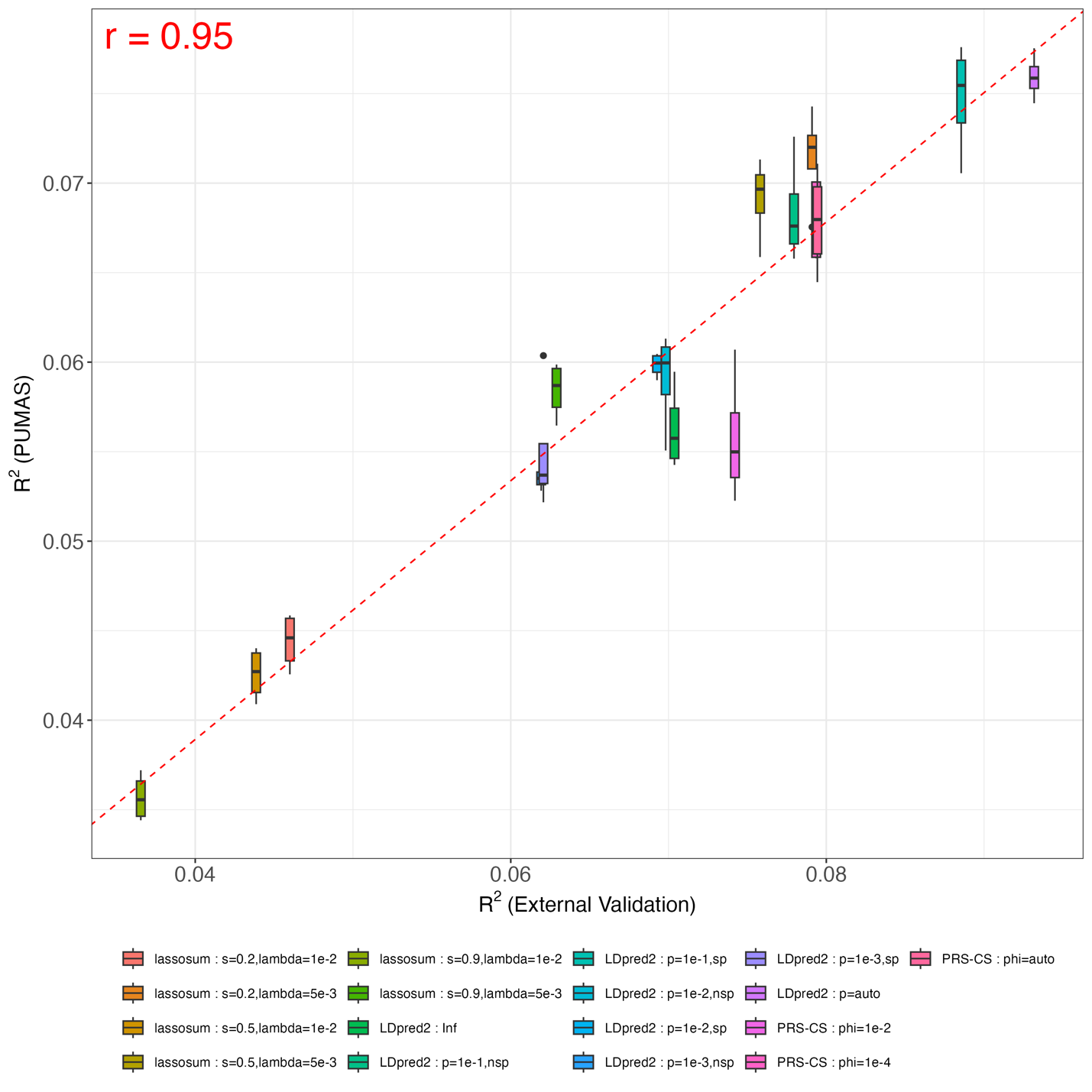

**Fig S26. Comparing PUMAS result for EC with external validation in UKB**. Y-axis: predictive $R^{2}$ across 4-fold replications from PUMAS; X-axis: predictive $R^{2}$ evaluated by external validation on the holdout dataset. The dashed red line is fitted regression line between averaged PUMAS $R^{2}$ and PRS $R^{2}$. Pearson correlation between averaged PUMAS $R^{2}$ and PRS $R^{2}$ is shown in the plot.

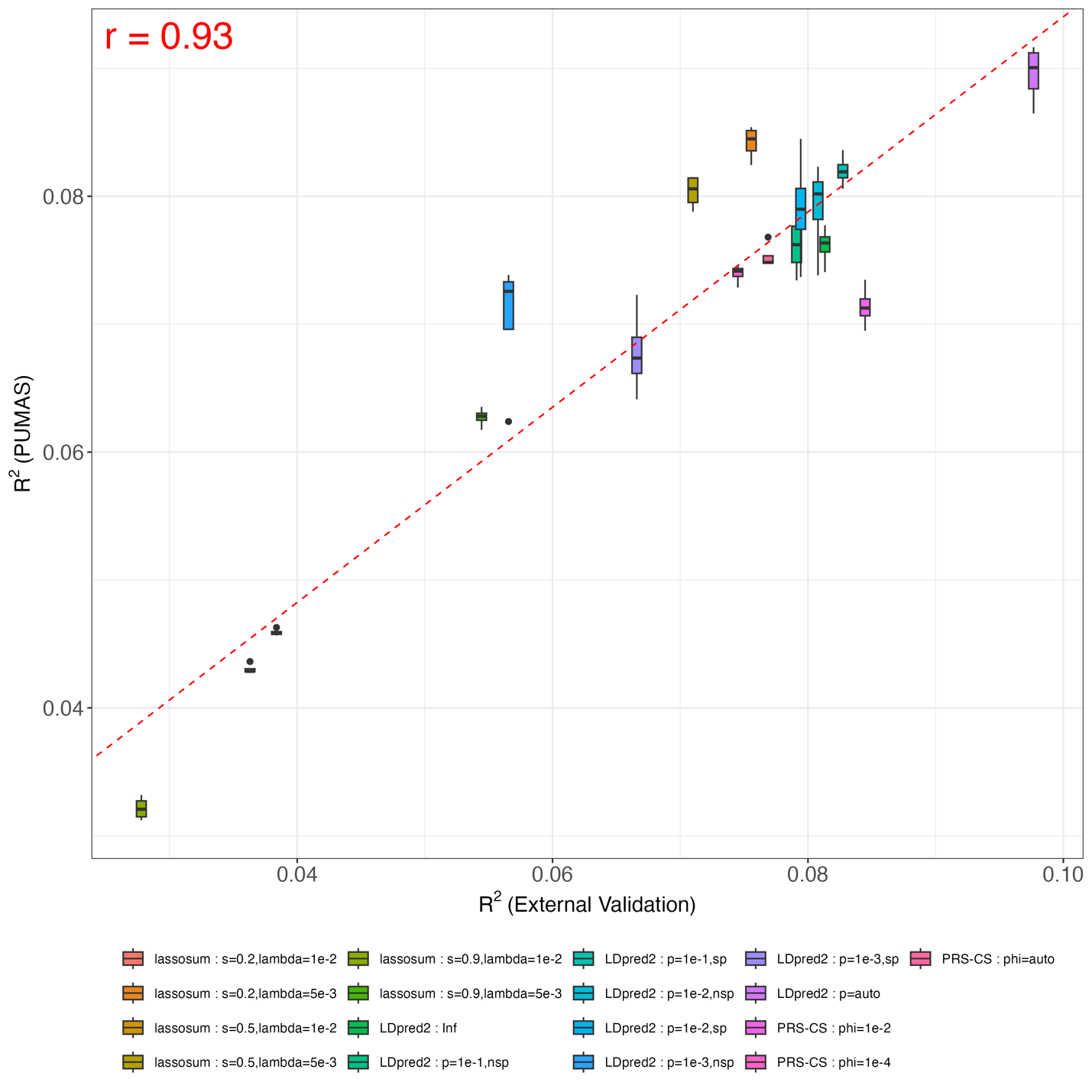

**Fig S27. Comparing PUMAS result for HC with external validation in UKB**. Y-axis: predictive $R^{2}$ across 4-fold replications from PUMAS; X-axis: predictive $R^{2}$ evaluated by external validation on the holdout dataset. The dashed red line is fitted regression line between averaged PUMAS $R^{2}$ and PRS $R^{2}$. Pearson correlation between averaged PUMAS $R^{2}$ and PRS $R^{2}$ is shown in the plot.

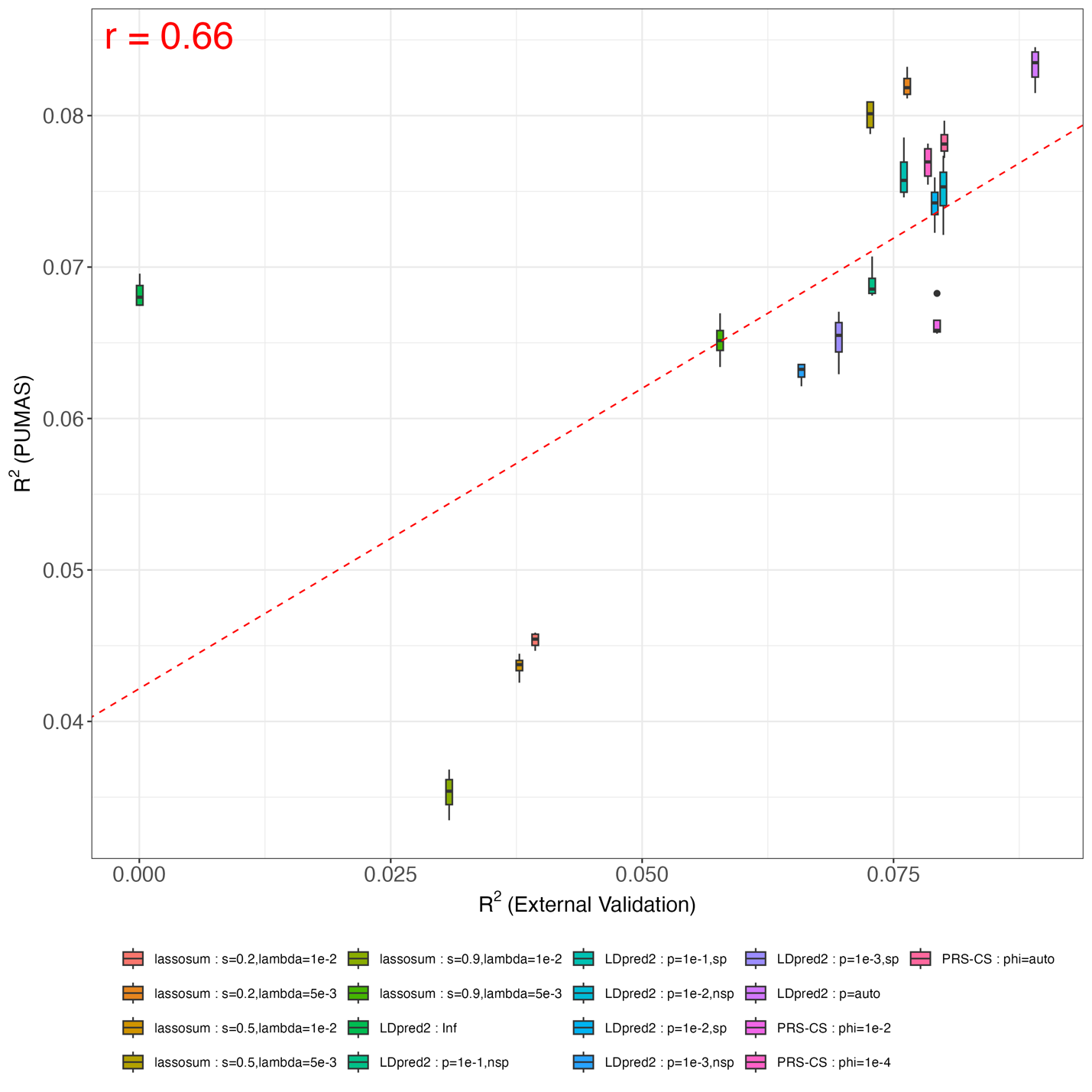

**Fig S28. Comparing PUMAS result for HP with external validation in UKB**. Y-axis: predictive $R^{2}$ across 4-fold replications from PUMAS; X-axis: predictive $R^{2}$ evaluated by external validation on the holdout dataset. The dashed red line is fitted regression line between averaged PUMAS $R^{2}$ and PRS $R^{2}$. Pearson correlation between averaged PUMAS $R^{2}$ and PRS $R^{2}$ is shown in the plot.

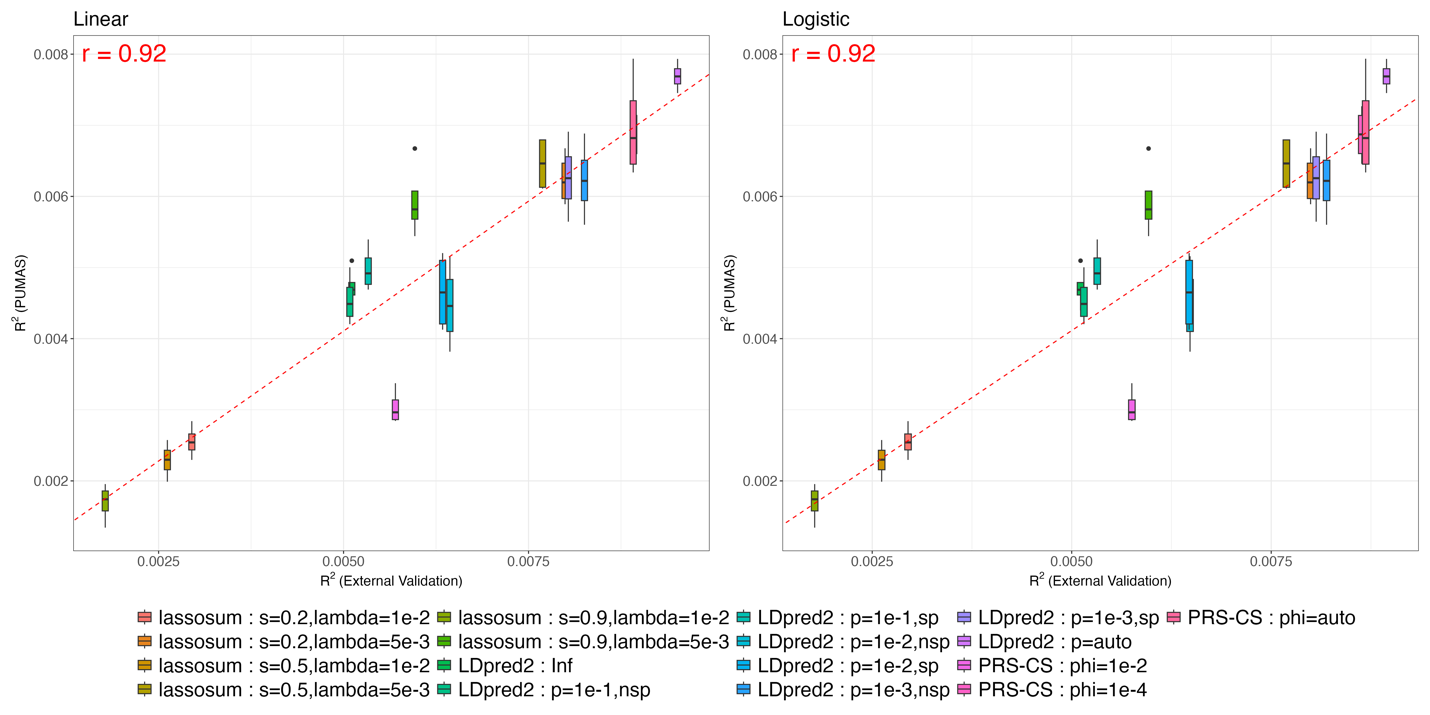

**Fig S29. Comparing PUMAS result for CAD with external validation in UKB**. Y-axis: predictive $R^{2}$ across 4-fold replications from PUMAS; X-axis (left): $R^{2}$ evaluated by external validation on the holdout dataset using transformed summary statistics on the linear scale; X-axis (right): predictive $R^{2}$ evaluated by external validation on the holdout dataset using original summary statistics on the logistic scale. The dashed red line is fitted regression line between averaged PUMAS $R^{2}$ and PRS $R^{2}$. Pearson correlation between averaged PUMAS $R^{2}$ and PRS $R^{2}$ is shown in each panel.

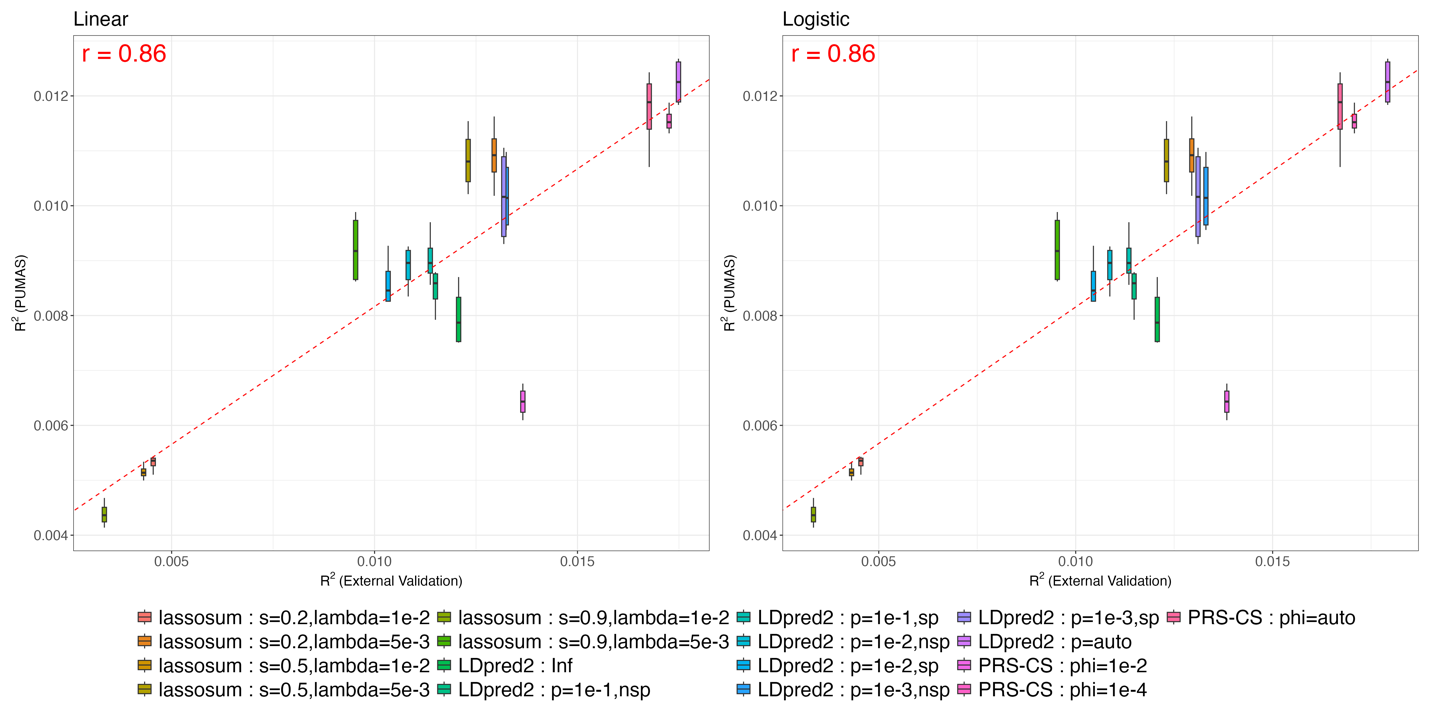

**Fig S30. Comparing PUMAS result for DBT with external validation in UKB**. Y-axis: predictive $R^{2}$ across 4-fold replications from PUMAS; X-axis (left): $R^{2}$ evaluated by external validation on the holdout dataset using transformed summary statistics on the linear scale; X-axis (right): predictive $R^{2}$ evaluated by external validation on the holdout dataset using original summary statistics on the logistic scale. The dashed red line is fitted regression line between averaged PUMAS $R^{2}$ and PRS $R^{2}$. Pearson correlation between averaged PUMAS $R^{2}$ and PRS $R^{2}$ is shown in each panel.

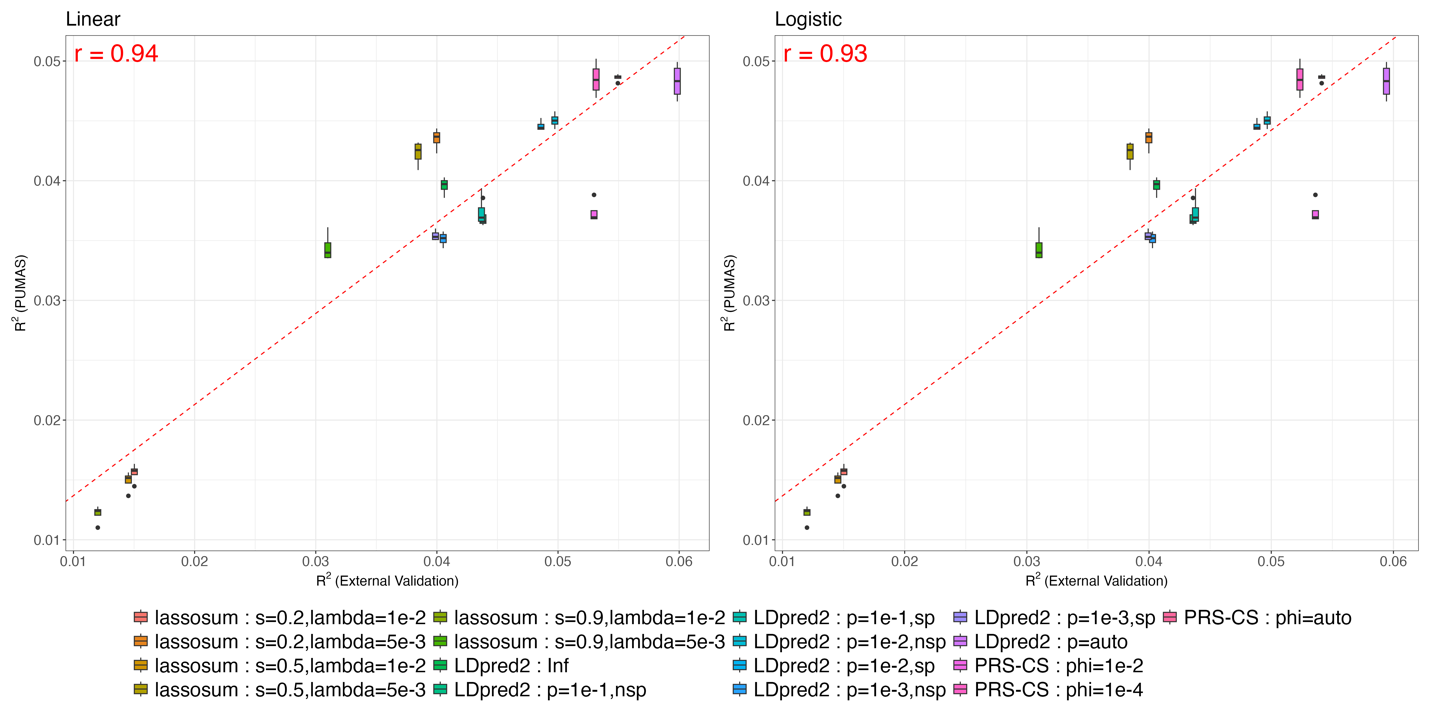

**Fig S31. Comparing PUMAS result for HBP with external validation in UKB**. Y-axis: predictive $R^{2}$ across 4-fold replications from PUMAS; X-axis (left): $R^{2}$ evaluated by external validation on the holdout dataset using transformed summary statistics on the linear scale; X-axis (right): predictive $R^{2}$ evaluated by external validation on the holdout dataset using original summary statistics on the logistic scale. The dashed red line is fitted regression line between averaged PUMAS $R^{2}$ and PRS $R^{2}$. Pearson correlation between averaged PUMAS $R^{2}$ and PRS $R^{2}$ is shown in each panel.

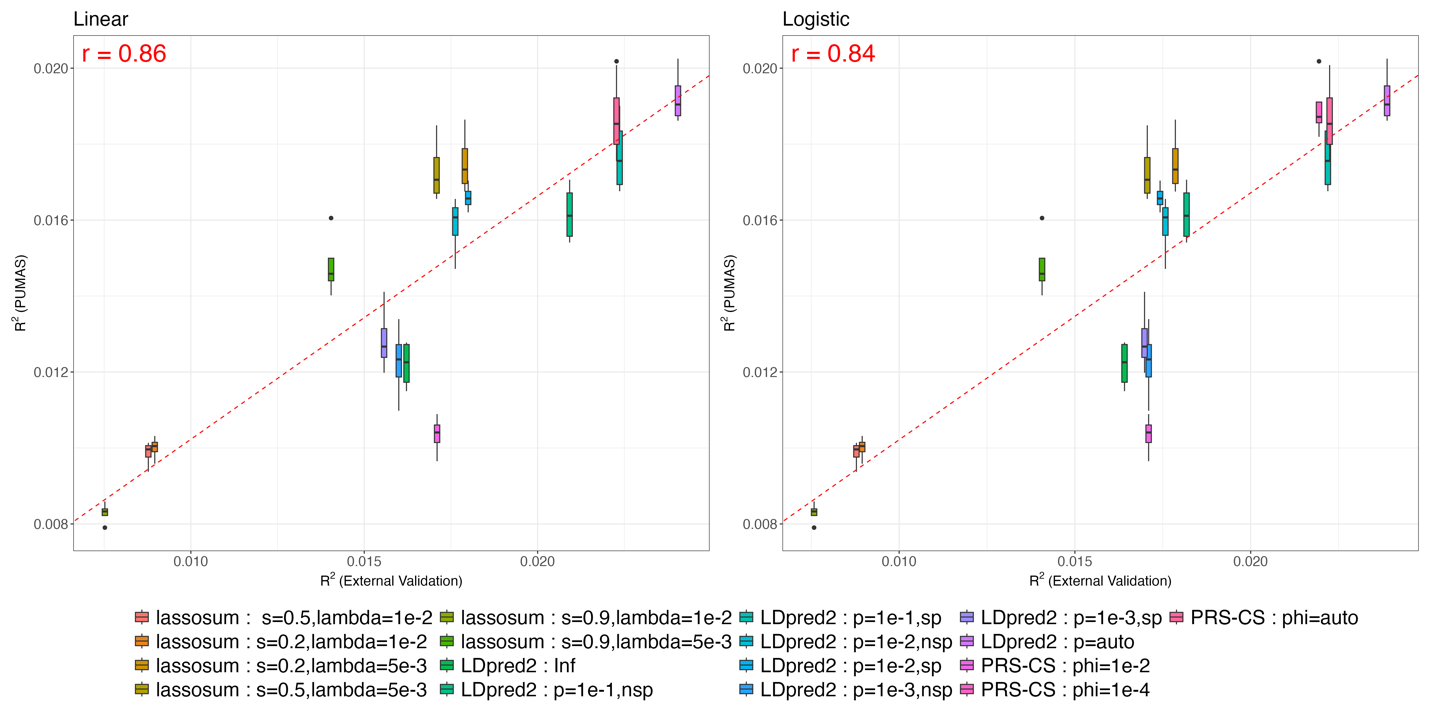

**Fig S32. Comparing PUMAS result for ATH with external validation in UKB**. Y-axis: predictive $R^{2}$ across 4-fold replications from PUMAS; X-axis (left): $R^{2}$ evaluated by external validation on the holdout dataset using transformed summary statistics on the linear scale; X-axis (right): predictive $R^{2}$ evaluated by external validation on the holdout dataset using original summary statistics on the logistic scale. The dashed red line is fitted regression line between averaged PUMAS $R^{2}$ and PRS $R^{2}$. Pearson correlation between averaged PUMAS $R^{2}$ and PRS $R^{2}$ is shown in each panel.

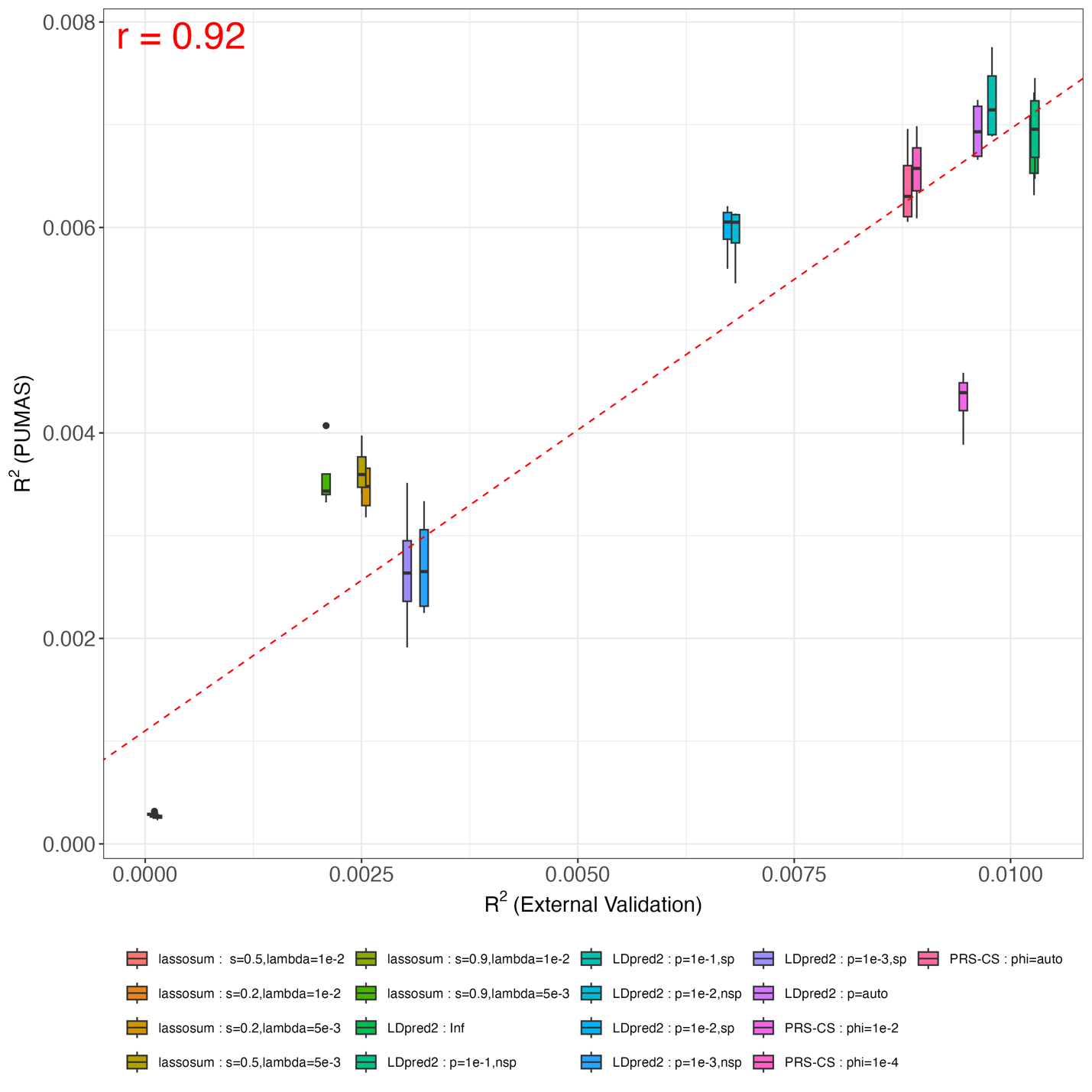

**Fig S33. Comparing PUMAS result for DEP with external validation in UKB**. Y-axis: predictive $R^{2}$ across 4-fold replications from PUMAS; X-axis: predictive $R^{2}$ evaluated by external validation on the holdout dataset. The dashed red line is fitted regression line between averaged PUMAS $R^{2}$ and PRS $R^{2}$. Pearson correlation between averaged PUMAS $R^{2}$ and PRS $R^{2}$ is shown in the plot.

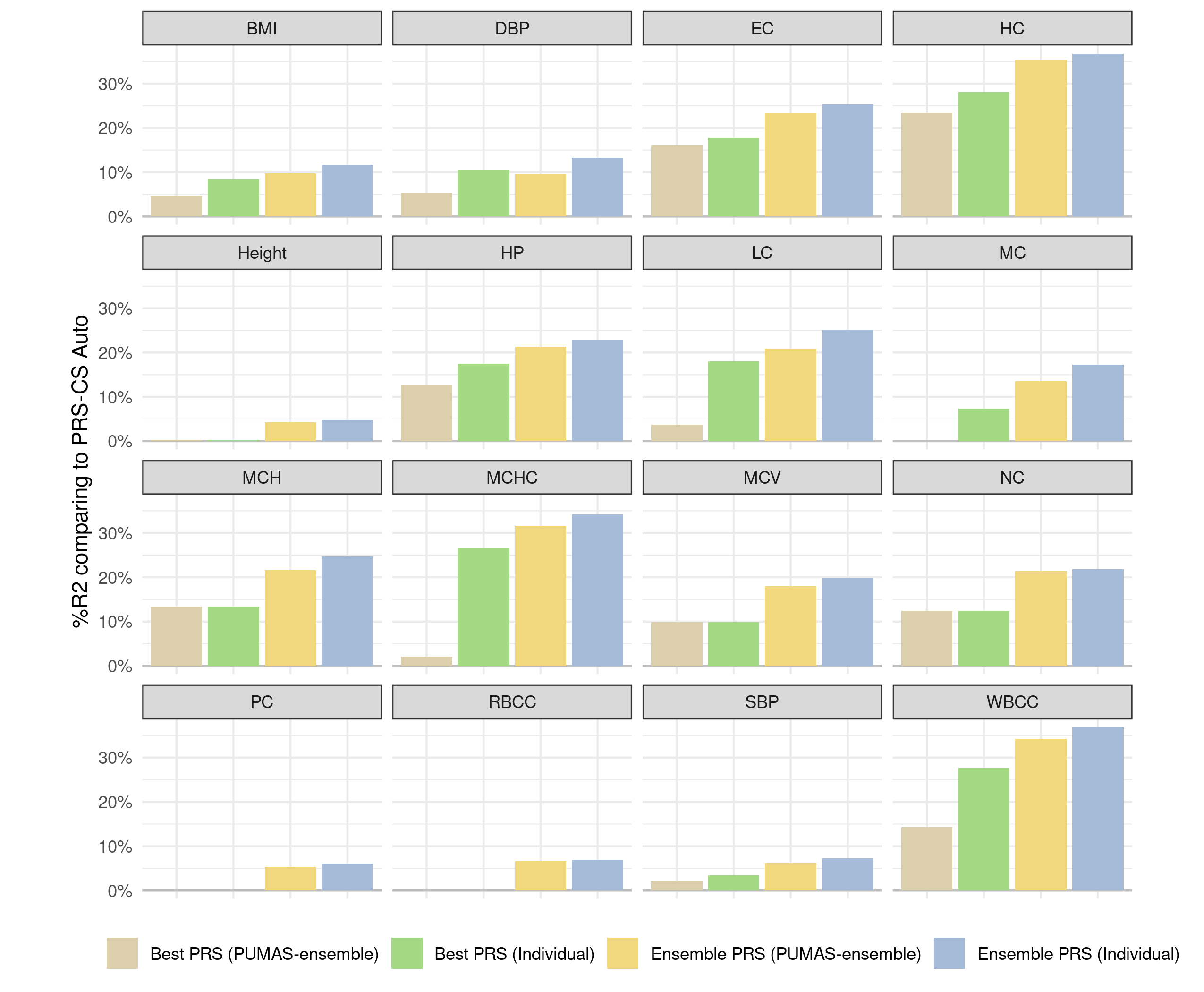

**Fig S34. Prediction gains of ensemble PRS in UKB analysis**. Y-axis: relative percentage increase in $R^{2}$ compared to PRS-CS-auto; X-axis: 4 sets of PRS models, including the best single PRS suggested by PUMAS-ensemble, the best single PRS selected based on the first individua-level holdout set, the ensemble PRS obtained from PUMAS-ensemble, and the ensemble PRS trained from individual-level data. All $R^{2}$ values were computed using the second half of holdout dataset.

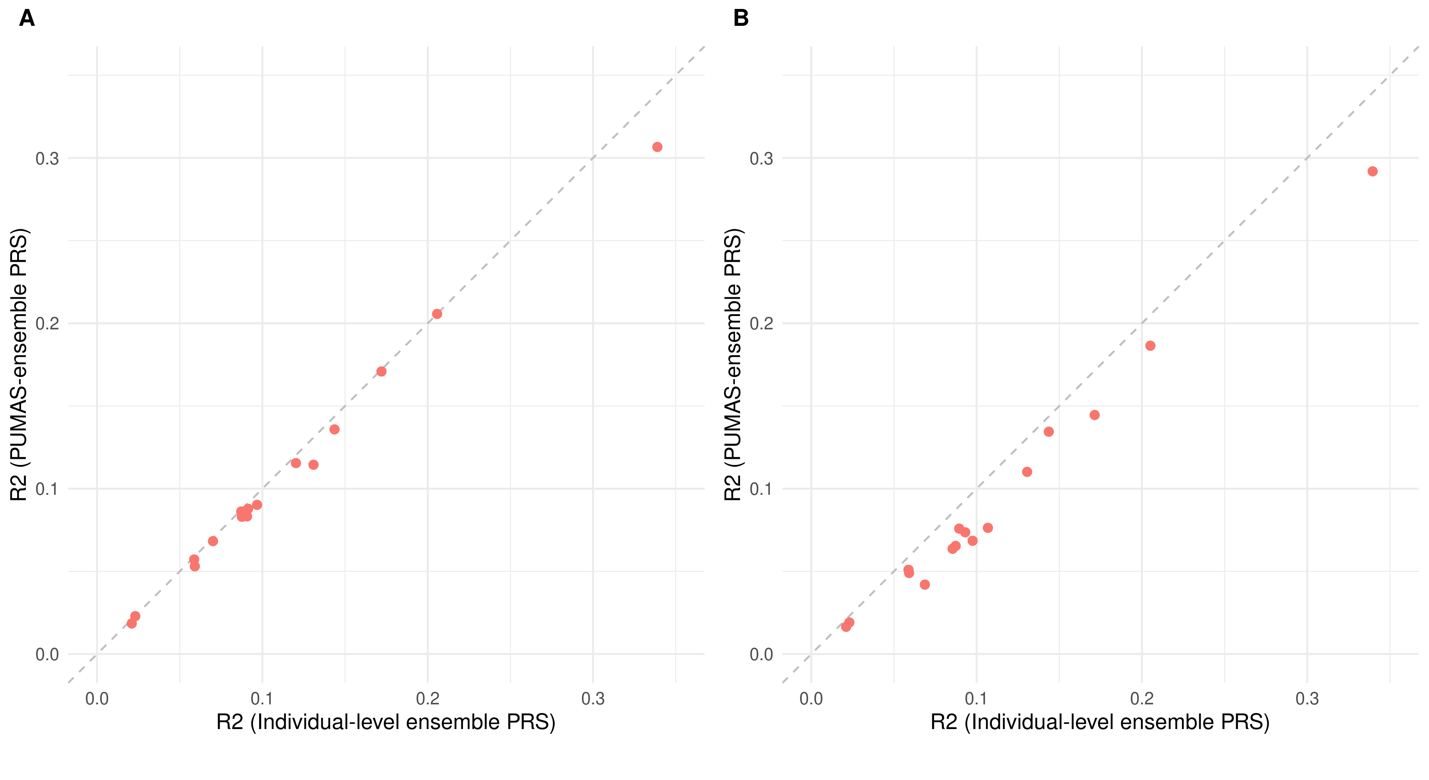

**Fig S35**. **Compare ensemble scores from PUMAS-ensemble and individual-level ensemble approach in UKB using UKB GWAS summary statistics**. PUMAS-ensemble used 1000 Genome Project EUR LD reference data in **A**, and 1000 Genome Project EAS LD reference data in **B**. Y-axis: predictive $R^{2}$ of ensemble scores trained by PUMAS-ensemble and evaluated on the second half of UKB holdout data. X-axis: predictive $R^{2}$ of ensemble scores trained on the first half of individual-level UKB holdout data and evaluated on the second half. The gray dashed line is the diagonal line.

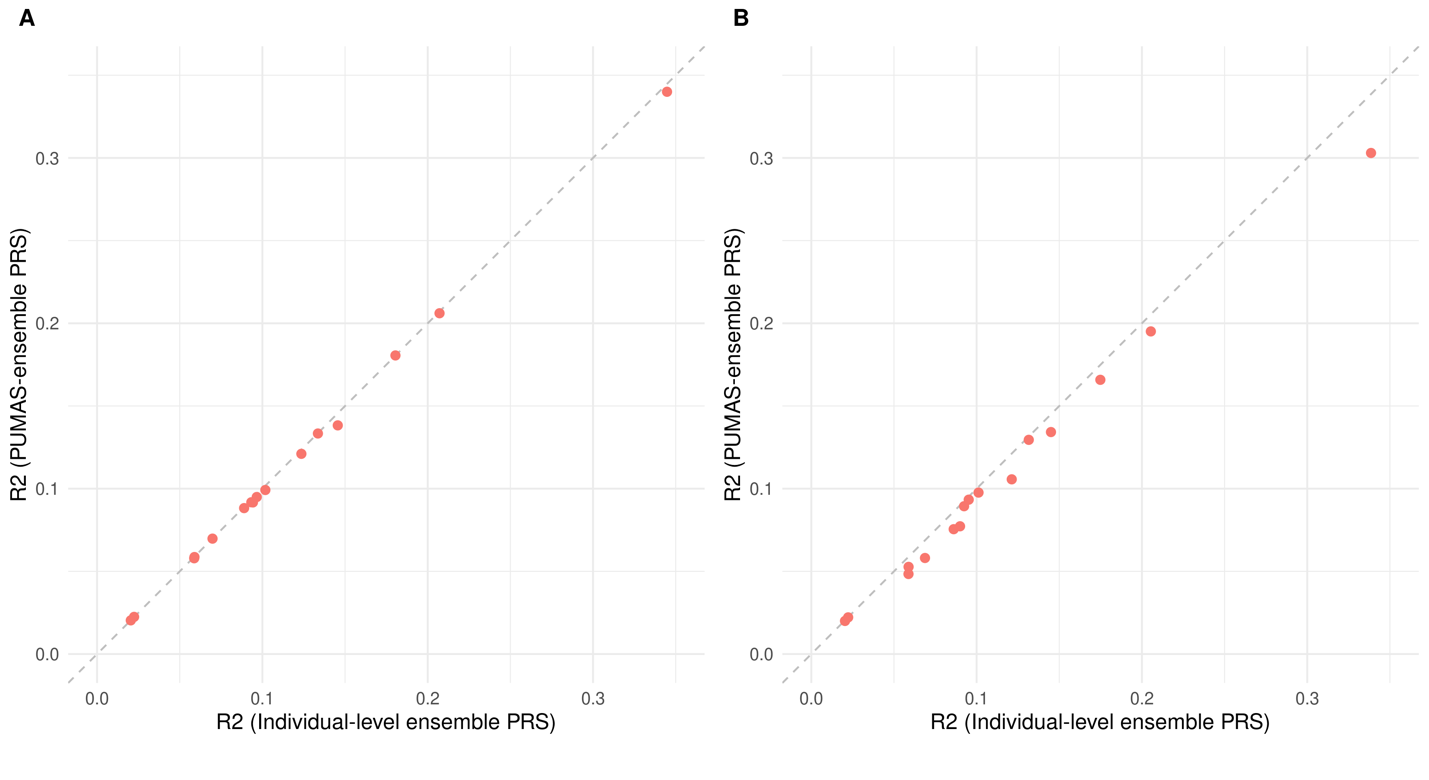

**Fig S36**. **Compare ensemble scores from PUMAS-ensemble and individual-level ensemble approach in UKB using meta-analytic GWAS summary statistics from UKB and BBJ**. PUMAS-ensemble used 1000 Genome Project EUR LD reference data in **A**, and 1000 Genome Project EAS LD reference data in **B**. Y-axis: predictive $R^{2}$ of ensemble scores trained by PUMAS-ensemble and evaluated on the second half of UKB holdout data. X-axis: predictive $R^{2}$ of ensemble scores trained on the first half of individual-level UKB holdout data and evaluated on the second half. The gray dashed line is the diagonal line.

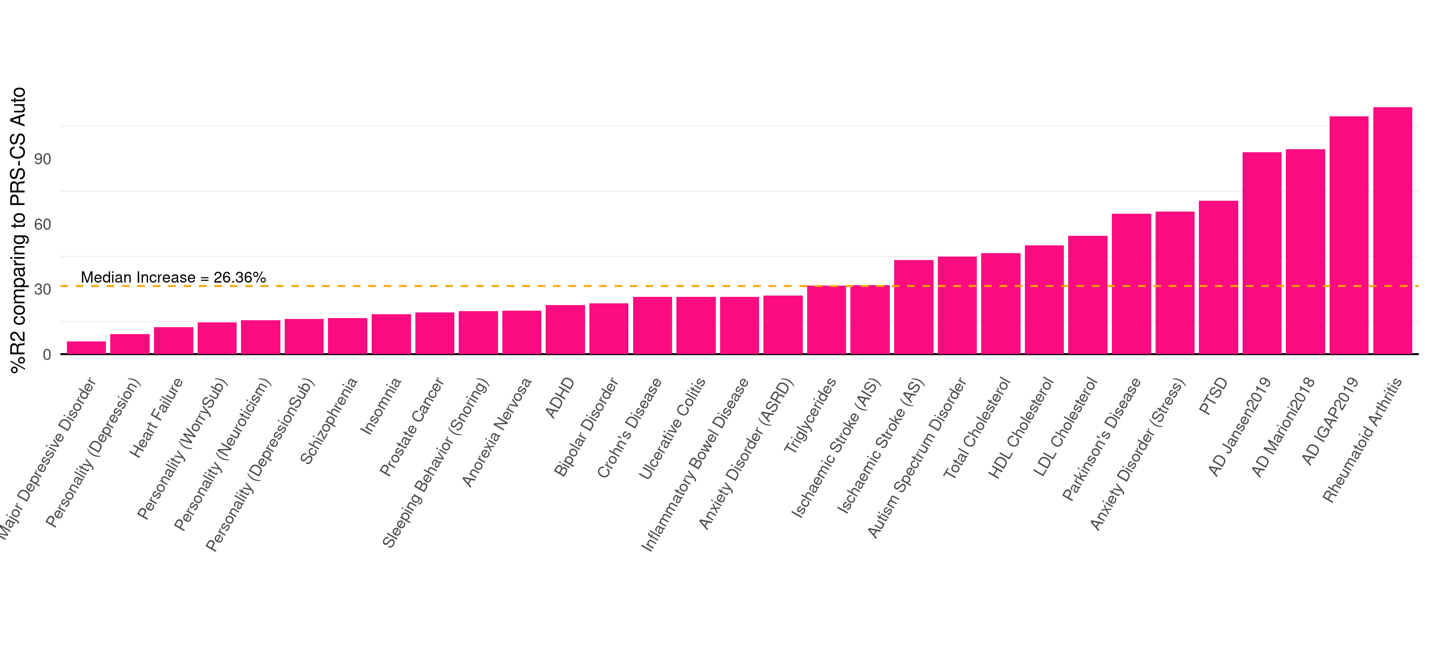

**Fig S37. Prediction gains of ensemble PRS for 31 complex traits**. Y-axis: relative percentage increase in $R^{2}$ compared to PRS-CS-auto; X-axis: trait names. Three Alzheimer’s disease GWAS summary statistics are labeled by study name and time.

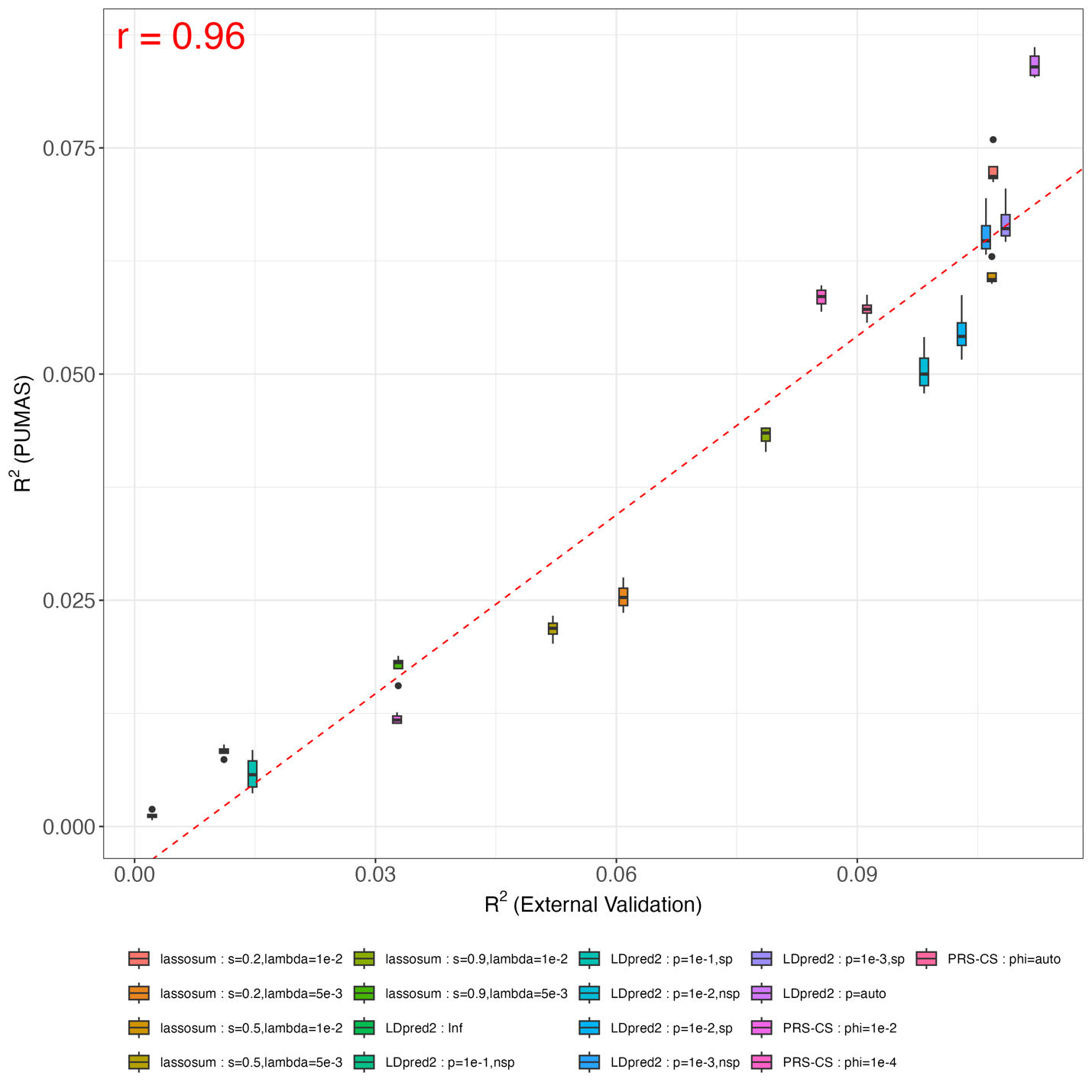

**Fig S38. Comparing PUMAS result for IGAP 2019 AD GWAS with external validation in UKB**. Y-axis: predictive $R^{2}$ across 4-fold replications from PUMAS; X-axis: predictive $R^{2}$ evaluated by external validation on the UKB holdout dataset with 2,600 AD cases and 5,200 healthy controls. The dashed red line is fitted regression line between averaged PUMAS $R^{2}$ and PRS $R^{2}$. Pearson correlation between averaged PUMAS $R^{2}$ and PRS $R^{2}$ is shown in the plot.
